# Hidden molecular relationships are revealed by bootstrap resampling of mass spectral pairs with SpecReBoot

**DOI:** 10.64898/2026.02.03.703446

**Authors:** Esteban Charria-Girón, Laura Rosina Torres-Ortega, Joelle Mergola Greef, Yasmina Marin Felix, Nelson H. Caicedo, Frank Surup, Marnix H. Medema, Justin J.J. van der Hooft

## Abstract

Mass spectral molecular networking organizes tandem mass spectrometry (MS/MS) data by connecting spectra based on similarity scores. However, these deterministic metrics provide no measure of uncertainty, so molecular networks often retain edges arising from noise or missing fragments, while authentic chemical relationships may remain obscured. To address this gap, we present SpecReBoot, a statistical framework that adapts Felsenstein’s bootstrap principle to MS/MS similarity scoring. By resampling fragment-level features and recomputing similarity across replicates, SpecReBoot transforms a single score into a confidence distribution and assigns bootstrap support to network edges. Using large, curated MS/MS spectral datasets, we demonstrate that SpecReBoot systematically removes unreliable connections and recovers robust low-similarity links, refining networks and revealing hidden relationships. We highlight how this confidence-aware “rebooting” guided the discovery of an unprecedented macrolactone scaffold from the endophytic fungus *Diaporthe caliensis*. Altogether, SpecReBoot provides the first general framework for quantifying confidence in MS/MS similarity and molecular network analysis.

---

Small molecules are studied across the life and environmental sciences, with applications ranging from mapping metabolic diversity in living organisms to monitoring environmental contaminants, characterizing drug candidates, and ensuring food safety. Tandem mass spectrometry (MS/MS) coupled to liquid chromatography (LC) has emerged as the central analytical platform enabling this at scale, offering high sensitivity and providing structural insights in complex biological samples.^1,2,3^ The volume and complexity of MS/MS data generated in these experiments has driven the development of computational approaches to organize and contextualize spectral information. Among these, molecular networking (MN) has become a widely adopted strategy that groups structurally related molecules by connecting spectra based on spectral similarity.^4,5^

The introduction of MN through the GNPS infrastructure has democratized the analysis of MS/MS spectra, enabling large-scale organization, visualization, and annotation of spectral relationships across fields.^4,6^ In these networks, MS/MS spectra are represented as nodes connected by similarity-based edges, allowing researchers to efficiently explore chemical space and prioritize unknown metabolites for downstream investigation^7,8^. Several spectral similarity scores have been introduced and can broadly be classified in: cosine-based scores that compare aligned fragment ion intensities, and machine learning–based approaches that embed spectra into a latent space to capture higher-order fragment context such as Spec2Vec, MS2DeepScore, and DreaMS.^9–13^ Across all similarity scores, users must define a similarity threshold to prune weak connections and enable network construction, a choice that – in the inherent absence of ground truth – is often arbitrary and strongly influences the resulting topology and downstream interpretation.

Despite the broad acceptance of MN as a cornerstone method to chart metabolome diversity, the underlying similarity scores on which these networks are built are treated as absolute. These scores remain deterministic and lack any explicit evaluation of the reproducibility of inferred spectral connections under perturbations of the underlying data. Regardless of the score choice, users must rely on fixed thresholds, often without a clear understanding of how individual connections are built. Accordingly, MNs often require careful manual inspection to distinguish true (bio)chemical relationships between connected nodes from spurious or noise-driven connections.^8,14,15^

Indeed, alongside the success stories following the introduction of GNPS, multiple studies have highlighted fundamental challenges inherent to the study of MS/MS data, including spectral noise, incomplete or missing fragment peaks, chimeric spectra, instrument- and acquisition-dependent variability, and the impact of in-source fragmentation.^15–19^ Remarkably, up to this date, no general statistical framework has been proposed and developed to quantify the reproducibility of spectral connections within MNs. This lack of confidence-aware metrics limits the reliability of MN-derived hypotheses and highlights a clear methodological gap.

To address the lack of uncertainty estimation in MN (Fig. 1a), we developed SpecReBoot, a general statistical framework that introduces bootstrap-based confidence measures into mass spectral similarity analysis. SpecReBoot draws inspiration from phylogenetic bootstrapping by treating fragment peaks as resampling units (Fig. 1b), generating pseudo-replicate spectra from which spectral similarities are recalculated. This procedure yields two complementary quantities for each spectral pair, a mean similarity, and an edge support score that reflects the stability of their mutual nearest-neighbor relationship across bootstrap replicates (Fig. 1c).

**Fig. 1.**
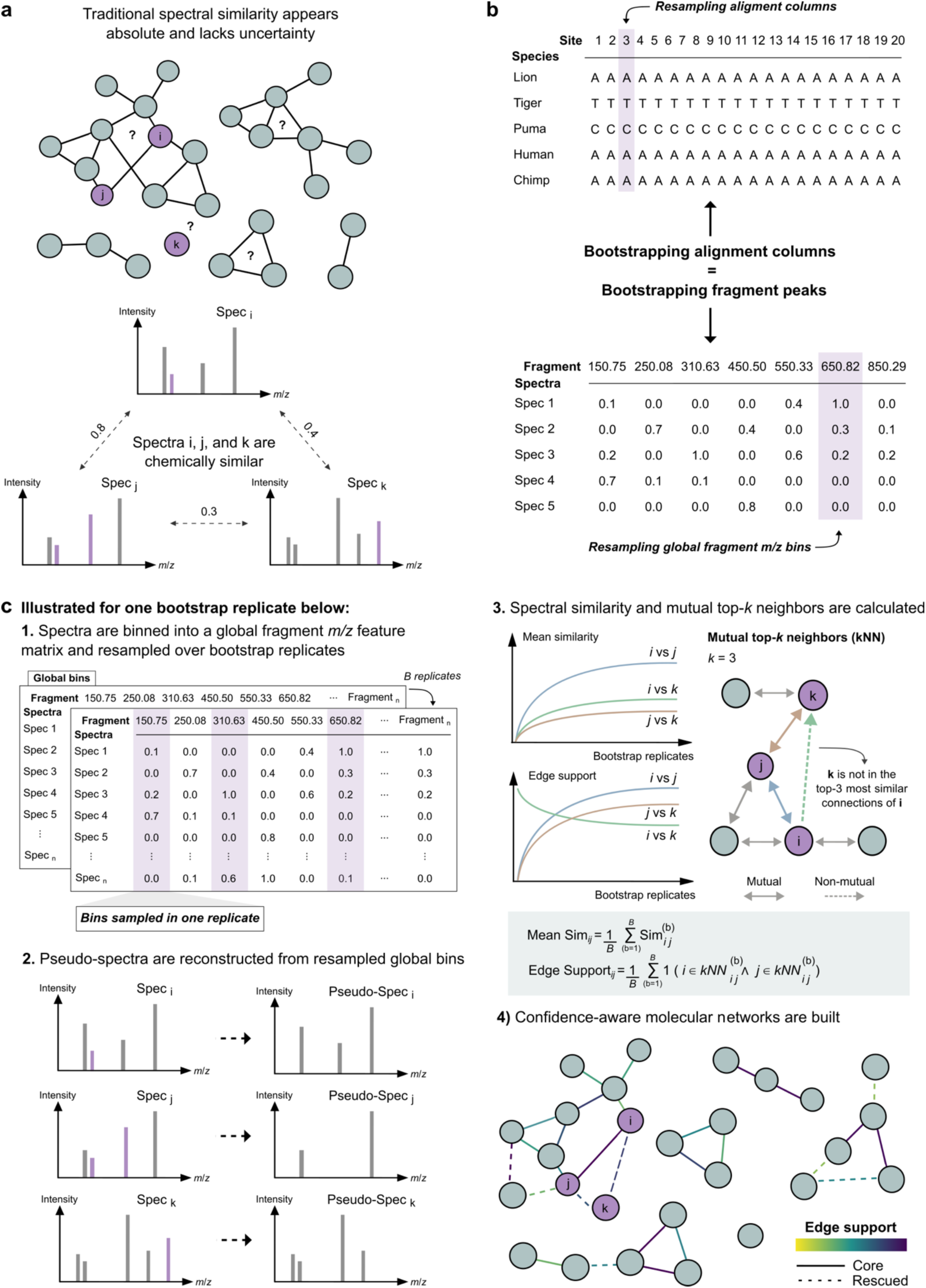
Bootstrapping fragment peaks introduces confidence-aware spectral similarity in metabolomics. **a,** in conventional molecular networking, MS/MS spectra are represented as nodes connected by edges constructed from pairwise similarity scores. These values are typically treated as absolute, even though they lack any estimate of stability under perturbations of the underlying spectra. **b**, SpecReBoot adapts nonparametric bootstrapping to MS/MS data. In analogy to phylogenetics, where alignment columns are resampled across replicates, fragment peaks are first binned into a global fragment *m*/*z* features matrix that includes all observed fragments across spectra. In each bootstrap replicate, fragment *m/z* bins are resampled with replacement (on average ∼63% of global bins are sampled per replicate), and pseudo-replicate spectra are reconstructed from resampled bins, altering which peaks contribute to similarity calculations. **c**, the workflow is illustrated for a single bootstrap replicate. For each replicate, spectral similarities are recalculated and mutual top-*k* nearest neighbors are identified, meaning that edge support increases only when, for a given spectral pair, each spectrum appears within the other’s top-*k* most similar spectra, as in the pairs i–j and j–k, while k is not in the top 3 of i. Repeating this process over *B* replicates produces two complementary measures: (i) a mean similarity, defined as the average spectral similarity across replicates, and (ii) an edge support, defined as the frequency with which two spectra remain mutual nearest neighbors. These confidence-aware measures are then used to “reboot” molecular networks, highlighting stable connections (core), filtering unstable edges, and rescuing previously missed relationships (rescued), enabling more reliable and interpretable MS/MS-based metabolomics analyses.

In practice, these measures allow MNs to be interpreted not solely based on similarity scores, but also on the robustness of spectral connections. Together, these quantities enable the construction of confidence-aware MNs, either by retaining only edges with both high similarity and high support, or by rescuing edges with low similarity but consistently high support across replicates (Fig. 1d). Hence, SpecReBoot improves the recovery of chemically meaningful spectral connections across different similarity scores, including cases in which chemical links were missed by just using similarity scores alone. By recording the fragment bins sampled in each bootstrap replicate, changes in edge support values can be linked to fragment-level information revealing key fragments that support reliable spectral connections, thus enhancing network interpretability and providing an explainable layer for recovered spectral relationships, including those derived from machine learning-based scores, ultimately improving metabolomics-based discovery workflows.

## Results

### Mean similarity and edge support reveal stable and chemically meaningful spectral relationships across metrics

To introduce SpecReBoot and illustrate the concept of bootstrap-derived confidence estimation applied to mass spectral relationships, we applied it to a curated subset of MS/MS spectra from ribosomally synthesized and post-translationally modified peptides (RiPPs). Whilst structures from this biosynthetic class have been known for decades, their actual biosynthesis RiPP pathways having been discovered relatively recently^20,21^, they are now known to be widely spread in nature^22,23^. Even with increased research into RiPPs^24^, the availability of MS/MS reference spectra remains limited. RiPPs exhibit extensive chemical diversity, as they may contain any canonical amino acid sequence, vary in length, and undergo diverse post-translational modifications. Ongoing discovery of new RiPP families further highlights their diversity^24^, while the presence of heterogeneous modifications often leads to complex and unpredictable fragmentation patterns. These properties make them a suitable test case for evaluating the robustness of inferred spectral relationships.

RiPP-associated MS/MS spectra were retrieved from annotated metabolite features from public GNPS repositories through manual curation of spectra classified as peptides by NPClassifier^25^. Spectra were cross validated against original reports to ensure data integrity, spectral quality, and consistency with a RiPP origin. This resulted in a curated set of 24 RiPP-associated spectra, from which, we selected the aerucyclamides as a representative case study. These are RiPPs first reported from the cyanobacterium *Microcystis aeruginosa* as hexacyclopeptides^26,27^. Their biosynthesis was later reassigned to a ribosomal pathway by comparative analyses to the patellamide pathway characterized in cyanobacterial symbionts^28^. Despite their high mutual structural similarity, these molecules exhibit near-zero modified cosine spectral similarity. Specifically, similarity values of 0.0004 (A–B), 0.0000 (A–C), and 0.0210 (B–C) are obtained, causing these relationships to be entirely missed when relying solely on spectral similarity thresholds.

Using SpecReBoot, RiPP-associated spectra were represented as global *m/z* bins, after which fragment bins were resampled with replacement across bootstrap replicates to generate pseudo-spectra. The general procedure is illustrated for the aerucyclamides in Fig. 2a. For each replicate, pairwise similarities were recalculated and mutual top-*k* nearest neighbors were identified, where *k* defines the number of nearest neighbors considered per spectrum (a user-defined parameter; see Methods). Repeating this process across *B* bootstrap replicates results in two complementary quantities for each spectral pair, (i) a mean similarity, defined as the average similarity across replicates, and (ii) the edge support, reflecting the frequency with which two spectra are recovered as mutual kNN neighbors, meaning that in a given replicate each spectrum is ranked among the other’s top-*k* most similar spectra. For example, if after 100 bootstrap replicates spectra *i* and *j* are recovered as mutual top-*k* neighbors 80 times, the resulting edge support will be 0.8.

**Fig. 2.**
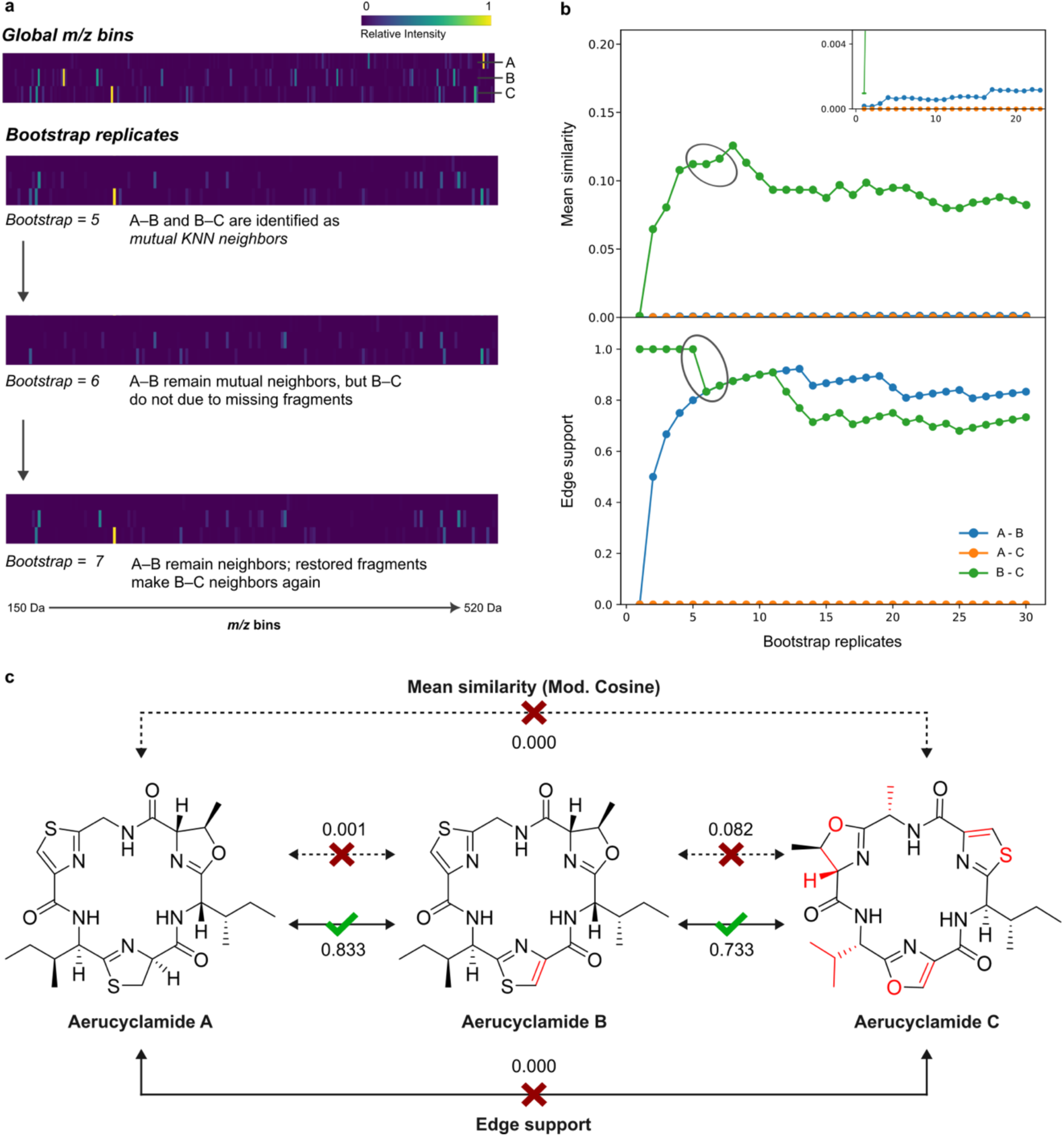
Fragment resampling uncovers confidence-aware spectral relationships in aerucyclamides. **a,** schematic illustration of bootstrap resampling for aerucyclamides A–C. MS/MS spectra are first represented as global *m/z* bins, after which bins are resampled with replacement to generate pseudo-replicate spectra. Example bootstrap replicates (5–7) show how changes in fragment composition affect *k*-nearest-neighbor (KNN) relationships between spectra. Across replicates, aerucyclamides A and B are consistently identified as mutual neighbors, whereas the relationship between B and C is variably present depending on the resampled fragment set. **b,** trajectories of mean spectral similarity (top) and edge support (bottom) across increasing numbers of bootstrap replicates for all pairwise comparisons among aerucyclamides A–C. Values are shown as a function of the number of bootstrap replicates. The inset on the top right corner highlights the low-amplitude variation in mean similarity for the A–B pair at an expanded scale. **c,** comparison of mean similarity (dashed lines) and edge support (solid lines) values for aerucyclamides A–C using the modified cosine. While the mean similarity results in a near-zero (≤ 0.01) score for these structurally related compounds, SpecReBoot recovers stable relationships between A–B and B–C based on high edge support. Structural differences between aerucyclamides are highlighted in red.

For aerucyclamides A–C, these measures revealed distinct stability patterns (Fig. 2b). The A–B pair, which exhibits near-zero similarity, showed a slight increase in the mean similarity across replicates (see inset in Fig. 2b), indicating that similarity emerges only under specific fragment subsets. In contrast, the A–C pair consistently showed both mean similarity and edge support equal to zero. Analysis of the edge support further demonstrated that the A–B and B–C pairs were recovered across replicates as mutual nearest neighbors, whereas the A–C relationship remained unsupported. These patterns reflect the stability of spectral relationships under fragment-level perturbation rather than the magnitude of chemical similarity observed. As the number of replicates increased (*B* = 30), both mean similarity and edge support converged, providing stable and interpretable measures of the robustness of spectral connections.

Because SpecReBoot records sampled *m/z* bins across replicates, it enables direct correlation of changes in mean similarity and edge support with fragmentation patterns. Informative fragments can be mapped back onto reference spectra, providing a mechanistic explanation of why particular relationships are stabilized or lost under resampling. For aerucyclamides (Fig. S1–S3), *m/z* bins repeatedly associated with gains or losses of mutual kNN relationships correspond to conserved substructures. For instance, fragment ions at *m/z* 425.2 and 436.2 Da could be traced back to differences in the amino acid composition between aerucyclamides A and B. The fragment at *m/z* 425.2 corresponded to a motif uniquely present in aerucyclamide A, comprising thiazoline, *L*-Ile, thiazole, and *D*-allo-Ile residues. In contrast, the fragment at *m/z* 436.2, observed exclusively in aerucyclamide B, arises from a closely related substructure containing *L*-Ile, thiazoline, *D*-allo-Ile, and an oxazoline ring (Fig. S3).

MS2DeepScore captures connections between aerucyclamides with mean similarities of 0.676 (A–B), 0.769 (A–C), and 0.571 (B–C), all below the suggested 0.85 threshold^13^. Although MS2DeepScore and modified cosine rely on distinct sets of *m/z* bins (Fig. S2b), they show partial overlap in fragment-level evidence. For example, both metrics identify the fragment at *m/z* 332.1 Da as positively correlated with edge support between aerucyclamides A and B. Beyond this shared fragment, however, the supporting fragments diverge. For this pair, positively correlated fragments above *m/z* 475 Da are predominantly even-mass ions for modified cosine, but mainly odd-mass ions for MS2DeepScore. These differences suggest that embedding- and alignment-based approaches can converge on the same stable spectral relationships while relying on chemically distinct fragment subsets.

Together, these results demonstrate how SpecReBoot enhances the explainability of spectral similarity by directly linking fragment-level information to edge support, rationalizing the recovery of reproducible relationships. In the context of this RiPP example, edge support is particularly informative, as it reconnects aerucyclamide spectra into a coherent MF within a small network comprising diverse RiPP families (Fig. S4).

### Rescuing hidden polyketide-lactone diversity in *Diaporthe caliensis* reveals an unprecedented macrolactone scaffold

To further assess the applicability of SpecReBoot for NPs discovery, and in the context of the growing need for new antimicrobials, we re-examined a previously published MN derived from the endophytic fungus *Diaporthe caliensis*. This species, isolated from the medicinal plant *Otoba gracilipes*, is part of our ongoing efforts to explore the biotechnological applications of Colombian fungal diversity^29–32^. Among the metabolites produced by this fungus are the bioactive 10-membered macrolactone phomol and the butanolides caliensolide A and B. In the original study, cosine-based MN clustered phomol and caliensolide B within the same MF, while caliensolide A belonged to a separate cluster^30^. Subsequent application of MS2LDA partially recovered these relationships through shared Mass2Motifs, largely composed of recurring neutral losses, highlighting limitations of similarity-based networking alone and suggesting complex underlying fragmental-level relationships between these molecules^30^.

We therefore applied SpecReBoot to re-examine spectral relationships among these polyketide-lactones, focusing first on caliensolides A and B introduced above. These two analogues sharing a common biosynthetic origin remained disconnected under conventional MN. Bootstrap analysis (*B* = 100) revealed a mean modified cosine similarity of 0.32 and an edge support of 0.22. Although the similarity is below a threshold of 0.7 for the modified cosine score, the non-zero edge support indicates that this relationship is repeatedly recovered across some bootstrap replicates, suggesting that fragment perturbations increase the overall similarity across specific replicates. Motivated by this finding we applied an exploratory edge support threshold of 0.2 to “reboot” network topology and evaluate whether bootstrap-derived confidence improves connectivity of known polyketide-lactones. This produced a markedly different topology compared to the original MN (Fig. 3). By filtering unstable edges and rescuing low-similarity connections with edge support over 0.2, SpecReBoot generated a coherent network in which multiple previously disconnected MFs became linked. Accordingly, caliensolides A and B appeared as neighboring nodes within the same MF as phomol, in agreement with earlier evidence obtained with MS2LDA that discovered shared Mass2Motifs between the mass spectra of these molecules^30^.

**Fig. 3.**
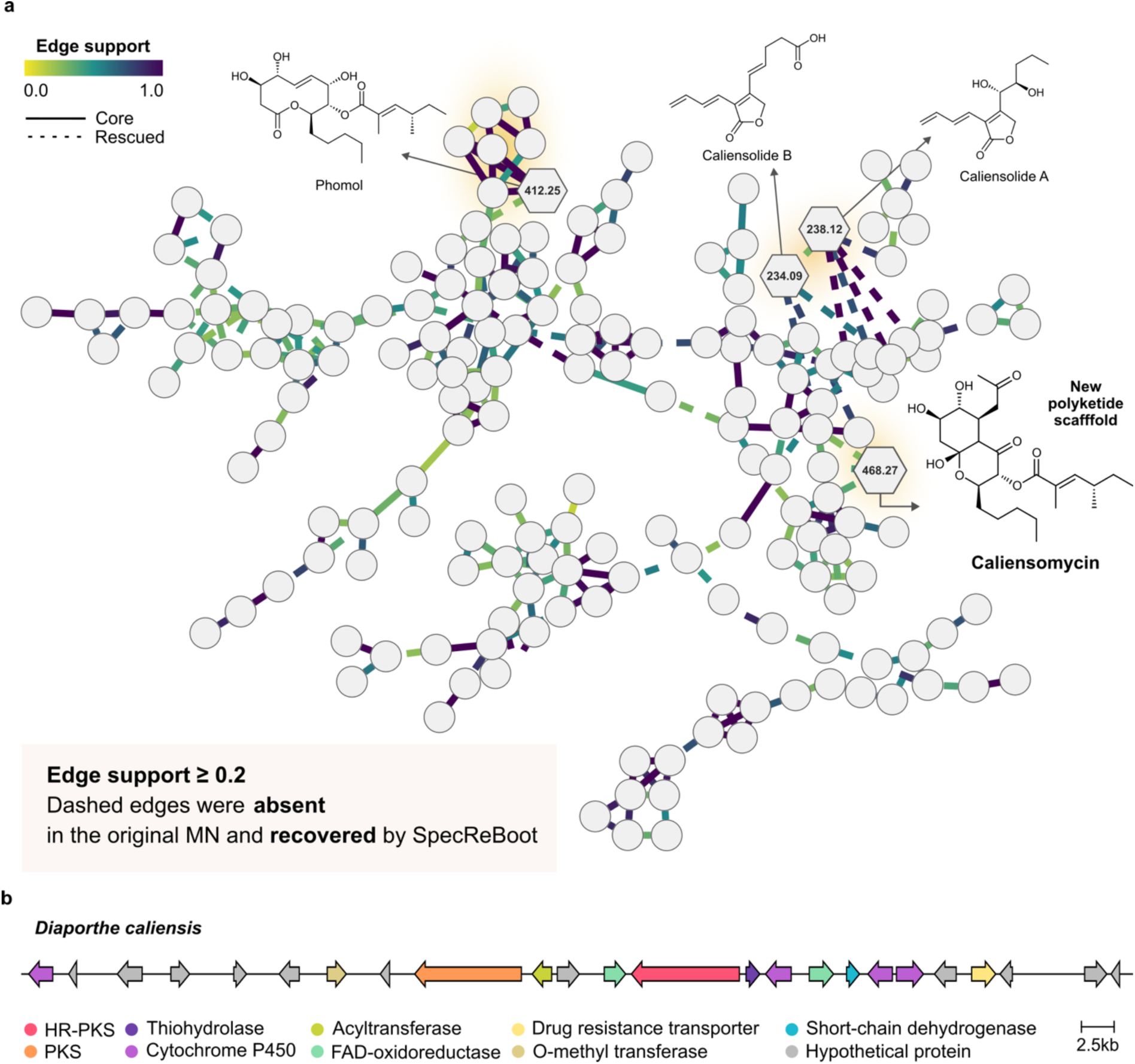
SpecReBoot rescues hidden polyketide-lactone diversity in *Diaporthe caliensis* and enables discovery of a novel macrolactone scaffold. **a**, molecular network reconstructed using bootstrap-derived edge support. Nodes represent MS/MS spectra and edges indicate spectral relationships. Solid edges denote core connections present in the original molecular network, whereas dashed edges correspond to relationships rescued by SpecReBoot that were absent under conventional similarity-based networking. Application of an edge-support threshold reconnects phomol with caliensolide A and B and reveals additional, previously disconnected chemistry. Guided by rescued connections, an uncharacterized node (*m/z* 468.27) was prioritized, leading to the isolation and structural elucidation of caliensomycin, an unprecedented polyketide-lactone scaffold. **b,** candidate biosynthetic gene cluster (BGC) region for the production of phomol and caliensomycin in the genome of *Diaporthe caliensis*.

Encouraged by these observations, we then hypothesized that additional, chemically uncharacterized polyketides associated with this MF had been initially overlooked. Guided by the SpecReBoot “rebooted” network, we reinvestigated fractions obtained after the scaled-up solid cultivation of *D. caliensis* and flash chromatographic pre-fractionation. Within this MF, comprising 179 features connected through many recovered edges, we detected a previously uncharacterized mass feature with a molecular weight of 468 Da and molecular formula of C_24_H_38_O_8_ in a fraction distinct from the one yielding phomol and caliensolides. This feature differed from phomol by an additional C_3_H_2_O unit, suggesting a related carbon skeleton and supporting its annotation as a putative polyketide-lactone. The corresponding structure was purified by preparative HPLC and elucidated by HRESI-MS and comprehensive 1D- and 2D-NMR spectroscopic analyses (Table S1, Fig. S5–10). This metabolite, representing an unprecedented polyketide macrolactone scaffold, was named caliensomycin, honoring Cali, where its fungal producer was originally isolated.

The close structural resemblance between phomol and caliensomycin, together with their shared relative and absolute configurations, suggested a common biosynthetic origin. We therefore formulated the hypothesis that both compounds are produced by the same or closely related biosynthetic gene clusters (BGCs), in which a highly reducing polyketide synthase (HR-PKS) assembles the macrolactone scaffold and a second PKS contributes the aliphatic side chain esterified at C11. This motivated us to investigate the biosynthetic potential of *D. caliensis*, prompting genome sequencing and systematic analysis of its predicted BGC repertoire. Among 90 BGC regions predicted by antiSMASH 8.0.4^33^, five candidate loci encoded two PKSs within the same cluster. Comparative synteny analysis against known macrolactone-producing clusters (Fig. S11), including those for aspinolide and brefeldin A, led to identification of a candidate with high similarity (Fig. 3b), supporting our hypothesis. In particular, the HR-PKS showed high sequence identity across all clusters, consistent with a conserved role in macrolactone formation, whereas the second PKS was shared only between the aspinolide and the phomol clusters, in agreement with O-substitutions at the macrolactone ring^34,35^.

The candidate BGC in *D. caliensis* is larger than the aspinolide and brefeldin A BGCs and encodes an expanded tailoring machinery, including multiple cytochrome P450 monooxygenases and oxidoreductases, providing a plausible basis for the late-stage oxidative rearrangements observed in caliensomycin (Fig. S12). This scaffold reorganization is consistent with oxidative transformations previously described for brefeldin A and SCH metabolites^34,36^. Together, these findings demonstrate that SpecReBoot not only improves confidence in MN but can directly facilitate biological discovery by prioritizing robust yet low-similarity spectral relationships that guided compound isolation and biosynthetic hypothesis generation.

### SpecReBoot provides scalable and interpretable spectral similarity across large-scale MS/MS mass spectral library datasets

We next used publicly available, curated MS/MS spectral datasets of different size and chemical diversity to evaluate SpecReBoot‘s scalability and how SpecReBoot impacts molecular networking using structural and chemical compound class term consistency of the formed molecular families. Specifically, we analyzed three representative spectral collections: a pesticide dataset previously introduced in the *matchms* framework^37^, the NIH Natural Products Library, comprising approximately 1,200 curated spectra, and the MSn-COCONUT subset from MSn-Lib, containing approximately 13,000 spectra, after filtering (see Methods). As a structural reference, we calculated pairwise Tanimoto similarities based on Morgan fingerprints (4,096 bits) derived from annotated SMILES. SpecReBoot was applied using widely adopted spectral similarity metrics, including cosine, modified cosine, Spec2Vec, and MS2DeepScore.

Despite fundamental differences in how these methods compute similarity, edge support exhibited consistent stability patterns. As illustrated for the pesticide dataset (Fig. 4a), selected for its relatively small size, mean similarity matrices differed substantially between methods, whereas edge support was markedly more conserved and, to a certain extent, reflected the underlying structure of pairwise chemical similarity (Fig. 4a–c). Importantly, edge support should not be interpreted as a proxy for the magnitude of chemical similarity itself. While it aligns visually with “ground-truth” chemical relationships, it is not intended as a standalone predictor of chemical similarity and shows lower precision–recall performance than spectral similarity scores alone (Fig. S13–S20). Instead, these observations motivated us to evaluate the use of edge support as a complementary filter applied on top of similarity scores to refine chemically meaningful spectral relationships.

**Fig. 4.**
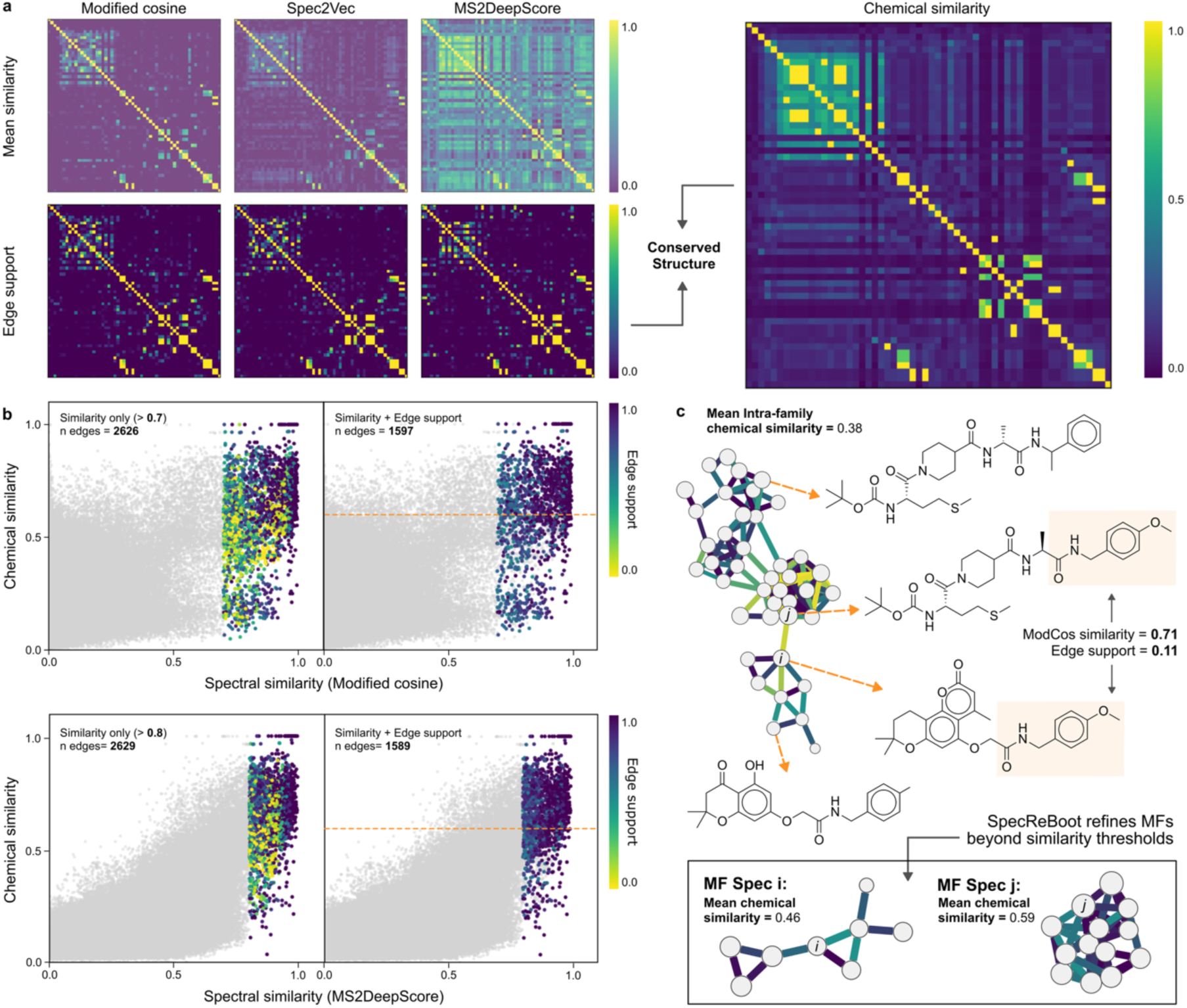
Edge support captures metric-independent spectral relationships and enriches chemically meaningful connections. **a,** pairwise similarity matrices for the pesticide dataset computed using modified cosine, Spec2Vec, and MS2DeepScore (top row), displayed using an identical, fixed scan-ID ordering and without method-specific clustering. The corresponding edge support matrices (bottom row) are shown under the same ordering and compared to the pairwise chemical similarity matrix (right). Despite substantial differences in raw similarity structure across metrics, edge support matrices are highly conserved and show a strong correspondence with the underlying chemical similarity matrix, highlighting stable spectral relationships that are largely independent of the chosen similarity metric. **b,** chemical versus spectral similarity for edges retained using similarity thresholds alone (left panels) or in combination with edge support (right panels). The horizontal dashed line indicates a chemical similarity threshold above 0.7. Applying edge support as a secondary filter substantially reduces the number of retained edges (n indicated in each panel) and enriches connections in regions of higher chemical similarity across both similarity metrics. Color indicates edge support. **c,** representative molecular family showing the impact of edge support on network refinement. Even though the edge between spectra *i* and *j* exceeded the modified cosine similarity threshold, their low edge support reflects inconsistent recovery across bootstrap replicates. Applying edge support as an additional constraint removes unstable connections and yields molecular families that are more chemically coherent as illustrated by the increased intra-family mean chemical similarity.

This complementary behavior was evaluated in detail using the NIH Natural Products Library (Fig. 4b). At this scale, the modified cosine score is stringent, as most pairwise comparisons yield low similarities, and applying a threshold of 0.7 retains only 2,626 edges. Nevertheless, many of these edges correspond to pairs with low chemical similarity. Incorporating edge support (≥ 0.5) as a secondary filter whilst still using a modified cosine threshold of 0.7, i.e., a dual-filter approach, reduced the number of retained edges to 1,581. This subset, herein defined as core edges, exhibited a clear enrichment toward high chemical similarity (Fig. 4b). A similar pattern was observed for MS2DeepScore, although embedding-based similarities are generally shifted toward higher ranges (Fig. S24). Applying a similarity threshold of 0.7 retained 8,378 edges (Fig. S29), and a more stringent threshold of 0.8 retained 2,629 edges, still comprising a significant fraction of pairs with low chemical similarity. Further adding edge support as a dual-filter (≥ 0.5) together with the similarity threshold of 0.8, reduced this set to 1,572 core edges, preferentially enriched in the high-chemical and high-spectral similarity regime (Fig. 4b). A similar scenario was observed for Spec2Vec; however, fewer unsupported edges were retained using similarity thresholds alone, consistent with its more conservative behavior (Fig. S30). Yet, applying the SpecReBoot dual-filter strategy also further refined the set of retained edges for this metric.

In general, edges with low support occurred across the full range of chemical similarity but were enriched among structurally dissimilar pairs (Tanimoto < 0.7). This chemical enrichment can be quantified by comparing the proportion of edges connecting chemically similar pairs before and after applying our dual-filter strategy. Specifically, the fraction of retained edges above the Tanimoto threshold increased by 1.32-fold, 1.33-fold, and 1.31-fold for modified cosine, MS2DeepScore, and Spec2Vec, respectively, relative to similarity thresholding alone in this dataset.

Having established that edge support improves chemical similarity enrichment, we next asked whether it also harmonizes retained edges across different similarity metrics. We therefore compared the overlap among retained edge sets for the NIH Natural Products Library (Table S2). When similarity thresholds were applied alone, all three metrics retained a similar number of edges (∼2,360 per metric), but only moderate pairwise agreement was observed (Jaccard index = 0.52–0.61). Although 1,444 edges were shared across metrics, only 515 of these connected spectra representing chemically similar pairs of molecules (Tanimoto similarity ≥ 0.7), indicating that agreement between metrics does not necessarily imply chemical coherence. Incorporating edge support (≥ 0.5) as a secondary filter reduced the shared edges (n = 783), while increasing its chemical consistency, with approximately half of shared edges now linking spectra representing chemically similar molecules (Tanimoto similarity ≥ 0.7). Notably, this enrichment occurred without increasing the proportion of shared edges, suggesting that edge support does not artificially enforce agreement between similarity measures. Instead, it refines metric-agnostic connections by preferentially removing unstable links and increasing the proportion of “true positive” edges.

Building on this observation, we next asked whether this refined, metric-agnostic core translates into more consistent MN topologies. We constructed MN using our dual-filter strategy, retaining edges satisfying both a similarity threshold (0.7 for cosine-based metrics and Spec2Vec, and 0.8 for MS2DeepScore) and a minimum edge support of 0.5, and compared them to networks constructed using similarity thresholding alone (Fig. S35–S42). In MNs built using solely similarity thresholds, the largest MFs were dominated by low-support edges, whereas smaller components tended to exhibit higher internal stability. Applying edge support ≥ 0.5 “rebooted” topology by removing unstable connections, resulting in more balanced MNs composed of chemically coherent families.

Consistent with this, all “rebooted” networks showed similar degree distributions, with predominantly low-degree nodes and a limited number of moderately connected MFs (Fig. S43). Despite metric-specific differences, particularly for Spec2Vec, global topological statistics revealed comparable overall organization across metrics (Table S3). Average degree (3.3–5.0), network centralization (0.13–0.28), and the number of connected MFs (140–157) were of similar magnitude across networks. Importantly, none of the “rebooted” networks exhibited a highly centralized “hairball“-like structure, and the proportion of singletons remained approximately 40% on average. Together, these results demonstrate that incorporating edge support stabilizes global network organization across similarity metrics and highlights a robust backbone of reproducible spectral relationships.

To illustrate how this topological refinement reflects at the level of individual MFs, we examined a representative MF within the NIH Natural Products Library (Fig. 4c). Here, spectra i and j correspond to metabolites of distinct biosynthetic origin, a linear tetrapeptide and a hybrid phenylpropanoid. Despite sharing only an O-methyl-tyrosine substructure, these compounds and their derivatives were grouped into the same MF based on a single edge exceeding the similarity threshold. Applying SpecReBoot resolved this cluster into two distinct MFs, each exhibiting coherent internal chemistry (Fig. S44–S46), quantitatively supported by an increase in mean intra-family chemical similarity after removing unstable connections.

To evaluate scalability, we applied SpecReBoot to the MSn-COCONUT dataset comprising 12,929 filtered spectra. Using a modified cosine similarity threshold of 0.7 alone, 96,423 edges were retained, including a substantially larger fraction of chemically unrelated pairs than in the previously analyzed datasets based on Tanimoto score comparison. Implementing our dual-filter strategy reduced the number of retained edges to 33,068 and enriched for chemically meaningful connections (Tanimoto similarity ≥ 0.7), indicating improved chemical specificity within individual MFs for all metrics at this scale (Fig. 5, S47–S52). As the SpecReBoot algorithm scales with the number of bootstrap replicates and pairwise spectral comparisons, computational cost increases substantially for large datasets; however, the current NumPy-based implementation helps mitigate this in practice and enables analysis at large-scale, and with appropriate infrastructure, at repository scale. For example, the MSn-COCONUT dataset completed 100 bootstrap replicates in approximately 46 minutes on a single computing node, leveraging the internal parallelization currently implemented for cosine-based scores in *matchms*.

**Fig. 5.**
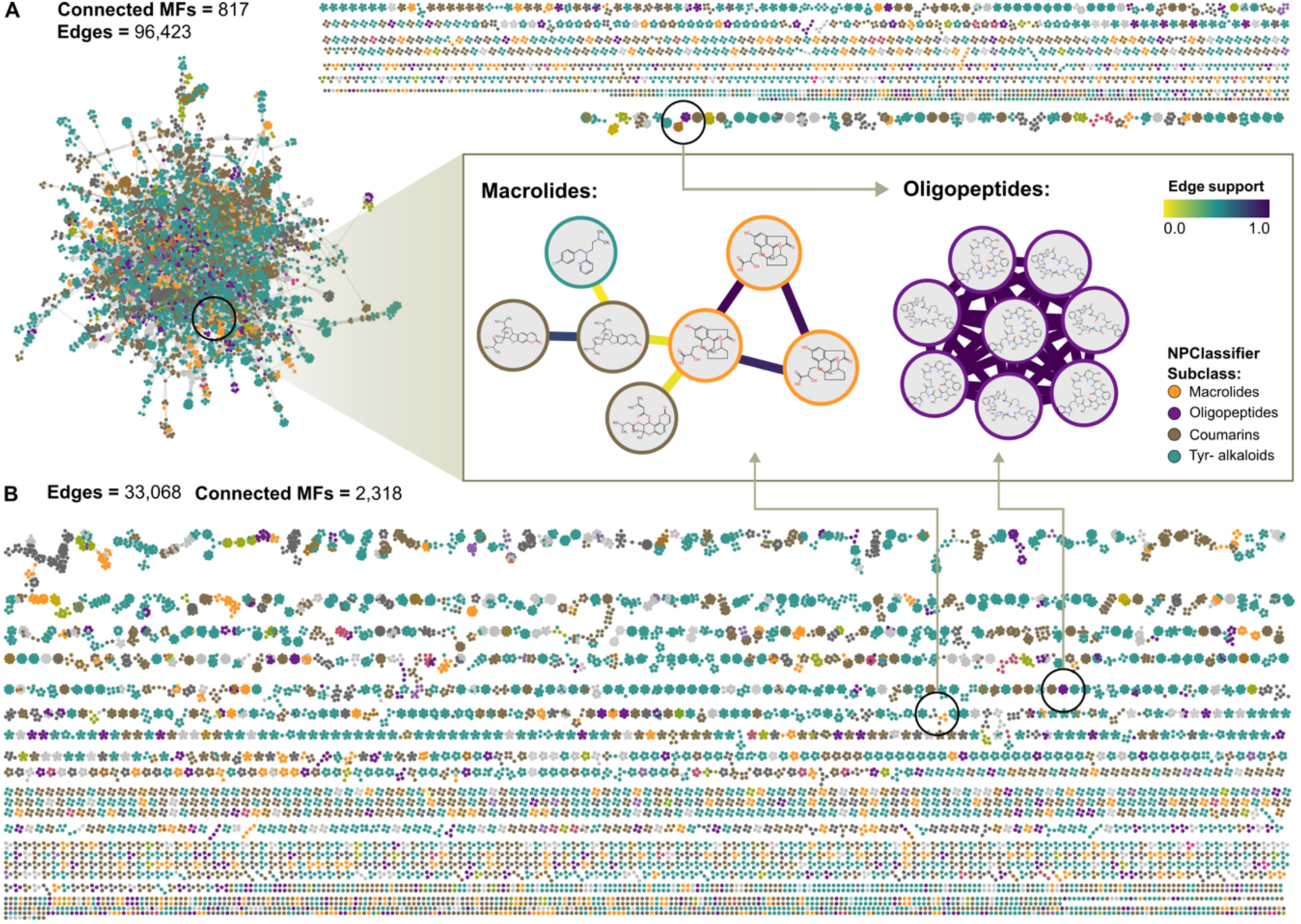
Repository-scale topology refinement in the MSn-COCONUT dataset. **a,** molecular network constructed using modified cosine similarity (≥ 0.7). **b,** molecular network after applying the SpecReBoot dual-filter strategy (similarity ≥ 0.7; edge support ≥ 0.5). Both panels use identical layout parameters. Node colors indicate NPClassifier-derived classifications. Dual filtering reduces unstable cross-class connections, decomposing the centralized topology into modular, chemically coherent molecular families. Insets illustrate the placement of representative molecular families for macrolide polyketides and oligopeptides before and after rebooting, with edges colored by edge support.

This is particularly relevant for large, chemically diverse datasets, where similarity-based networking is more prone to connecting unrelated scaffolds, for example due to recurrent structural motifs shared across distinct compound classes. Consistent with this, the fraction of retained edges above the Tanimoto similarity threshold increased by 2.56-fold, and we also observed a 1.78-fold reduction in average degree and a 2.84-fold increase in the number of connected MFs. Using similarity thresholding alone produced a highly interconnected “hairball” with 9,206 nodes spanning diverse NPClassifier-derived pathway classifications, whereas dual filtering decomposed this structure into more modular, chemically coherent MFs (Fig. 5). This is illustrated by the sulfur-containing curvularin-type macrolides and unguisin-type cycloheptapeptides, whose respective MFs become more chemically coherent after rebooting. For the curvularin-type macrolides, incomplete resolution even under the current settings for our dual-filter strategy highlights how edge support can guide further optimization and enable additional refinement for targeted analysis of specific compound classes in the future.

## Discussion

The introduction of molecular networking has transformed natural product discovery by enabling systematic exploration of MS/MS data and large-scale community annotation efforts. However, despite advances including Feature-Based and Ion Identity MN^6,38,39^, the core principle, linking spectra by pairwise similarity, remains sensitive to noise, missing fragments, chimeric fragmentation spectra, and experimental variability. As a result, most networks lack an explicit, edge-level notion of confidence^15,17,40^, in contrast to other data-driven fields in which uncertainty estimates are a standard practice, such as ultrafast approximations for phylogenetic bootstrap and the confidence index pLDDT recently introduced for AlphaFold 2.0^41,42^. SpecReBoot addresses this challenge by quantifying the stability of spectral relationships under fragment-level perturbations. By adapting non-parametric bootstrapping from phylogenetics to MS/MS data^43,44^, SpecReBoot converts a single similarity score into a distribution-aware assessment and reports on edge support as a confidence layer. Edge support captures how often two spectra are recovered as mutual top-*k* nearest neighbors across bootstrap replicates, even when full-spectrum similarity is diluted by noise or missing fragments (Fig. 2). In practice, this complementary measure can be used as a criterion to filter unstable connections and enrich MNs for chemically related spectral pairs.

SpecReBoot is agnostic to the underlying spectral similarity metric. Across cosine-based and embedding model-based similarity scores, edge support preserves highly consistent structural patterns that closely resemble chemical similarity relationships derived from molecular fingerprints (Fig. 4a). Implementing our dual-filter strategy, in which edge support acts as a complementary filter, reduces network density while retaining chemically enriched connectivity, despite operating on fundamentally different similarity metrics. At the same time, stable metric-specific connections persist after dual filtering, supporting continued development of improved similarity scores.

Beyond cross-metric consistency, SpecReBoot enables fragment-level interpretability by documenting bootstrap histories at the bin level. This allows direct inspection of which *m/z* bins support an edge and how this evidence varies across datasets and scoring methods, exemplified for the aerucyclamides (Fig. 2). Such fragment-level insights are naturally compatible with and complementary to substructure-oriented approaches such as MS2LDA and consensus-based strategies such as Multiple Mass Spectral Alignment (MMSA)^45,46^. We note that recording the bootstrap history adds computational overhead during bootstrapping, and this feature is therefore optional and disabled by default, as currently the primary performance bottleneck lies in network construction rather than history storage.

We next assessed whether these benefits extend to repository-scale datasets. Analysis of the MSn-COCONUT subset showed that confidence-aware MN benefits hold across large, chemically diverse libraries. While similarity thresholding alone produced highly interconnected topologies across similarity metrics, incorporating edge support reduced unstable edges and decomposed networks into more modular MFs (Fig. 5). Consistent with this topological “rebooting,” chemical enrichment increased markedly for the resulting core edges, alongside a reduced average degree and an increased number of connected components. Together, these results support edge support as a practical confidence layer for scalable MN in public repositories.

### Confidence-aware molecular networking enables discovery of new chemistry

In the *Diaporthe caliensis* case study, SpecReBoot rescued robust low-similarity edges that reconciled relationships previously supported only by shared MS2LDA-derived Mass2Motifs, reconnecting fragmented polyketide-lactones into a coherent molecular family (Fig. 3). In particular, the relationship between caliensolides A and B, which was missed under conventional similarity thresholding, was repeatedly recovered under fragment resampling. Applying an exploratory edge support threshold then “rebooted” network topology by filtering unstable edges while incorporating high-support links, revealing polyketide-lactone diversity that was fragmented in the original study. This confidence-aware strategy guided prioritization beyond previously annotated molecules and led to the isolation of an unprecedented macrolactone polyketide scaffold, caliensomycin.

Moreover, placing caliensomycin in the same molecular family as phomol provided a concrete hypothesis of shared biosynthetic origin and motivated targeted genome mining. Comparative analysis of candidate loci identified a biosynthetic gene cluster region encoding two PKSs and expanded oxidative tailoring capacity, supporting a model in which conserved macrolactone assembly is followed by late-stage oxidative remodeling, as proposed for macrolactone diversification in related systems^34,47,48^. Together, this case study illustrates how SpecReBoot can translate confidence-aware spectral relationships into actionable hypotheses, linking network topology “rebooting” to natural products discovery and biosynthetic prioritization.

### Integration, limitations, and practical considerations

SpecReBoot is designed for compatibility with widely used metabolomics infrastructures. Implemented within the *matchms* ecosystem, it supports cosine-based approaches and embedding-based similarity models such as Spec2Vec and MS2DeepScore and accepts GNPS2 molecular networking outputs as inputs. These design choices enable straightforward adoption while preserving existing workflows.

SpecReBoot introduces several user-defined parameters, including the number of nearest neighbors (*k*) for mutual kNN neighbors’ detection and the number of bootstrap replicates (*B*). Although these parameters exhibited stable behavior in our analyses, optimal values may depend on dataset size, chemical diversity, and downstream application; we therefore recommend reporting (*k*, *B*) alongside networking parameters. As with other resampling-based approaches, bootstrapping increases computational cost relative to single similarity calculations, although algorithmic optimization and parallelization strategies currently implemented in SpecReBoot mitigate this challenge in practice. Moreover, the current resampling strategy does not weight fragment peaks by their frequency across spectra, meaning that fragments may be sampled at unequal rates depending on spectral complexity and the number of bootstrap replicates selected. Future efforts should explore resampling strategies that account for spectral complexity and fragment peak distributions inherent to tandem mass spectrometry data.

This framework provides a foundation for confidence-aware molecular networking rather than a finalized implementation. Systematic evaluation of alternative perturbation schemes, hierarchical resampling strategies, and model-based confidence estimators may further improve the framework. Finally, edge support is not intended to replace similarity scores or embeddings, it provides a complementary confidence layer that filters unstable connections and prioritizes reproducible relationships. Because edge support alone does not capture the magnitude of chemical similarity, it should therefore be interpreted jointly with conventional mass spectral similarity measures.

### Outlook

Taken together, SpecReBoot shifts molecular networking from a purely deterministic framework toward one that explicitly accounts for confidence and reproducibility, enabling more reliable interpretation of spectral relationships in complex mixtures of small molecules. By identifying a robust, metric-agnostic core of spectral connections that is consistently recovered across fundamentally different similarity metrics, SpecReBoot helps to harmonize molecular network topology without artificially forcing agreement between similarity scores. It also offers a new avenue for interpreting ML-based methods: embedding models such as Spec2Vec and MS2DeepScore expand the scope of detectable relationships, yet their limited explainability remains a key challenge. By linking edge support to fragment-level resampling information, SpecReBoot reconnects learned similarities to underlying spectral evidence, providing an interpretable confidence layer that complements AI-driven similarity scoring and may guide the development of improved embedding-based approaches.

At the same time, we demonstrate how individual similarity metrics retain stable, metric-specific connections not captured by others, underscoring that any single similarity score provides an incomplete view of spectral relatedness. This opens opportunities for consensus network construction, where a shared core of reproducible edges defines a common backbone and metric-specific links capture additional chemical diversity. Looking forward, bootstrap-derived confidence measures will enable confidence-aware metabolomics analyses that include principled network comparison, harmonization across datasets and similarity metrics. Furthermore, the concept can be extended to obtain improved annotation^49^, repository-wide searches^50^, prioritization, and substructure discovery with reliability measures. While the current study intended to provide a general overview of the framework applicability, fragment-level insights provided by SpecReBoot are expected to deepen our understanding of fragmentation patterns across diverse chemical contexts, naturally extending toward optimization-based approaches or class-specific strategies, informed by and supplementing MS2Query and CANOPUS^51,52^. Given that our resampling approach is context-agnostic, beyond natural product discovery, SpecReBoot provides a general framework for assessing robustness of MS/MS-derived relationships, with potential applications in exposomics, clinical metabolomics, and large-scale mining of public spectral repositories.

## Methods

### Datasets and accessions

#### RiPP reference set construction

MS/MS spectra were mined from the GNPS platform using publicly available MGF files with associated structural identifiers (SMILES, InChIKey, InChI or InChIAux). Duplicate entries were removed based on identical structural identifiers and/or redundant spectrum metadata. For spectra lacking SMILES, structural representations were generated from available InChI-based identifiers where possible. Compounds were classified using NPClassifier via the GNPS2 web service (npclassifier.gnps2.org/classify; accessed November 2025) by querying the REST API with SMILES strings for each candidate compound^25^. Peptide-class candidates were manually curated against the literature to confirm RiPP biosynthetic origin. The final RiPP reference library comprised 24 curated MS/MS spectra with literature-supported structures and biosynthetic context.

#### Benchmark datasets

The pesticides dataset, introduced by the *matchms* team, comprised 76 reference spectra (pesticides.mgf). The NIH Natural Products Library dataset comprised 1,267 reference spectra (GNPS-NIH-NATURALPRODUCTSLIBRARY.mgf). The MSn-COCONUT benchmark set was derived from a harmonized collection of MS/MS spectra assembled from GNPS and MSn-Lib^53^. From the initial 1,017,531 spectra, only MS2 spectra acquired in positive ionization mode were retained. To restrict the dataset to natural products, annotated spectra were matched by InChIKey connectivity block (first 14 characters since it contains the 2D-structural information) against the COCONUT database^55^, retaining only spectra whose compounds were present in COCONUT (407,029 registered compounds). This yielded 15,947 spectra, of which 12,929 remained after preprocessing (see below) and were used for downstream analyses.

### Implementation, required inputs and outputs

SpecReBoot operates on standard GNPS-compatible data formats and supports two execution modes: (i) full_matchms mode, which starts from a MGF file (can be derived from mzmine for example) and exports new networks in GraphML format (can be read-in by CytoScape), and (ii) gnps_workflow mode, which starts from a GNPS2 GraphML network with its merged MGF (or the original preprocessed MGF file) and appends the bootstrap-derived supported edges.

### Required inputs

1. **MS/MS spectra (both modes):** An MGF file containing MS/MS spectra and associated metadata. For datasets assembled from multiple sources or libraries, each spectrum must include a stable, unique identifier (e.g., feature_id) or be added. This is required to ensure consistent spectrum tracking and unambiguous mapping across intermediate outputs and molecular network nodes.
2. **GNPS2 workflow mode only (gnps_workflow):** a GraphML network generated by a GNPS2 molecular job, plus the corresponding merged MGF used to construct that GNPS2 network.

### Key user parameters

- Binning resolution (number of decimal places; default: 2 decimals), number of bootstrap replicates *B*, mutual kNN parameter *k*, similarity metric (cosine, modified cosine, Spec2Vec, MS2DeepScore), optional dual thresholds: similarity threshold (ι−_sim_) and support threshold (ι−_sup_), as well as the number of parallel worker threads (n_jobs), and number of replicates per thread-pool batch (batch-size).

### Primary outputs

- **Pairwise tables (CSV):** (i) bootstrap mean similarity and (ii) edge support for spectrum pairs.
- **Networks (GraphML):** depending on the SpecReBoot mode used, either newly constructed GraphML networks or a GNPS2 GraphML with appended bootstrap-derived edge support attributes.
- **Optional provenance/history file:** bootstrap replicate-level diagnostic information (e.g., sampled/missing bin sets) when enabled for reproducibility and debugging.

### MS/MS preprocessing

All datasets were imported from MGF format. For datasets assembled from multiple libraries, a unique identifier field (feature_id) was added to the metadata to ensure stable spectrum tracking and unambiguous mapping across intermediate outputs. Spectra were processed using *matchms* (v0.31.0) default harmonization and cleaning pipelines (DEFAULT_FILTERS and CLEAN_PEAKS)^37,54^. By default, intensities were normalized to unit height (scaled by the maximum intensity). Cleaning steps followed the *matchms* pipeline implementation and included precursor-window peak removal (±17 Da around the precursor, excluding the precursor peak), retention of peaks within ±50 Da of at least one of the six most intense peaks, and an upper bound of 1,000 fragment peaks per spectrum. Users can modify these filtering steps depending on dataset characteristics and metadata completeness.

### Peak binning and global fragment-bin space

Fragment *m/z* values were discretized by rounding to a user-defined number of decimal places (default: two decimals; bin width 0.01 Da). A dataset-wide global bin list *G* was constructed as the union of all discretized fragment *m/z* values observed across all spectra, yielding P = |G| unique bins. Each spectrum (now spectra binned) was represented by its discretized fragment *m/z* values and the corresponding normalized intensities.

### SpecReBoot algorithm

#### Bootstrap resampling over global bins

SpecReBoot applies bootstrap perturbations by repeatedly constructing replicate-specific subsets of the global bin list *G*. For each bootstrap replicate b, P bin indices are sampled uniformly with replacement from 0 to P, to obtain a unique sampled bin subset *G*^(*b*)^ ⊆ *G* where each bin value represents a collection of binned original fragments. As the sampling is with replacement, each replicate resamples ∼36.8% of global bins, with the remaining (on average) 1−(1−1/P)^P ≈ 63.2% of the global bins being retained. Each original spectrum is converted into a pseudo-spectrum by retaining only those peaks whose discretized *m/z* value is contained in *G*^(*b*)^; all other peaks per pseudo-spectrum are removed. Importantly, this process does not shuffle or exchange intensities across spectra: intensities remain attached to the original spectrum and are only retained or removed in each bootstrap replicate, and no re-normalization takes place (Fig. S55). As the binned spectra are typically sparse relative to the global bin space, a given replicate may remove few peaks from some spectra and many from others, depending on each spectrum’s fragment composition. This motivates the use of multiple bootstrap replicates to robustly estimate pairwise similarity and edge stability across the dataset.

If a spectrum contains no peaks after masking, a dummy peak (*m/z* set to the first global bin and intensity 0.0) is inserted to ensure similarity computation remains well-defined; such spectra effectively contribute zero similarity and do not form mutual kNN relationships for that replicate. For each replicate, pairwise similarities are computed using the selected similarity metric, producing a replicate similarity matrix. Two outputs are then aggregated across replicates:

1. **Mean similarity**: defined for each spectrum pair (i,j) as the mean of replicate similarity values across B bootstraps.
2. **Edge support**: defined for each spectrum pair (i,j) as the fraction of bootstraps in which spectra i and j are recovered as **mutual** top-*k* nearest neighbors. Specifically, for each spectrum i, the top-*k* neighbors are obtained by ranking similarity values within replicate b (excluding self), and an undirected edge is counted once per replicate when i is in the top-*k* neighbors of j and j is in the top-*k* neighbors of i. Edge support is reported as support_ij_ = c_ij_/B, where c_ij_ is the number of replicates in which the mutual kNN relationship is registered (default: *k*=5).

### Spectral similarity metrics and configurations

Cosine and modified cosine similarities were computed using *matchms* similarity implementations with user-defined precursor *m/z* tolerance where applicable. Spec2Vec similarities were computed using a model retrained for positive ionization mode spectra using curated positive-mode training spectra from [https://zenodo.org/records/12543129], files associated can be downloaded here: https://zenodo.org/records/15857387; and MS2DeepScore similarities were computed using a publicly available pretrained model [https://zenodo.org/records/17826815]. Model sources and versions are reported in the Code availability section.

For network construction, SpecReBoot supports dual filtering, i.e., applying thresholds on both the bootstrap mean similarity and edge support. In this way, an edge between spectra i and j is retained only if bootstrap mean similarity ≥ ι−_sim_ and edge support ≥ ι−_sup,_ where ι−_sim_ and ι−_sup_ are user-defined.

### Network construction and graph parameters

SpecReBoot supports two execution modes:

**(i) full_matchms mode.** Starting from an MGF file, SpecReBoot performs preprocessing, binning, and bootstrap similarity estimation, and can export up to three GraphML networks:
1. a **base network** thresholded solely on bootstrap mean similarity (bootstrap mean similarity ≥ ι−_sim_);
2. a **dual-threshold network** requiring both bootstrap mean similarity and edge support (bootstrap mean similarity ≥ ι−_sim_ and edge support ≥ ι−_sup_); and
3. a **core/rescue network** that labels edges as *core* (high similarity and high support) or *rescued* (lower similarity but high support) based on user-defined criteria.
**(ii) gnps_workflow mode.** SpecReBoot loads a GNPS-exported GraphML network and the corresponding merged MGF, maps spectra to nodes via identifiers, preserves all original node and edge attributes, and appends bootstrap-derived edge attributes (bootstrap mean similarity and edge support). Optionally, it generates rescued-edge annotations following the same core/rescue logic. Also here, the user can define threshold criteria.

### Benchmarking against chemical similarity

To evaluate whether edge support enriches chemically meaningful relationships beyond similarity alone, structural similarity was computed for annotated compounds using RDKit Morgan fingerprints (radius 2, 4,096 bits) and Tanimoto similarity. Pairwise similarity tables were generated by merging bootstrap mean similarity, edge support, and Tanimoto similarity by spectrum identifiers.

### Statistical analysis

Binary classification performance was assessed using receiver operating characteristic (ROC) and precision–recall (PR) analyses, defining “chemically similar” pairs as those with Tanimoto threshold ≥ **0.7**. In addition, we visualized the relationship between mean spectral similarity (x-axis) and structural similarity (y-axis), with points that are colored to indicate the amount of edge support, to highlight stable spectral relationships across bootstrap replicates.

### Experimental procedures for *Diaporthe caliensis* and structure elucidation Fungal genomic DNA extraction, and genome sequencing

For genomic DNA isolation, *D. caliensis* was cultivated on sterile 50 µm nylon mesh placed on the surface of Petri dishes containing potato dextrose agar (PDA). Cultures were prepared in duplicate to obtain sufficient biomass for high-quality DNA extraction. After 7 days of cultivation, the mycelia developing on the mesh was harvested by scraping with a sterile scalpel, snap-frozen in liquid nitrogen, and stored at −80 °C until further processing. Genomic DNA was extracted using the NucleoBond HMW DNA kit following the manufacturer’s protocol. Briefly, frozen mycelia were ground under liquid nitrogen and lysed in the presence of proteinase K. DNA was subsequently purified using the kit’s affinity columns and eluted in HE buffer. DNA integrity and purity were assessed by electrophoresis on a 1% agarose gel prior to sequencing. Whole-genome sequencing was performed using Illumina technology. Raw sequencing reads were subjected to quality assessment and adapter trimming using FASTQ, and de novo genome assembly was carried out with SPAdes version 3.15.3.

### Cultivation of *D. caliensis* and isolation of caliensomycin

*Diaporthe caliensis* CBS 149729, originally isolated from the medicinal plant *Otoba gracilipes* in Cali, Colombia, was cultivated as previously described^30^. Briefly, seed cultures were prepared in 200 mL yeast malt extract broth (YM broth) by transferring colonies from fresh cultures in yeast malt agar (YM agar) and incubating at 23 °C with agitation at 140 rpm for 7 days. Aliquots (6 mL) of this seed culture were then used to inoculate 12 flasks containing solid oat medium (OFT), which were incubated statically at 23 °C for 15 days. Metabolite extraction was performed following the procedure reported in Charria-Girón et al^30^, yielding 1,153 mg of crude extract. The crude extract was pre-fractionated by flash chromatography (Grace Reveleris, Columbia, MD, USA) using a 40 g silica cartridge. The mobile phase consisted of solvent A (DCM), solvent B (acetone), and solvent C [(DCM/acetone 8:2):MeOH], with the following gradient: 100% A for 5 min, increasing to 100% B over 20 min, followed by a transition to 100% C over 15 min and a final hold at 100% C for 5 min. Five fractions (F1–F5) were collected and subsequently analyzed for the presence of previously uncharacterized metabolites. Fraction F3 was purified by preparative RP-HPLC (Synergi Polar-RP, 250 Å∼ 50 mm, 10 μm; Phenomenex) with H₂O (A) and ACN (B) (0.1% formic acid) at 45 mL/min (UV 210, 240, and 300 nm), using a 5–32% B (15 min), 32–36% B (40 min), 36–100% B (25 min) gradient and 100% B (5 min), affording caliensomycin (1.4 mg, tR = 21–23 min).

### Spectral data

Optical rotations were recorded employing an MCP 150 circular polarimeter (Anton Paar, Seelze, Germany) at 20°C. UV/Vis spectra were recorded with a UV-2450 spectrophotometer (Shimadzu, Kyoto, Japan). Spectral data were measured in MeOH (Uvasol, Merck, Darmstadt, Germany) for all compounds. The 1D- and 2D-nuclear magnetic resonance (NMR) spectra were recorded with an Avance III 700 spectrometer with a 5 mm TCI cryoprobe (^1^H NMR: 700 MHz, ^13^C: 175 MHz, Bruker, Billerica, MA, USA) and an Avance III 500 spectrometer (^1^H NMR: 500 MHz, ^13^C: 125 MHz, Bruker, Billerica, MA, USA). The chemical shifts δ were referenced to the solvents DMSO-*d* _6_ (^1^H, δ = 2.50; ^13^C, δ = 39.51).

### Caliensomycin

White powder; UV (MeOH) λmax (log ε) 263 (3.35), (log ε) 202.5 (3.06); HRESI-MS: *m*/*z* 469.2719 [M + H]^+^ (calculated for C_24_H_39_O_8_^+^: 469.5882 Da).

## Data and Code Availability

SpecReBoot codebase is publicly available in our GitHub repository (https://github.com/ECharria/SpecReBoot). The documentation is available in the README.md file for installation and execution. The datasets used for benchmarking and the case study with the Spec2Vec + MS2DeepScore models and the results from this work are stored in https://zenodo.org/records/18466976.

## Author contributions

E.C.-G., L.R.T.-O. and J.J.J.vdH. conceived the concept behind SpecReBoot. E.C.-G. and L.R.T.-O. developed the methodology, implemented the computational framework, performed data analysis, and led the manuscript writing. J.M.G. contributed the RiPP-associated case study and performed manual curation of public spectral repositories. Y.M.F. supervised and supported the initial analysis of the experimental part of the study. N.H.C. contributed the genome analysis, sequencing data, and project coordination. F.S. performed structural elucidation analysis and supervised the experimental part of the study. M.H.M. contributed to the initial development of the SpecReBoot concept, interpretation of results, and performed critical revision of the manuscript. J.J.J.v.d.H. supervised the project, performed critical interpretations of results, and performed manuscript writing and editing. All authors reviewed and approved the final version of the manuscript.

## Acknowledgements

We thank Esther Surges for recording NMR data and Ulrike Beutling for performing tandem mass spectrometry measurements related to the *D. caliensis* case study. We also thank Jorick van IJcken for his contributions to the codebase. Access to genetic resources for the study of *D. caliensis* was granted under contract RGE244-44#190 by the Autoridad Nacional de Licencias Ambientales (ANLA), Ministerio de Ambiente y Desarrollo Sostenible, Colombia. Whole-genome sequencing was conducted within the framework of a BiGTREE mobility scholarship awarded to Rafael Góngora. Laboratory work, including molecular biology procedures, sequence analysis, and genome assembly, was carried out at the University of Oslo under the supervision of Professor Inger Skrede, with results reported in the corresponding BiGTREE scholarship activity report. Transport of fungal cultures was conducted under export and import authorization for specimens of biological diversity listed in the appendices of the CITES Convention (Permit No. 3018), issued by ANLA. L.R.T.-O. gratefully recognizes the Marie Skłodowska-Curie grant under the European Union’s Horizon Europe programme MAGiC-MOLFUN (grant no. 101072485).

## References

1. Alseekh S, Fernie AR. Metabolomics 20 years on: what have we learned and what hurdles remain?. Plant J. 94, 933–942 (2018). doi:10.1111/tpj.13950.

2. Wishart DS. Metabolomics for investigating physiological and pathophysiological processes. Physiol. Rev. 99(4), 1819–1875 (2019). doi:10.1152/physrev.00035.2018.

3. Bauermeister A, Mannochio-Russo H, Costa-Lotufo LV, Jarmusch AK, Dorrestein PC. Mass spectrometry-based metabolomics in microbiome investigations. Nat. Rev. Microbiol. 20(3), 143–160 (2022). doi:10.1038/s41579-021-00621-9.

4. Wang M, Carver JJ, Phelan VV, et al. Sharing and community curation of mass spectrometry data with Global Natural Products Social Molecular Networking. Nat. Biotechnol. 34, 828–837 (2016). doi:10.1038/nbt.3597.

5. Jarmusch SA, van der Hooft JJJ, Dorrestein PC, Jarmusch AK. Advancements in capturing and mining mass spectrometry data are transforming natural products research. Nat. Prod. Rep. 38(11), 2066–2082 (2021). doi:10.1039/d1np00040c.

6. Nothias LF, Petras D, Schmid R, et al. Feature-based molecular networking in the GNPS analysis environment. Nat. Methods. 17, 905–908 (2020). doi:10.1038/s41592-020-0933-6.

7. Aron AT, Gentry EC, McPhail KL, et al. Reproducible molecular networking of untargeted mass spectrometry data using GNPS. Nat. Protoc. 15, 1954–1991 (2020). doi:10.1038/s41596-020-0317-5.

8. Beniddir MA, Kang KB, Genta-Jouve G, Huber F, Rogers S, van der Hooft JJJ. Advances in decomposing complex metabolite mixtures using substructure- and network-based computational metabolomics approaches. Nat. Prod. Rep. 38, 1967–1993 (2021). doi:10.1039/d1np00023c.

9. van der Hooft JJJ, Padmanabhan S, Burgess KE, Barrett MP. Urinary antihypertensive drug metabolite screening using molecular networking coupled to high-resolution mass spectrometry fragmentation. Metabolomics. 12, 125 (2016a). doi:10.1007/s11306-016-1064-z.

10. Huber F, Ridder L, Verhoeven S, et al. Spec2Vec: Improved mass spectral similarity scoring through learning of structural relationships. PLoS Comput. Biol. 17, e1008724 (2021). doi:10.1371/journal.pcbi.1008724.

11. Bittremieux W, Schmid R, Huber F, van der Hooft JJJ, Wang M, Dorrestein PC. Comparison of cosine, modified cosine, and neutral loss based spectrum alignment for discovery of structurally related molecules. J. Am. Soc. Mass. Spectrom. 33, 1733–1744 (2022). doi:10.1021/jasms.2c00153.

12. Bushuiev R, Bushuiev A, Samusevich R, et al. Self-supervised learning of molecular representations from millions of tandem mass spectra using DreaMS. Nat. Biotechnol. (2025). doi: 10.1038/s41587-025-02663-3.

13. de Jonge, N.F., Chekmeneva, E., Schmid, R. et al. Cross ionization mode chemical similarity prediction between tandem mass spectra in metabolomics. Nat. Commun. 17, 2483 (2026). 10.1038/s41467-026-69083-y.

14. van der Hooft JJJ, Wandy J, Barrett MP, Burgess KE, Rogers S. Topic modeling for untargeted substructure exploration in metabolomics. Proc. Natl. Acad. Sci. USA 113, 13738–13743 (2016b). doi:10.1073/pnas.1608041113.

15. Sachsenberg T, Pino LK, Brunet M, Bludau I, Kohlbacher O, Vizcaino JA, Bittremieux W. Perspectives in computational mass spectrometry: recent developments and key challenges. Bioinformatics Advances 5, vbaf301 (2025). doi:10.1093/bioadv/vbaf301.

16. Stein S. Mass spectral reference libraries: an ever-expanding resource for chemical identification. Anal. Chem. 84, 7274–7282 (2012). doi:10.1021/ac301205z.

17. Lawson TN, Weber RJ, Jones MR, et al. msPurity: Automated evaluation of precursor ion purity for mass spectrometry-based fragmentation in metabolomics. Anal. Chem.89(4), 2432–2439 (2017). doi:10.1021/acs.analchem.6b04358.

18. Clark TN, Houriet J, Vidar WS, et al. Interlaboratory comparison of untargeted mass spectrometry data uncovers underlying causes for variability. J. Nat. Prod. 84, 824–835 (2021). doi:10.1021/acs.jnatprod.0c01376.

19. El Abiead Y, Mohanty I, Xing S, et al. A perspective on unintentional fragments and their impact on the dark metabolome, untargeted profiling, molecular networking, public data, and repository scale analysis. JACS Au 5, 5828–5850 (2025). doi:10.1021/jacsau.5c01063.

20. Arnison PG, Bibb MJ, Bierbaum G, et al. Ribosomally synthesized and post-translationally modified peptide natural products: overview and recommendations for a universal nomenclature. Nat. Prod. Rep. 30, 108–160 (2013). doi:10.1039/c2np20085f.

21. Zdouc MM, van der Hooft JJJ, Medema MH. Metabolome-guided genome mining of RiPP natural products. Trends Pharmacol. Sci. 44, 532–541 (2023). doi:10.1016/j.tips.2023.06.004.

22. Skinnider MA, Johnston CW, Edgar RE, et al. Genomic charting of ribosomally synthesized natural product chemical space facilitates targeted mining. Proc. Natl. Acad. Sci. U S A. 113, E6343–E6351 (2016). doi:10.1073/pnas.1609014113.

23. Nie Q, Sun C, Liu S, Gao X. Correction to “Exploring bioactive fungal RiPPs: Advances, challenges, and future prospects”. Biochemistry 64, 3485 (2025). doi:10.1021/acs.biochem.5c00383.

24. Montalbán-López M, Scott TA, Ramesh S, et al. New developments in RiPP discovery, enzymology and engineering. Nat Prod Rep. 38, 130–239 (2021). doi:10.1039/d0np00027b.

25. Kim HW, Wang M, Leber CA, et al. NPClassifier: A deep neural network-based structural classification tool for natural products. J. Nat. Prod. 84(11), 2795–2807 (2021). doi:10.1021/acs.jnatprod.1c00399.

26. Portmann C, Blom JF, Kaiser M, Brun R, Jüttner F, Gademann K. Isolation of aerucyclamides C and D and structure revision of microcyclamide 7806A: heterocyclic ribosomal peptides from *Microcystis aeruginosa* PCC 7806 and their antiparasite evaluation. J. Nat. Prod. 71, 1891–1896 (2008). doi:10.1021/np800409z.

27. Portmann C, Blom JF, Gademann K, Jüttner F. Aerucyclamides A and B: Isolation and synthesis of toxic ribosomal heterocyclic peptides from the cyanobacterium *Microcystis aeruginosa* PCC 7806. J. Nat. Prod. 71, 1193–1196 (2008). doi:10.1021/np800118g.

28. Ziemert N, Ishida K, Quillardet P, et al. Microcyclamide biosynthesis in two strains of *Microcystis aeruginosa*: from structure to genes and vice versa. Appl. Environ. Microbiol. 74, 1791–1797 (2008). doi:10.1128/AEM.02392-07.

29. Charria-Girón E, Espinosa MC, Zapata-Montoya A, et al. Evaluation of the antibacterial activity of crude extracts obtained from cultivation of native endophytic fungi belonging to a tropical montane rainforest in Colombia. Front. Microbiol. 12, 716523 (2021). doi:10.3389/fmicb.2021.716523.

30. Charria-Girón E, Marin-Felix Y, Beutling U, et al. Metabolomics insights into the polyketide-lactones produced by *Diaporthe caliensis* sp. nov., an endophyte of the medicinal plant *Otoba gracilipes*. Microbiol. Spectr. 11, e0274323 (2023a). doi:10.1128/spectrum.02743-23.

31. Charria-Girón E, Vasco-Palacios AM, Moncada B, Marin-Felix Y. Colombian fungal diversity: Untapped potential for diverse applications. Microbiol. Res. 14, 2000–2021 (2023b). doi:10.3390/microbiolres14040135.

32. Hoyos LV, Vasquez-Muñoz LE, Osorio Y, et al. Tailored culture strategies to promote antimicrobial secondary metabolite production in *Diaporthe caliensis*: a metabolomic approach. Microb. Cell Fact. 23, 328 (2024). doi:10.1186/s12934-024-02567-y.

33. Blin K, Shaw S, Vader L, et al. antiSMASH 8.0: extended gene cluster detection capabilities and analyses of chemistry, enzymology, and regulation. Nucleic Acids Res. 53(W1), W32–W38 (2025). doi:10.1093/nar/gkaf334.

34. Trenti F, Lebe K, Adelin E, et al. Investigating the biosynthesis of Sch-642305 in the fungus *Phomopsis* sp. CMU-LMA. RSC Advances 10, 27369–27376 (2020). doi: 10.1039/D0RA05311B.

35. Cardoza RE, McCormick SP, Izquierdo-Bueno I, et al. Identification of polyketide synthase genes required for aspinolide biosynthesis in *Trichoderma arundinaceum*. Appl. Microbiol. Biotechnol. 106, 7153–7171 (2022). doi:10.1007/s00253-022-12182-9.

36. Snider BB, Zhou J. Synthesis of (+)-sch 642305 by a biomimetic transannular Michael reaction. Org. Lett. 8, 1283–1286 (2006). doi:10.1021/ol052948+.

37. Huber F, Verhoeven S, Meijer C, et al. matchms-processing and similarity evaluation of mass spectrometry data. J. Open Source Softw. 5, 2411 (2020). doi:10.21105/joss.02411.

38. Schmid R, Petras D, Nothias LF, et al. Ion identity molecular networking for mass spectrometry-based metabolomics in the GNPS environment. Nat. Commun. 12, 3832 (2021). doi:10.1038/s41467-021-23953-9.

39. Gentry EC, Collins SL, Panitchpakdi M, et al. Reverse metabolomics for the discovery of chemical structures from humans. Nature. 626, 419–426 (2024). doi:10.1038/s41586-023-06906-8.

40. Gloaguen Y, Kirwan JA, Beule D. Deep learning-assisted peak curation for large-scale LC-MS metabolomics. Ana.l Chem. 94, 4930–4937 (2022). doi:10.1021/acs.analchem.1c02220.

41. Minh BQ, Nguyen MA, von Haeseler A. Ultrafast approximation for phylogenetic bootstrap. Mol. Biol. Evol. 30, 1188–1195 (2013). doi:10.1093/molbev/mst024.

42. Vander Meersche Y, Diharce J, Gelly JC, Galochkina T. Flexibility or uncertainty? A critical assessment of AlphaFold 2 pLDDT. Structure. 33, 2157–2163.e2 (2025). doi:10.1016/j.str.2025.09.001.

43. Felsenstein J. Confidence limits on phylogenies: An approach using the bootstrap. Evolution 39, 783–791 (1985). doi:10.1111/j.1558-5646.1985.tb00420.x.

44. Efron B, Halloran E, Holmes S. Bootstrap confidence levels for phylogenetic trees. Proc. Natl. Acad. Sci. U S A 93, 13429–13434 (1996). doi:10.1073/pnas.93.23.13429.

45. Torres-Ortega LR, Dietrich J, Wandy J, Mol H, van der Hooft JJJ. Large-scale discovery and annotation of hidden substructure patterns in mass spectrometry profiles. Preprint at 10.1101/2025.06.19.659491.

46. Lau A, Wang X, Xu T, et al. Multiple spectrum alignment for molecular networking exploration and discovery. J. Am. Soc. Mass Spectrom. 37, 126–134 (2026). doi:10.1021/jasms.5c00237.

47. Zabala AO, Chooi YH, Choi MS, Lin HC, Tang Y. Fungal polyketide synthase product chain-length control by partnering thiohydrolase. ACS Chem. Biol. 9, 1576–1586 (2014). doi:10.1021/cb500284t.

48. Guengerich FP. Mechanisms of cytochrome P450-catalyzed oxidations. ACS Catal. 8, 10964–10976 (2018). doi:10.1021/acscatal.8b03401.

49. de Jonge NF, Mildau K, Meijer D, et al. Good practices and recommendations for using and benchmarking computational metabolomics metabolite annotation tools. Metabolomics 18, 103 (2022). doi:10.1007/s11306-022-01963-y.

50. Wang M, Jarmusch AK, Vargas F, et al. Mass spectrometry searches using MASST. Nat. Biotechnol. 38(1), 23–26 (2020). doi:10.1038/s41587-019-0375-9.

51. de Jonge NF, Louwen JJR, Chekmeneva E, et al. MS2Query: reliable and scalable MS2 mass spectra-based analogue search. Nat. Commun. 14(1), 1752 (2023). doi:10.1038/s41467-023-37446-4.

52. Dührkop K, Nothias LF, Fleischauer M, et al. Systematic classification of unknown metabolites using high-resolution fragmentation mass spectra. Nat. Biotechnol. 39(4), 462–471 (2021). doi:10.1038/s41587-020-0740-8.

53. Brungs C, Schmid R, Heuckeroth S, et al. MS^n^Lib: efficient generation of open multi-stage fragmentation mass spectral libraries. Nat. Methods 22, 2028–2031 (2025). doi:10.1038/s41592-025-02813-0.

54. de Jonge NF, Hecht H, Strobel M, Wang M, van der Hooft JJJ, Huber F. Reproducible MS/MS library cleaning pipeline in matchms. J. Cheminform. 16, 88 (2024). doi:10.1186/s13321-024-00878-1.

55. Chandrasekhar V, Rajan K, Kanakam SRS, et al. COCONUT 2.0: a comprehensive overhaul and curation of the collection of open natural products database. Nucleic Acids Res. 53(D1), D634–D643 (2025). doi:10.1093/nar/gkae1063.

