## Supplementary information for "Hidden molecular relationships are revealed by bootstrap resampling of mass spectral pairs with SpecReBoot"

### In-depth analysis of aerucyclamide spectral connections

To further understand how aerucyclamide spectral connections are successfully recovered by our confidence-aware framework, we first correlated the presence/absence of fragment bins across bootstrap replicates with cumulative changes in the edge support values for these connections. This allowed us to prioritize fragment peaks important for these links, which could represent conserved structural motifs in these molecules. Notably, due to the metric-agnostic nature of SpecReBoot, these correlations can be computed for different spectral similarity measures, whether cosine-based or ML-based. For instance, it is evident that modified cosine and MS2DeepScore weight fragment peaks differently when drawing spectral connections. Ultimately, these fragments can be structurally annotated, increasing our understanding of the recovered connections in the obtained molecular network.

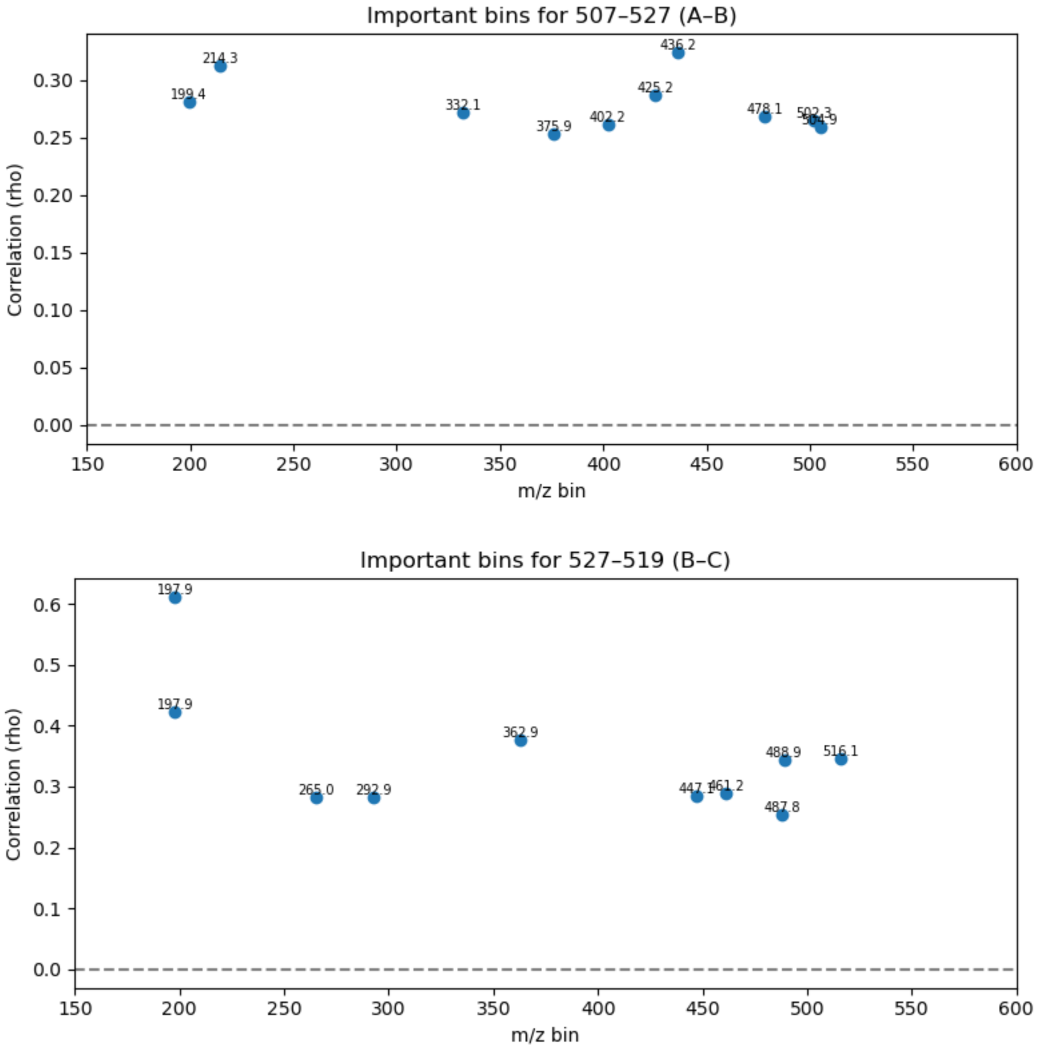

**Fig. S1.** Correlation analysis of resampled *m/z* bins underlying bootstrap-derived spectral relationships among aerucyclamides A–C. For each pairwise comparison (A–B, top; B–C, bottom), *m/z* bins showing the strongest positive correlation with changes in edge support across bootstrap replicates are shown. These bins correspond to fragment ions whose repeated presence stabilizes the corresponding spectral relationships, highlighting fragment-level features that drive the recovery of aerucyclamide connections under SpecReBoot despite low conventional spectral similarity.

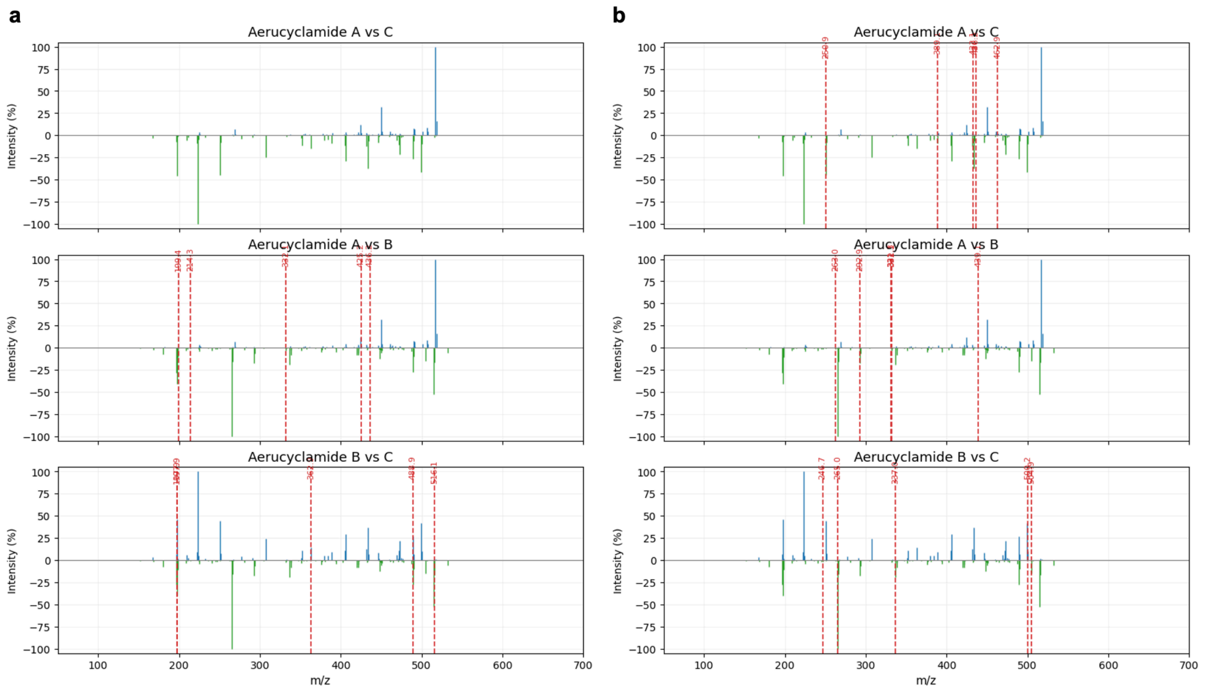

**Fig. S2.** Fragment-level interpretation of bootstrap-supported spectral relationships among aerucyclamides A–C, computed using modified cosine similarity (a) and MS2DeepScore (b). Mirror plots show pairwise MS/MS spectral comparisons for aerucyclamide A vs C (top), A vs B (middle), and B vs C (bottom). Red dashed lines indicate *m/z* bins identified as informative by SpecReBoot, corresponding to fragment bins whose resampling frequency correlates positively with increases in consensus similarity and edge support across bootstrap replicates.

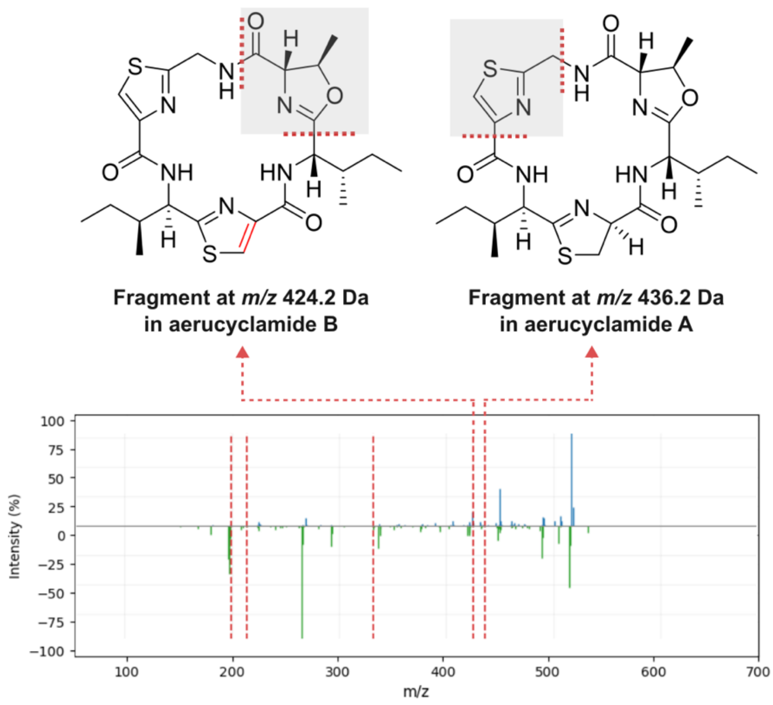

**Fig. S3.** Annotated MS/MS main fragments correlated with the high edge support observed between aerucyclamides A and B. Top panel shows structural assignments of characteristic fragment ions at *m/z* 424.2 (aerucyclamide B) and *m/z* 436.2 (aerucyclamide A), corresponding to conserved backbone substructures. The bottom panel shows the mirror plot of the corresponding MS/MS spectra, with dashed red lines indicating the recurrent fragment ions that consistently contribute to the bootstrap-supported spectral connection.

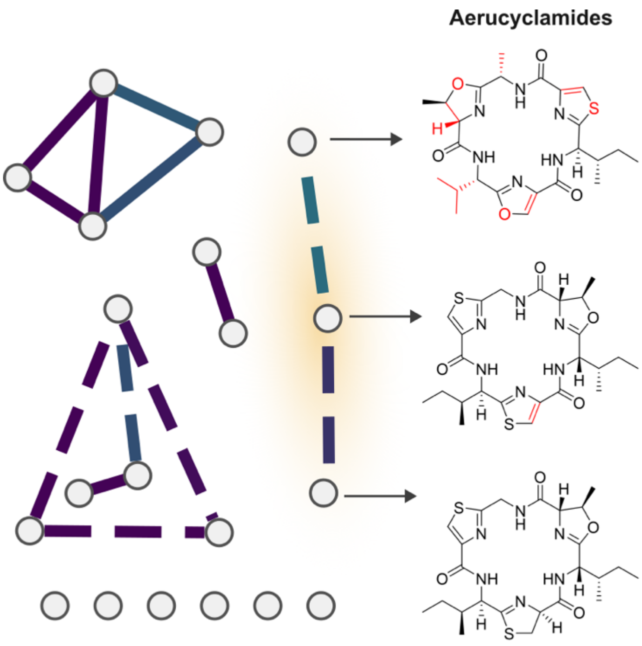

**Fig. S4.** Molecular network reconstructed using bootstrap-derived edge support for RiPP-associate spectra. Solid edges denote core connections identified by conventional similarity-based networking, whereas dashed edges correspond to relationships rescued by SpecReBoot.

**Structure elucidation of caliensomycin**

$^1$H and HSQC NMR spectra displayed five methyl, seven methylene and eight methine groups, including four oxymethines and one olefinic methine. The $^{13}$C spectrum further indicated the presence of two ketone carbonyls, one carboxyl carbon, as well as one olefinic and one dioxygenated sp$^3$ hybridized carbons devoid bound protons. The C–1‘ to C–8‘ side chain and the C-11 to C–17 moieties were readily identified by COSY and HMBC correlations, including their mutual connection via C–11, confirming the structural similarity of **1** to phomol. However, pronounced differences relative to phomol were observed for the remaining carbon framework. A large spin system extending from $H_2$–3 to H–9 was established by COSY, TOCSY and H2BC correlations. HMBC correlations from $H_3$–1, $H_2$–3 and H–4 located one carbonyl at C–2. Additional key HMBC correlations from H–11 to C–9 and C–10 from H–9 to C–8 and C–10, from 8–OH to C–8 and C–9 as well as from H–12 to C–8, enabled the assignment of the planar structure of this compound. The relative configuration of the bicyclic core was determined by ROESY correlations. Strong ROESY cross-peaks between H–4/H–6, H–5/H–9, H–6/8–OH, 8–OH/H–12, and H–9/H–11 indicated diaxial relationships among these protons, which were further supported by large $^3J_{H,H}$ coupling constants.

**Table S1.** NMR data ($^1$H 500 MHz, $^{13}$C 125 MHz) of caliensomycin in DMSO–$d_6$.

| pos | $d_C$, mult. | $d_H$, mult. |
|---|---|---|
| 1 | 30.1, $CH_3$ | 2.03, s |
| 2 | 207.5, C | |
| 3 | 44.1, $CH_2$ | 2.41, dd (15.9, 3.4) |
| | | 2.32, dd (15.9, 6.9) |
| 4 | 34.6, CH | 2.13, m |
| 5 | 77.8, CH | 2.99, ps t (9.6) |
| 6 | 70.1, CH | 3.35, m |
| 7 | 43.4, $CH_2$ | 2.07, dd (12.7, 4.8) |
| | | 1.63, ps t (12.1) |
| 8 | 99.3, CH | OH: 6.41, br s |
| 9 | 57.1, CH | 2.86, d (11.6) |
| 10 | 200.0, C | |
| 11 | 77.1, CH | 4.88, br d (9.5) |
| 12 | 71.8, CH | 4.06, ddd (9.5, 7.3, 4.0) |
| 13 | 32.2, $CH_2$ | 1.56, m |
| 14 | 24.2, $CH_2$ | 1.41, m |
| 15 | 31.0, $CH_2$ | 1.25, m |
| 16 | 22.1, $CH_2$ | 1.26, m |
| 17 | 14.0, $CH_3$ | 0.85, m |
| 1’ | 165.9, C | |
| 2’ | 125.5, C | |
| 3’ | 148.6, CH | 6.50, dq (10.1, 1.3) |
| 4’ | 34.3, CH | 2.44, m |
| 5’ | 29.0, $CH_2$ | 1.40, m |
| | | 1.30, m |
| 6’ | 11.8, $CH_3$ | 0.82, t (7.3) |
| 7’ | 12.4, $CH_3$ | 1.79, d (1.3) |
| 8’ | 19.4, $CH_3$ | 0.97, d (6.7) |

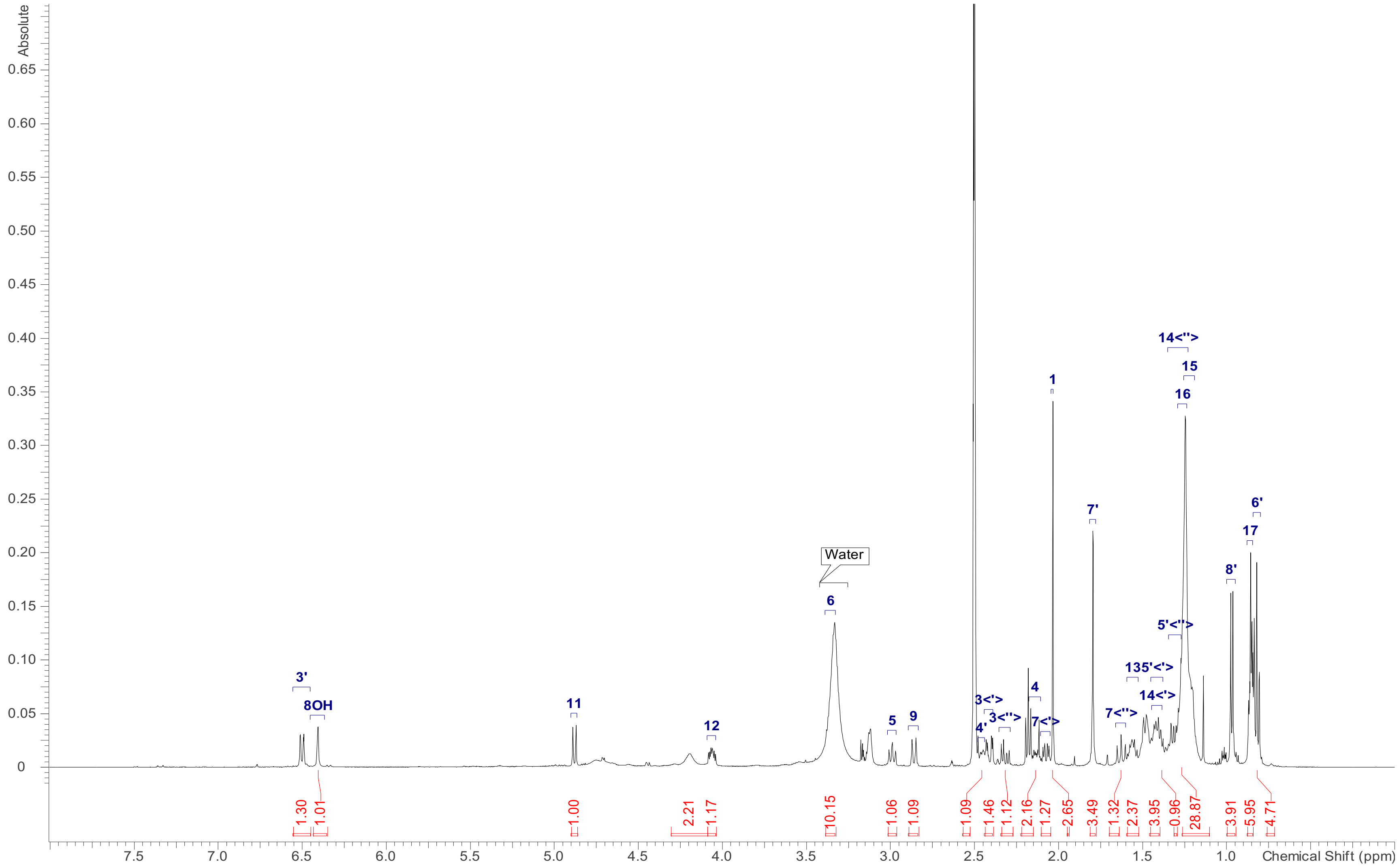

**Fig. S5.** $^{1}$H NMR (500 MHz) spectrum of caliensomycin in DMSO–$d_6$.

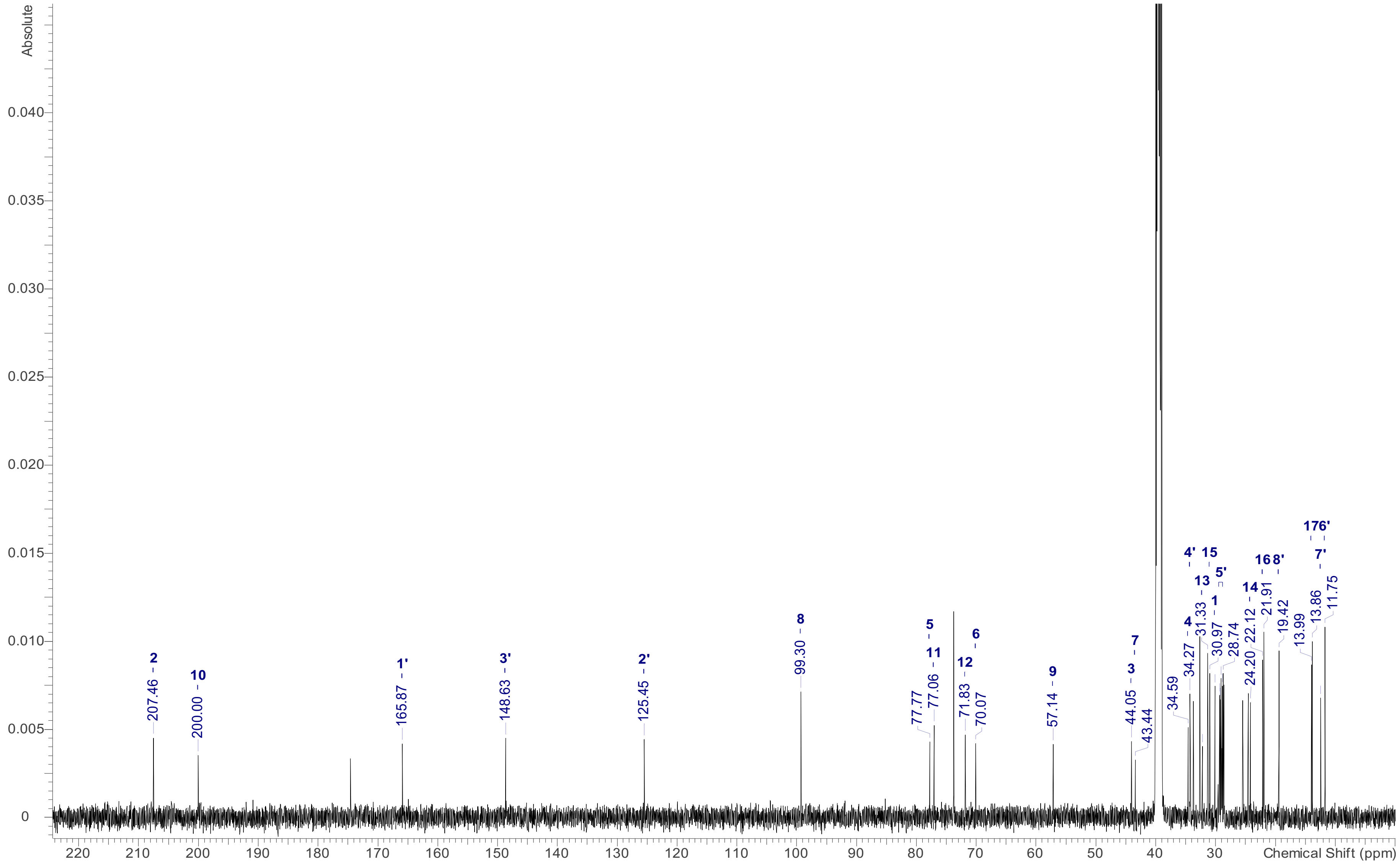

**Fig. S6.** $^{13}C$ NMR (125 MHz) spectrum of caliensomycin in DMSO–$d_6$.

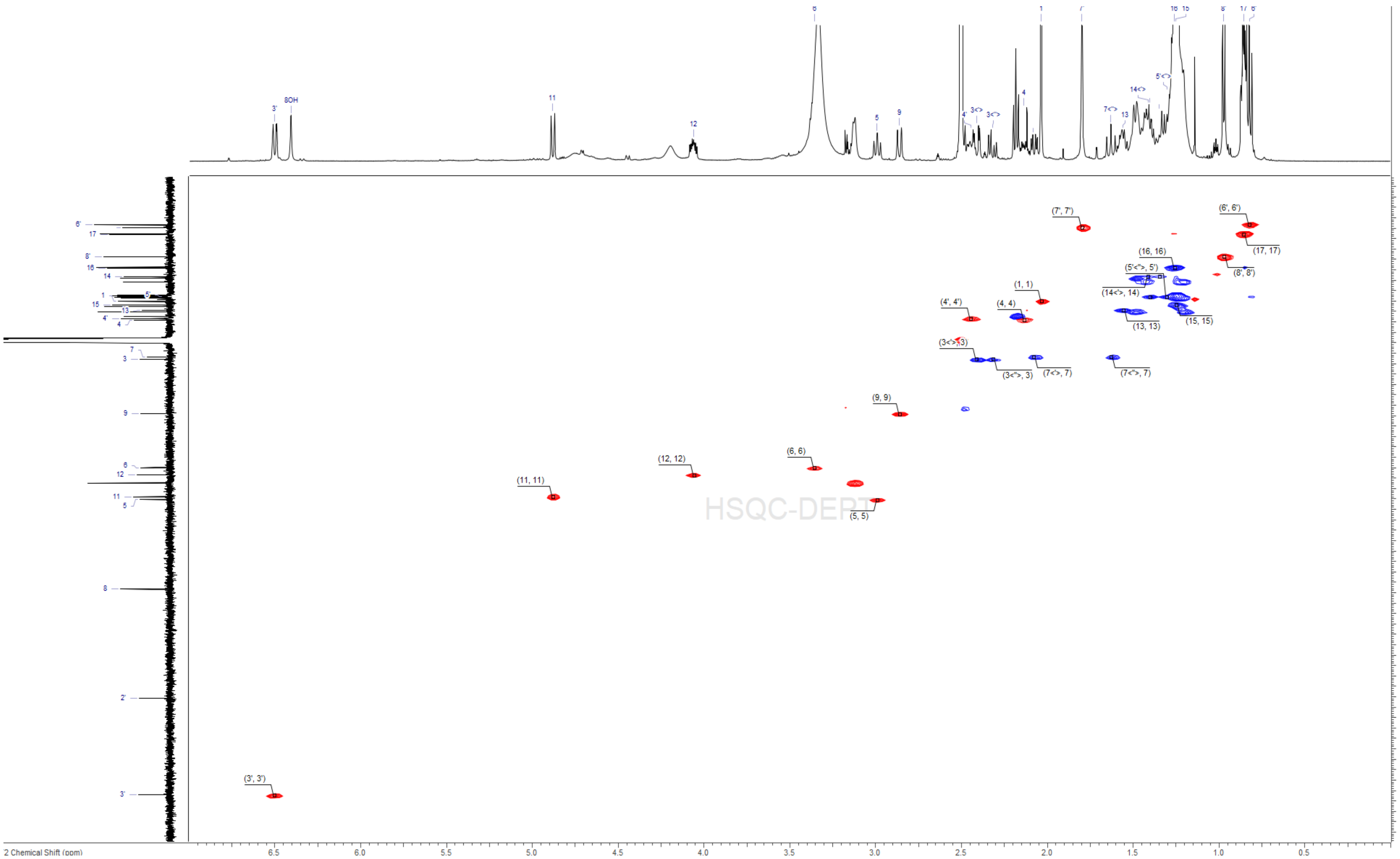

**Fig. S7.** HSQC NMR (500 MHz) spectrum of caliensomycin in DMSO–$d_6$.

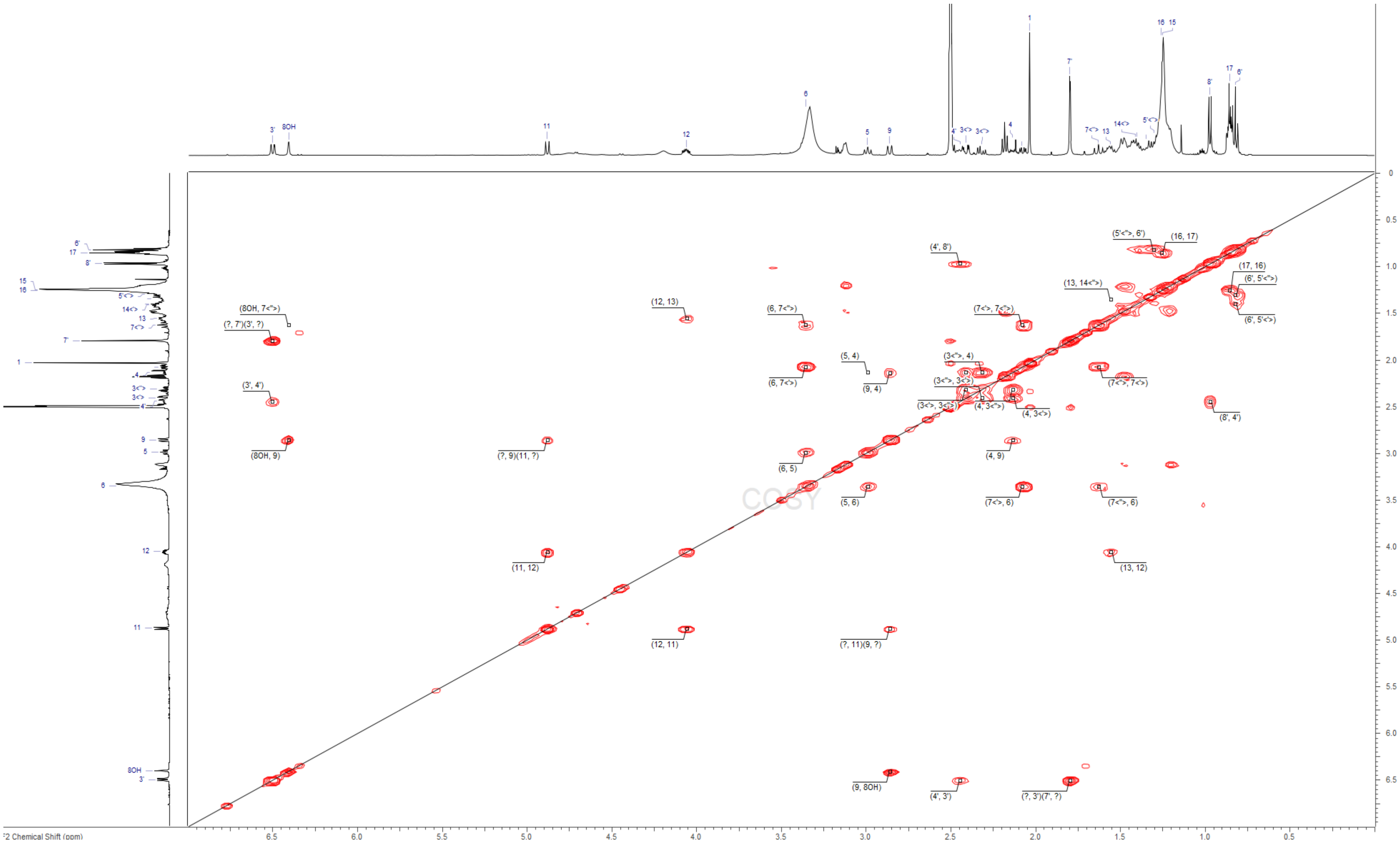

**Fig. S8.** COSY NMR (500 MHz) spectrum of caliensomycin in DMSO–$d_6$.

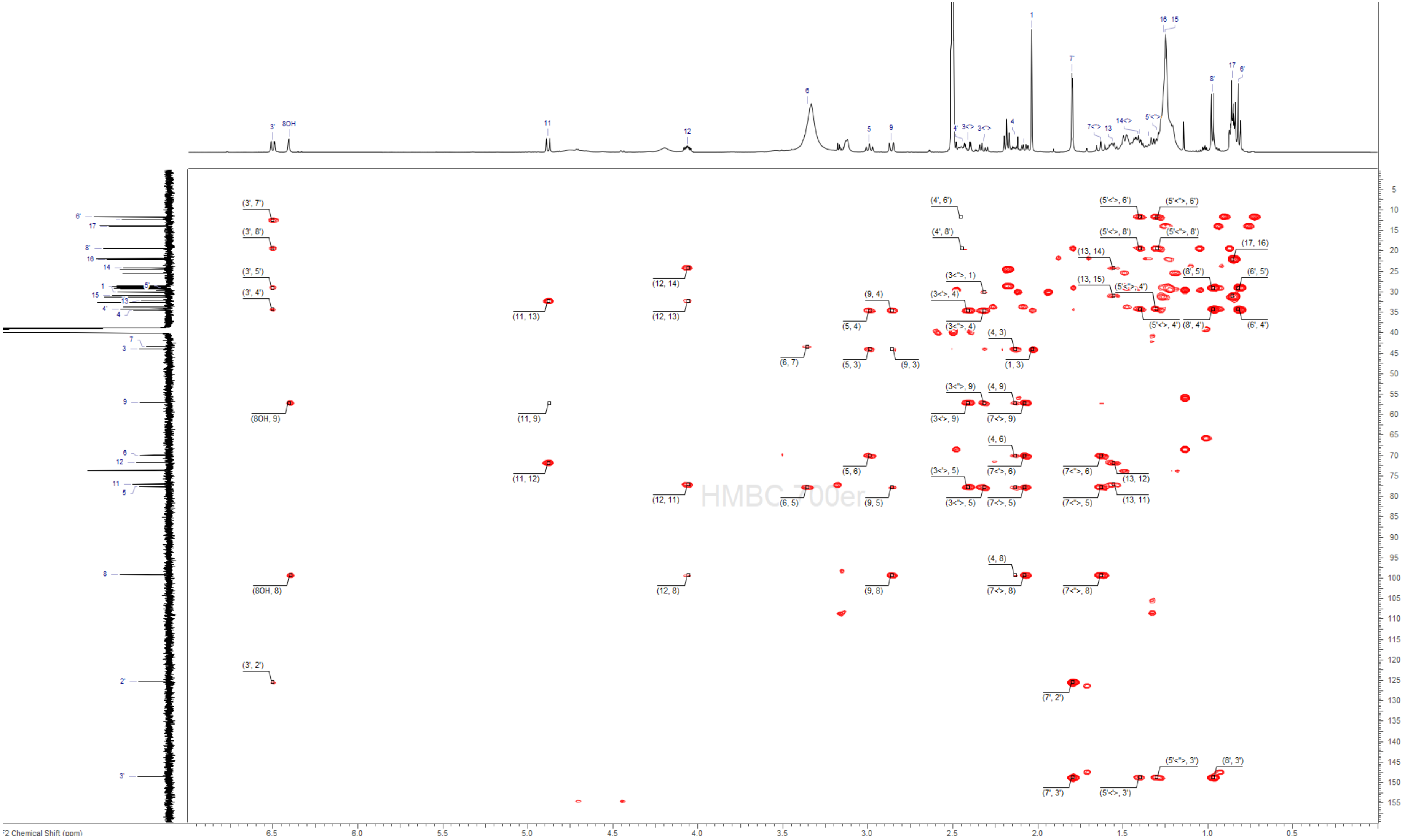

**Fig. S9.** HMBC NMR (500 MHz) spectrum of caliensomycin in DMSO–$d_6$.

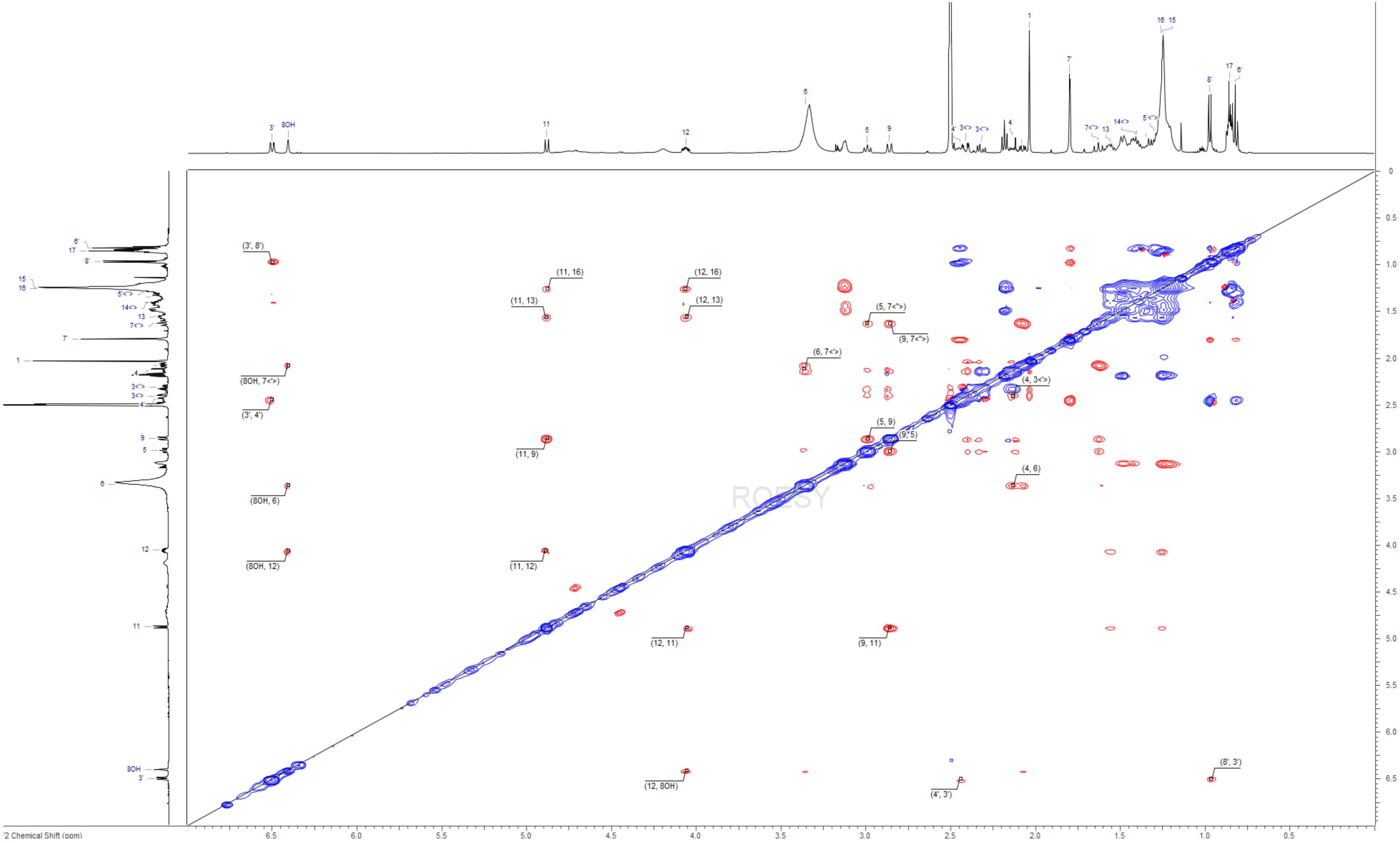

**Fig. S10.** ROESY NMR (500 MHz) spectrum of caliensomycin in DMSO–$d_6$.

### Biosynthetic characterization of phomol and caliensomycin

Synteny analysis of the candidate BGC for phomol and caliensomycin and characterized BGCs responsible for aspinolide and brefeldin A. Conserved polyketide synthases and shared tailoring enzymes indicate a common macrolactone assembly logic, supporting the assignment of the candidate cluster to the production of these compounds. Based on these insights, we propose a biosynthetic model in which phomol and caliensomycin share a common pathway involving a highly reducing PKS and cytochrome P450-mediated oxidative tailoring steps, yielding phomol as a key intermediate. Subsequent FAD-dependent oxidation drives further structural elaboration toward caliensomycin, although the precise sequence of the remaining oxidative chemical steps is yet to be elucidated.

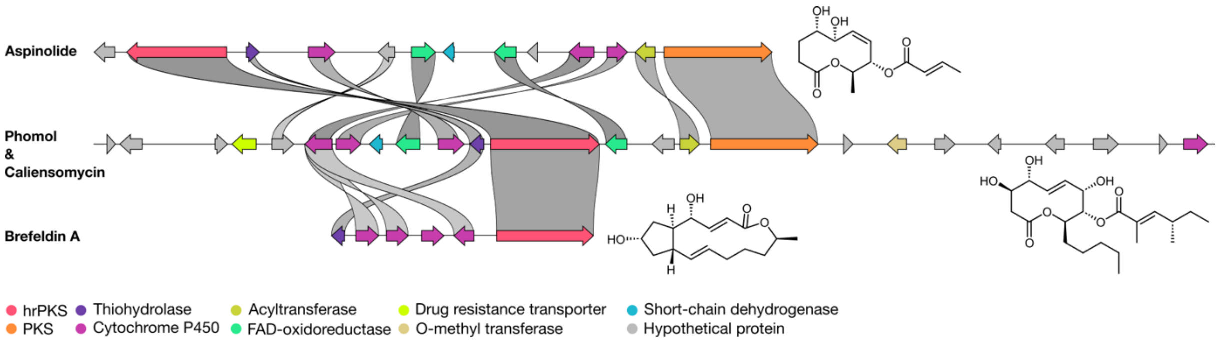

**Fig. S11.** Synteny analysis of the candidate BGC for phomol and caliensomycin in *Diaporthe caliensis*, shown in comparison with the characterized biosynthetic gene clusters responsible for aspinolide and brefeldin A production. Conserved polyketide synthases and shared tailoring enzymes highlight a common macrolactone biosynthetic logic.

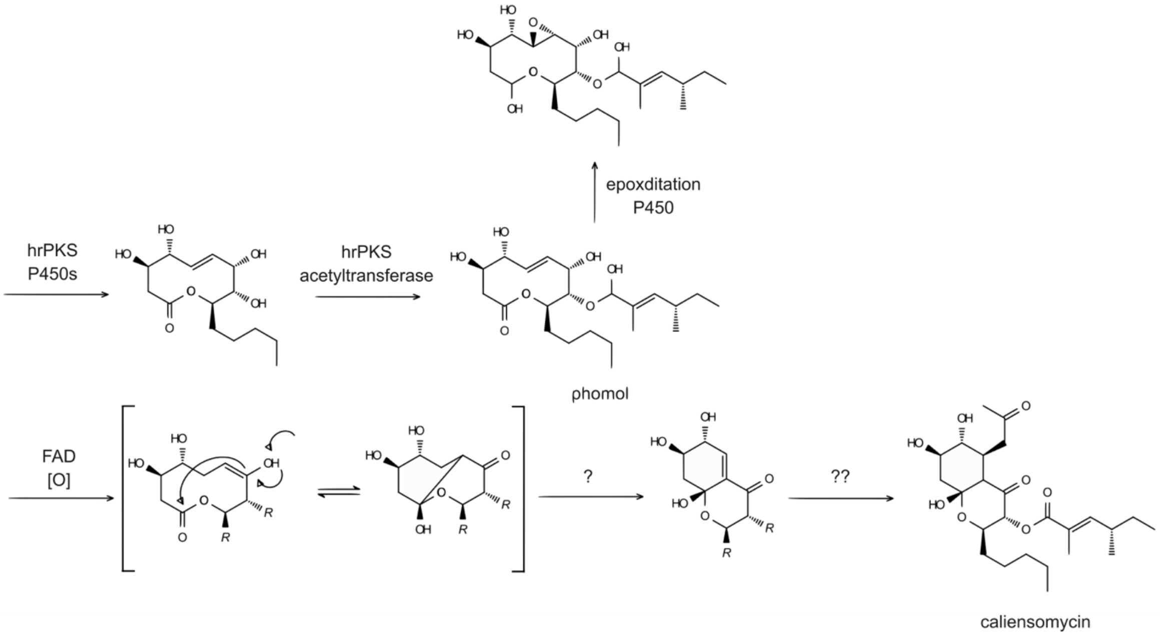

**Fig. S12.** Proposed biosynthetic model for phomol and caliensomycin. Both compounds share a common assembly route involving a highly reducing PKS (hrPKS) and cytochrome P450-mediated oxidative steps, yielding phomol as an intermediate. Subsequent late oxidative steps drive further tailoring toward caliensomycin, although the precise sequence of the remaining steps has not yet been fully elucidated.

#### Evaluation of bootstrap edge support as a chemical similarity predictor

To assess the performance of bootstrap-derived edge support as an independent chemical predictor, we evaluated the ability of three scoring schemes to discriminate chemically related compound pairs (Tanimoto similarity ≥ 0.7) using receiver operating characteristic (ROC) and precision-recall (PR) curves. The three schemes compared were: spectral similarity alone (spec_sim), bootstrap edge support alone (support), and their multiplicative combination (product). This analysis was performed across all evaluated spectral similarity metrics: cosine, modified cosine, Spec2Vec, and MS2DeepScore, and on the NIH Natural Products Library and the MSn-COCONUT datasets. In terms of overall discriminative power (ROC-AUC), spectral similarity alone performs best across all metrics and datasets. However, in the precision-recall space, edge support and its combination with spectral similarity performs very similar to the spectral similarity alone.

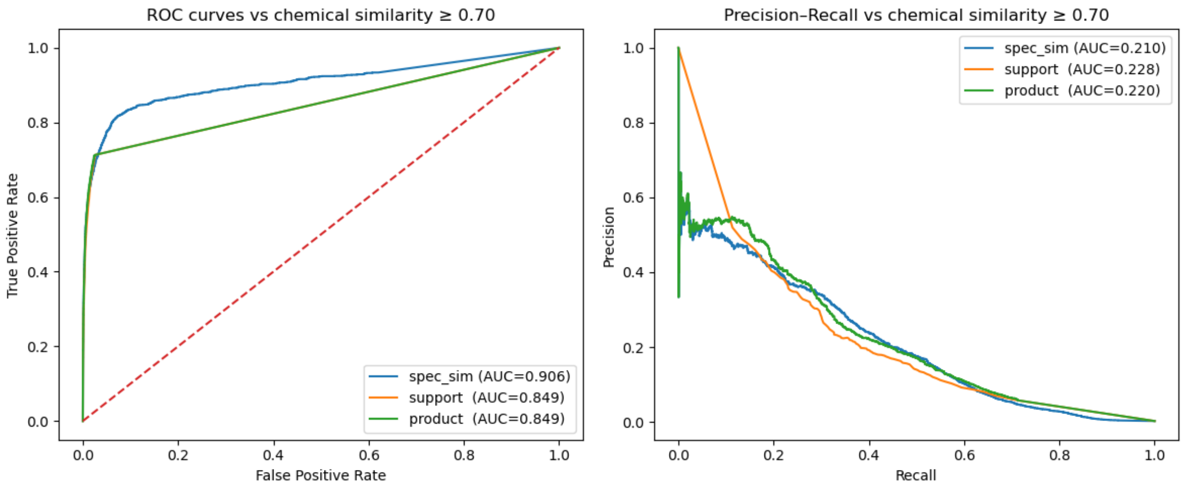

**Fig. S13.** Receiver operating characteristic (ROC) and precision–recall (PR) curves for cosine spectral similarity on the NIH Natural Products Library using a chemical similarity threshold of ≥ 0.7. Performance is shown for spectral similarity alone, (spec_sim) bootstrap-derived edge support alone (support), and their multiplicative combination (product).

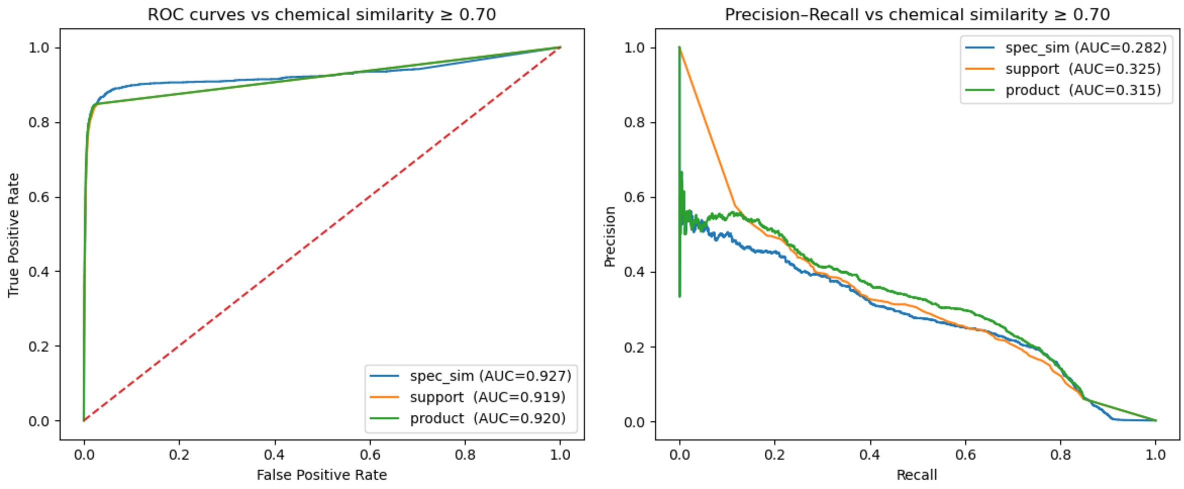

**Fig. S14.** Receiver operating characteristic (ROC) and precision–recall (PR) curves for modified cosine spectral similarity on the NIH Natural Products Library using a chemical similarity threshold of ≥ 0.7. Performance is shown for spectral similarity alone, (spec_sim) bootstrap-derived edge support alone (support), and their multiplicative combination (product).

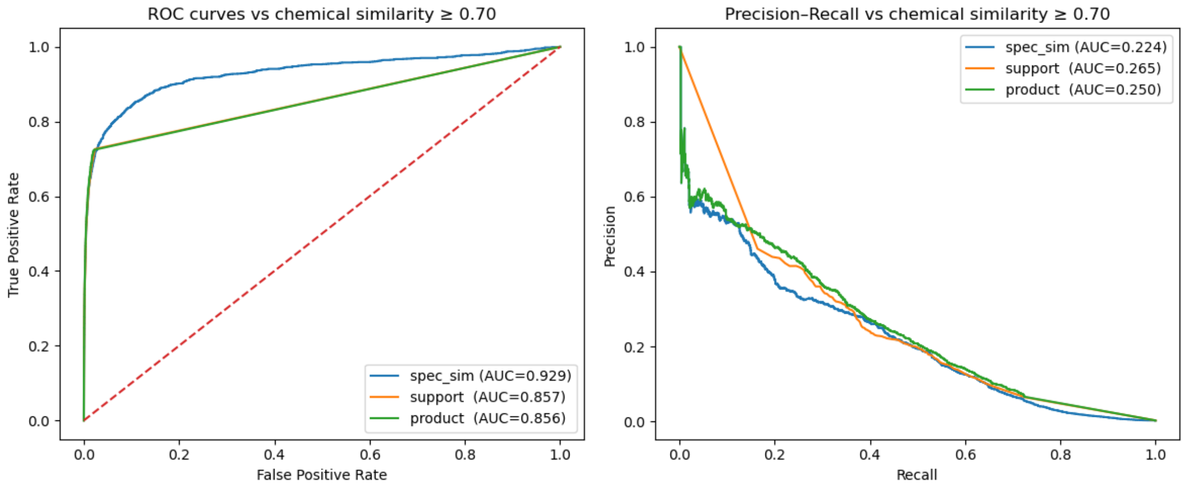

**Fig. S15.** Receiver operating characteristic (ROC) and precision–recall (PR) curves for Spec2Vec spectral similarity on the NIH Natural Products Library using a chemical similarity threshold of ≥ 0.7. Performance is shown for spectral similarity alone, (spec_sim) bootstrap-derived edge support alone (support), and their multiplicative combination (product).

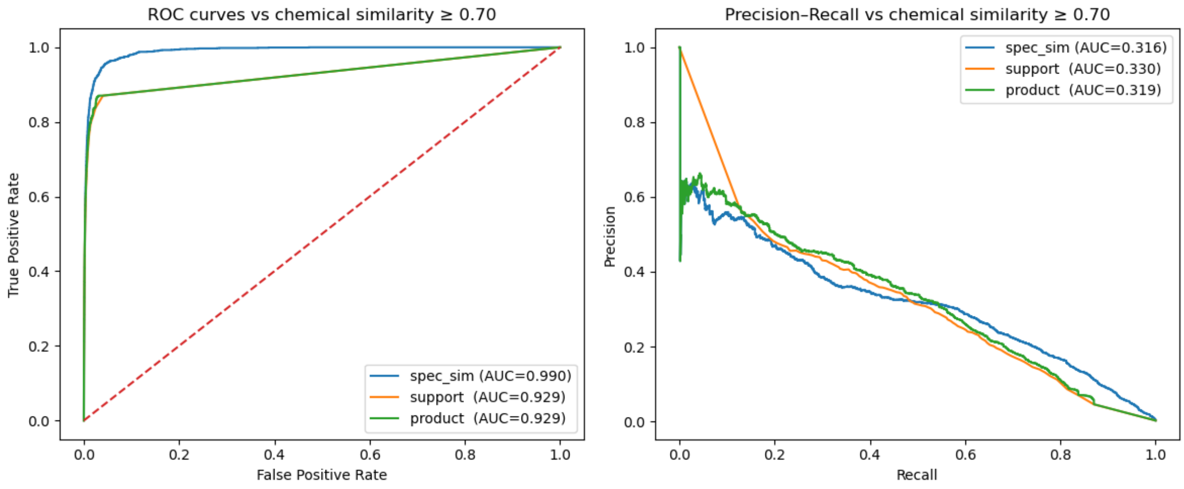

**Fig. S16.** Receiver operating characteristic (ROC) and precision–recall (PR) curves for MS2DeepScore spectral similarity on the NIH Natural Products Library using a chemical similarity threshold of ≥ 0.7. Performance is shown for spectral similarity alone, (spec_sim) bootstrap-derived edge support alone (support), and their multiplicative combination (product).

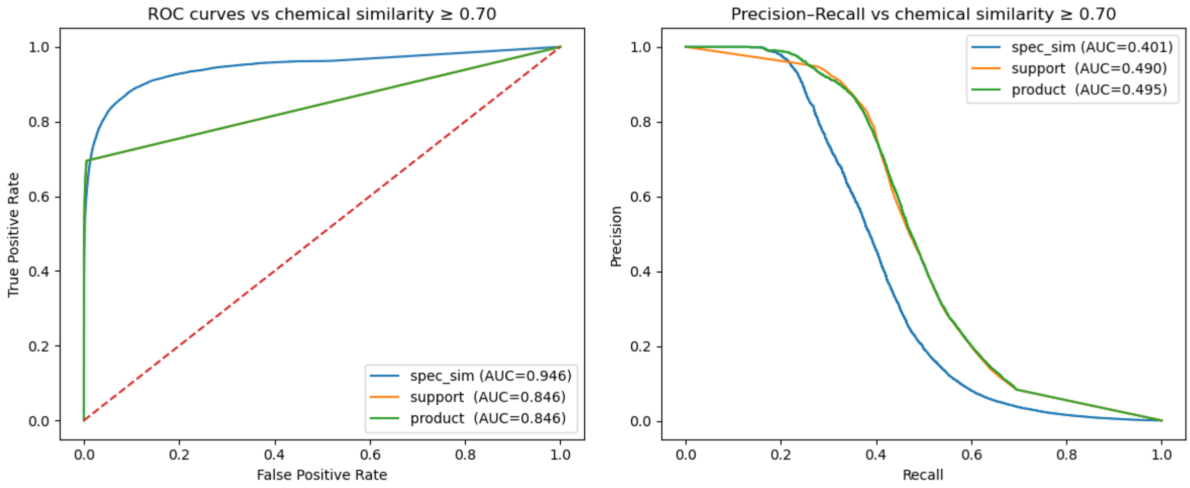

**Fig. S17.** Receiver operating characteristic (ROC) and precision–recall (PR) curves for flash cosine spectral similarity on the MSn-COCONUT library using a chemical similarity threshold of ≥ 0.7. Performance is shown for spectral similarity alone, (spec_sim) bootstrap-derived edge support alone (support), and their multiplicative combination (product).

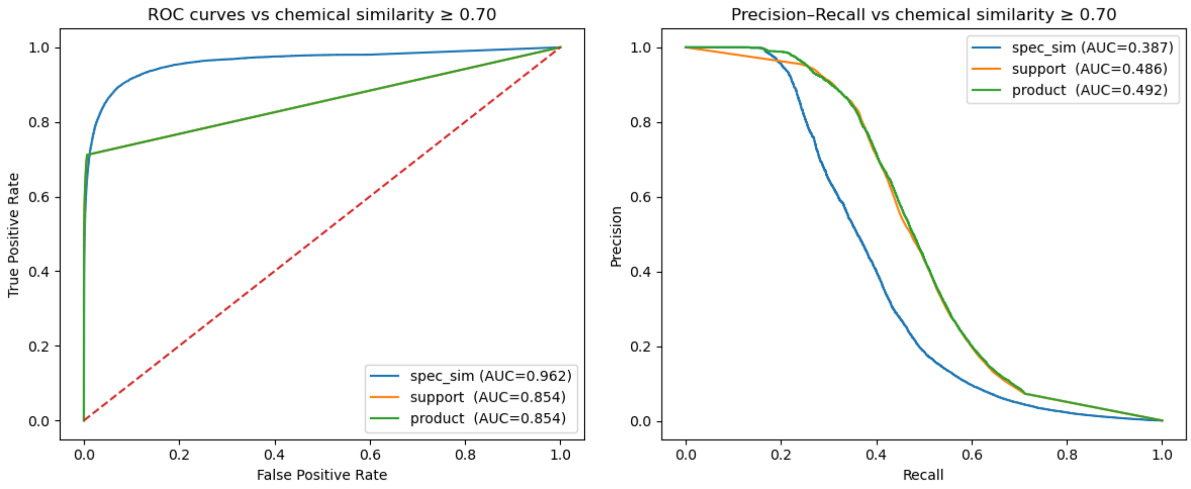

**Fig. S18.** Receiver operating characteristic (ROC) and precision–recall (PR) curves for flash modified cosine spectral similarity on the MSn-COCONUT library using a chemical similarity threshold of ≥ 0.7. Performance is shown for spectral similarity alone, (spec_sim) bootstrap-derived edge support alone (support), and their multiplicative combination (product).

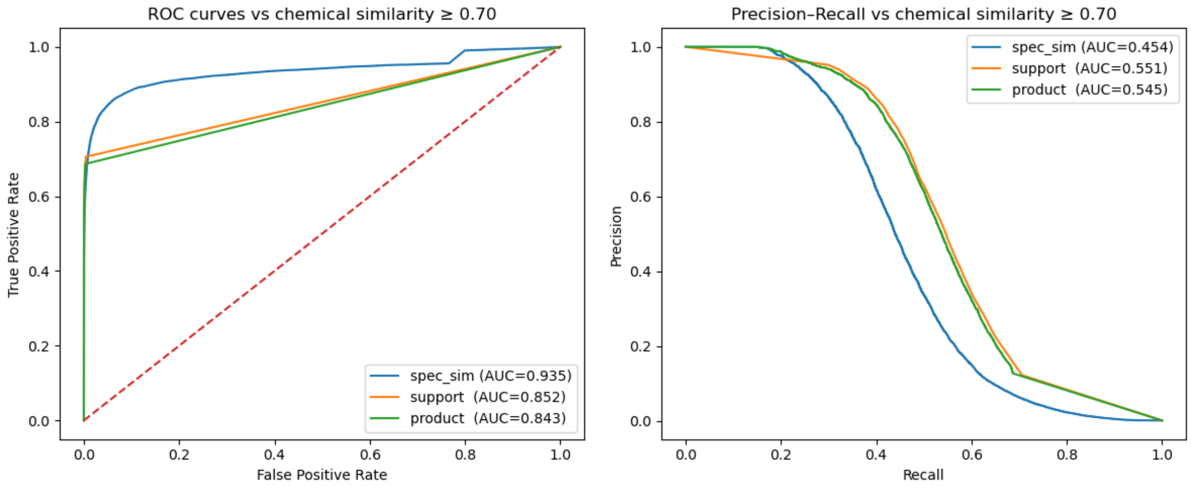

**Fig. S19.** Receiver operating characteristic (ROC) and precision–recall (PR) curves for Spec2Vec spectral similarity on the MSn-COCONUT library using a chemical similarity threshold of ≥ 0.7. Performance is shown for spectral similarity alone, (spec_sim) bootstrap-derived edge support alone (support), and their multiplicative combination (product).

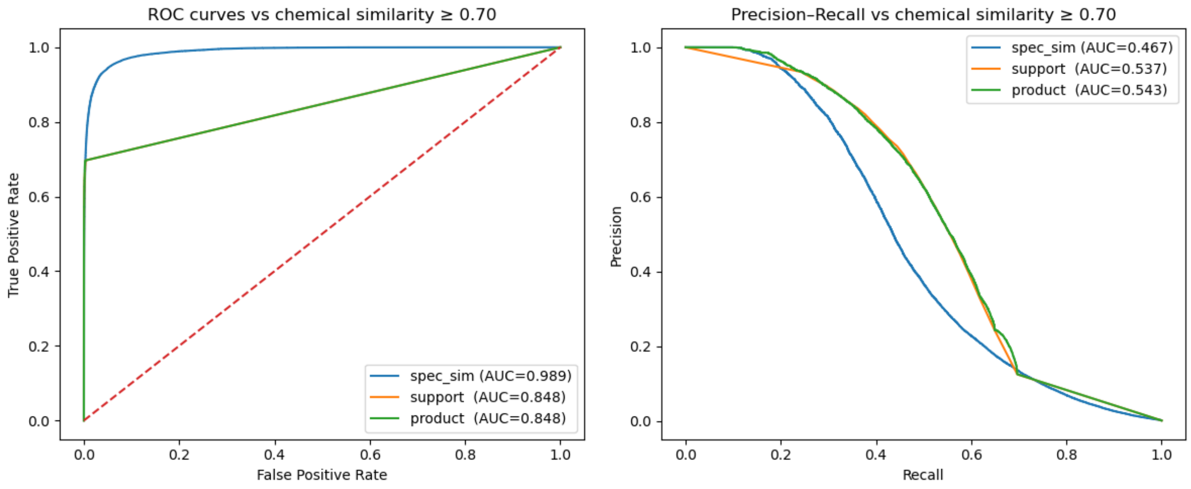

**Fig. S20.** Receiver operating characteristic (ROC) and precision–recall (PR) curves for MS2DeepScore spectral similarity on the MSn-COCONUT library using a chemical similarity threshold of ≥ 0.7. Performance is shown for spectral similarity alone, (spec_sim) bootstrap-derived edge support alone (support), and their multiplicative combination (product).

**Joint distribution of spectral and chemical similarity across metrics and datasets**

We examined the joint distribution of spectral similarity and chemical similarity (Tanimoto coefficient on Morgan fingerprints) across all pairwise spectrum comparisons, for each of the four metrics and datasets. Most pairs are concentrated at low values of both axes, reflecting the sparsity of chemically related pairs in structurally diverse datasets. A key observation across all metrics and datasets is that high spectral similarity does not reliably translate into high chemical similarity. The degree of this effect varies across metrics: ML-based methods such as MS2DeepScore result in higher similarity scores to a greater proportion of pairs overall, including chemically unrelated ones, reflecting score inflation relative to cosine-based approaches. Increasing dataset size amplifies this issue, as larger libraries encompass greater chemical scaffold diversity, making the discrimination of truly related pairs more challenging.

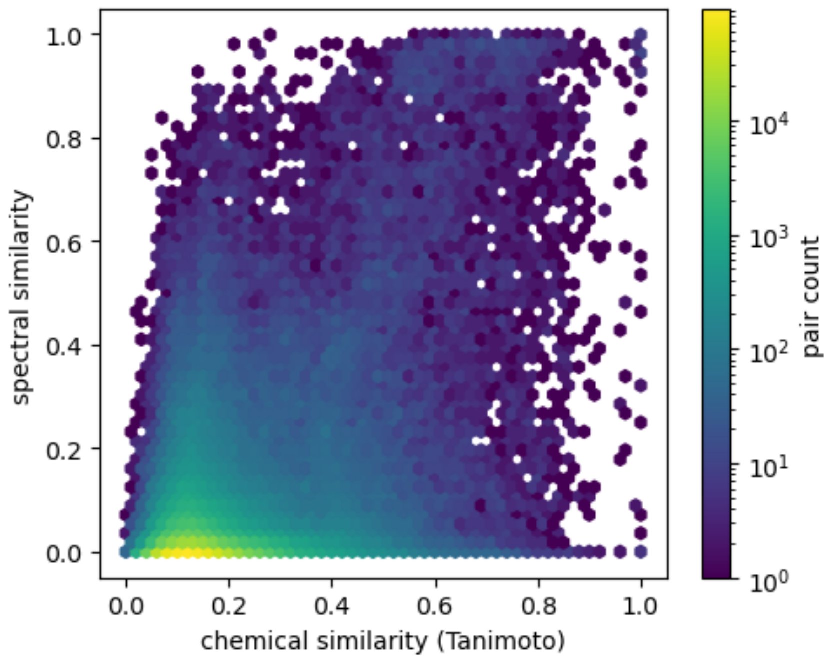

**Fig. S21.** Joint distribution of spectral similarity (cosine score) and chemical similarity (Tanimoto) for all pairwise spectral comparisons in the NIH Natural Products Library. The majority of comparisons are concentrated at low chemical similarity and low spectral similarity, reflecting the sparsity of chemically related pairs. Notably, high spectral similarity values span a broad range of chemical similarity, illustrating that cosine similarity alone does not reliably discriminate chemically related from unrelated spectra.

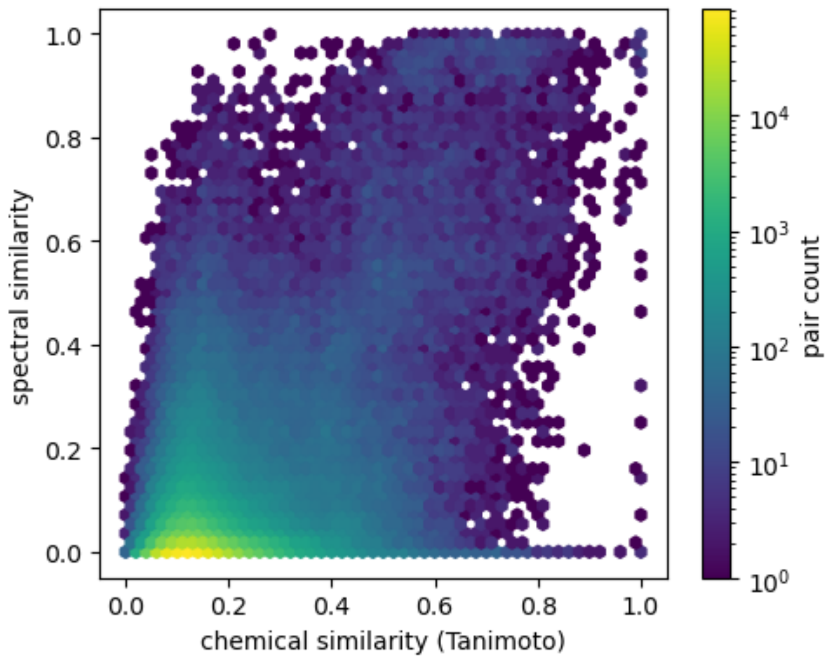

**Fig. S22.** Joint distribution of spectral similarity (modified cosine) and chemical similarity (Tanimoto) for all pairwise comparisons. Modified cosine exhibits a wide dispersion of chemical similarity values at high spectral similarity, indicating that high similarity scores do not necessarily correspond to high structural relatedness.

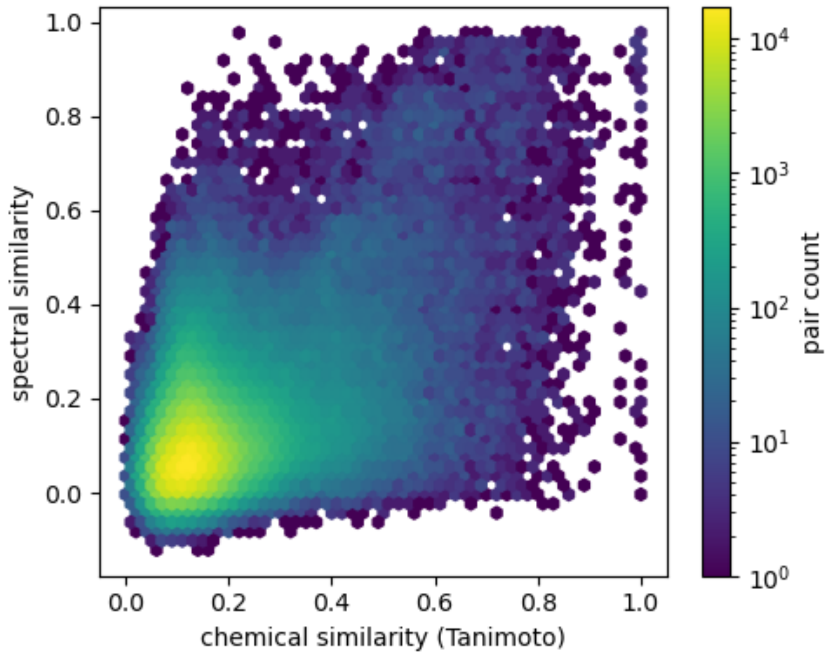

**Fig. S23.** Joint distribution of spectral similarity (Spec2Vec) and chemical similarity (Tanimoto) for all pairwise comparisons in the NIH Natural Products Library. Similar to cosine-based metrics, most comparisons cluster at low chemical similarity, indicating that structurally related pairs are rare. Although Spec2Vec exhibits a broader spread of spectral similarity values at intermediate chemical similarity, high spectral similarity values still correspond to a wide range of chemical similarity.

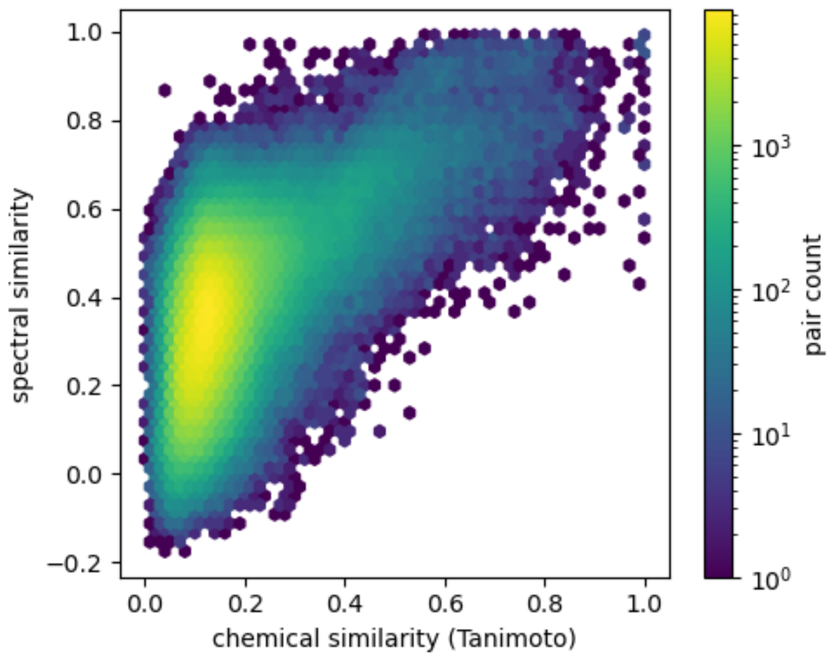

**Fig. S24.** Joint distribution of spectral similarity (MS2DeepScore) and chemical similarity (Tanimoto) for all pairwise comparisons. Compared to cosine-based metrics and Spec2Vec, MS2DeepScore shows a stronger monotonic trend between spectral and chemical similarity.

Nevertheless, a substantial fraction of high MS2DeepScore values correspond to low chemical similarity.

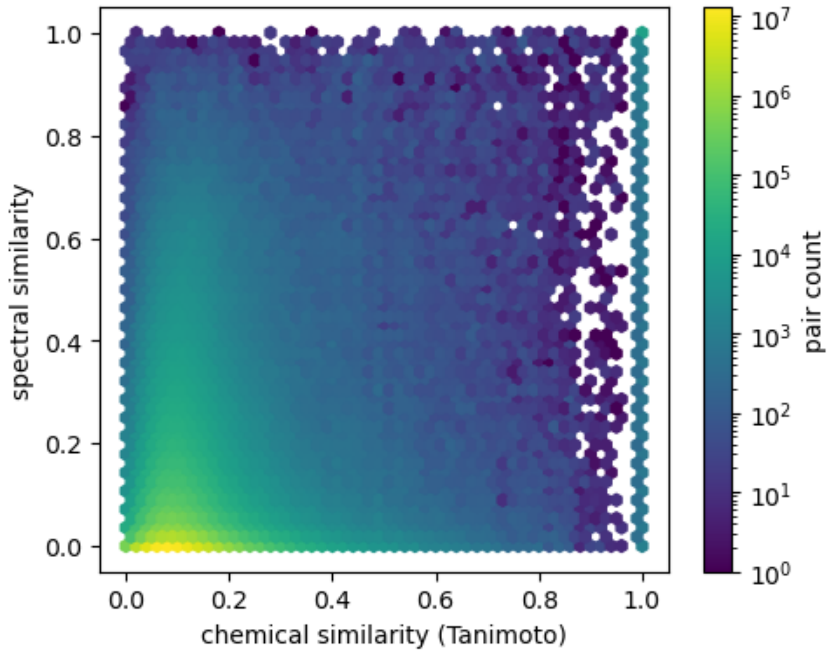

**Fig. S25.** Joint distribution of spectral similarity (flash cosine) and chemical similarity (Tanimoto) for all pairwise comparisons in the MSn-COCONNUT dataset. High flash cosine similarity values cover a broad range of chemical similarities, demonstrating that high spectral similarity does not necessarily correspond to high structural similarity.

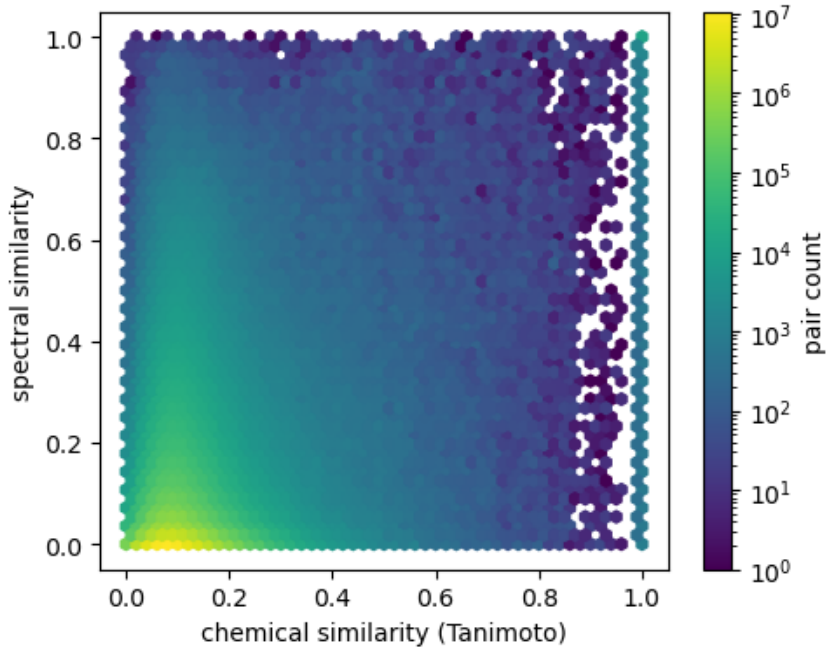

**Fig. S26.** Joint distribution of spectral similarity (flash modified cosine) and chemical similarity (Tanimoto) for all pairwise comparisons in the MSn-COCONNUT dataset.

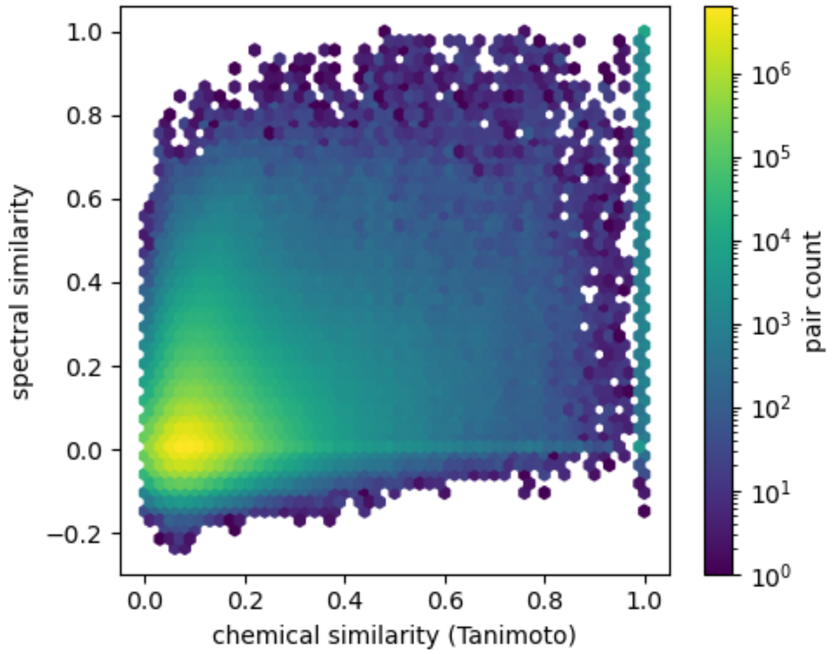

**Fig. S27.** Joint distribution of spectral similarity (Spec2Vec) and chemical similarity (Tanimoto) for all pairwise comparisons in the MSn-COCONNUT dataset.

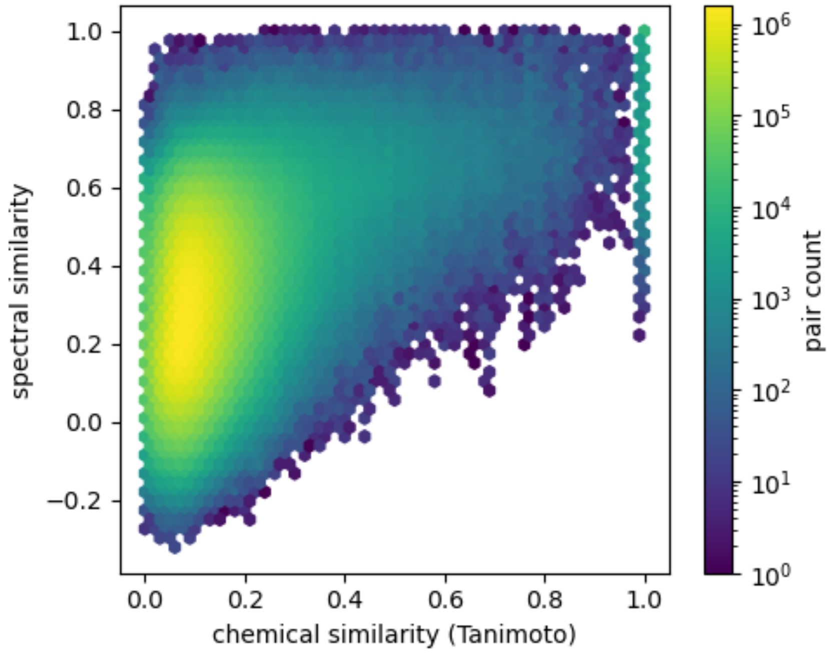

**Fig. S28.** Joint distribution of spectral similarity (MS2DeepScore) and chemical similarity (Tanimoto) for all pairwise comparisons in the MSn-COCONNUT dataset.

### Effect of bootstrap edge support filtering on chemically meaningful edge retention

To directly visualize the effect of bootstrap-derived edge support, we compared the distribution of retained edges before and after applying our dual-filter strategy, which combines a spectral similarity threshold (≥ 0.7) with an additional edge support threshold (≥ 0.5), across all four spectral similarity metrics and both datasets. In each figure, edges retained using spectral similarity alone are shown alongside those additionally filtered by the dual-filter strategy, with points colored by their edge support value. Across all metrics and datasets, application of the edge support threshold substantially reduces the number of retained edges while preferentially removing pairs with low chemical similarity.

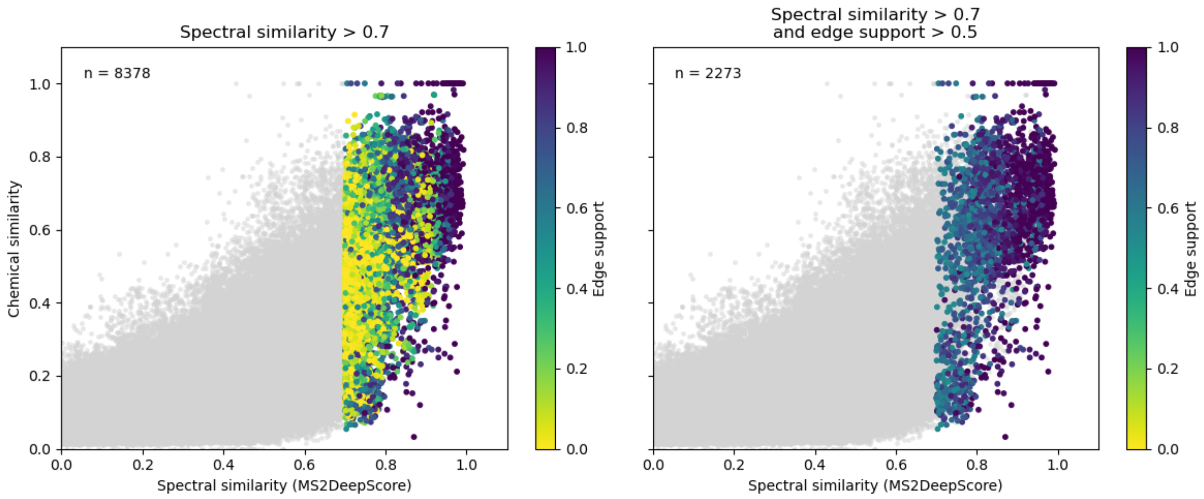

**Fig. S29.** Spectral versus chemical similarity for MS2DeepScore before and after application of bootstrap-derived edge support filtering for the NIH Natural Products Library dataset. Left, edges retained using a spectral similarity threshold alone (MS2DeepScore ≥ 0.7). Right, edges retained after applying an additional edge support threshold (≥ 0.5). Points are colored by edge support. Incorporation of edge support substantially reduces the number of retained edges (from $n$ = 8378 to $n$ = 2273) and preferentially removes edges with low chemical similarity.

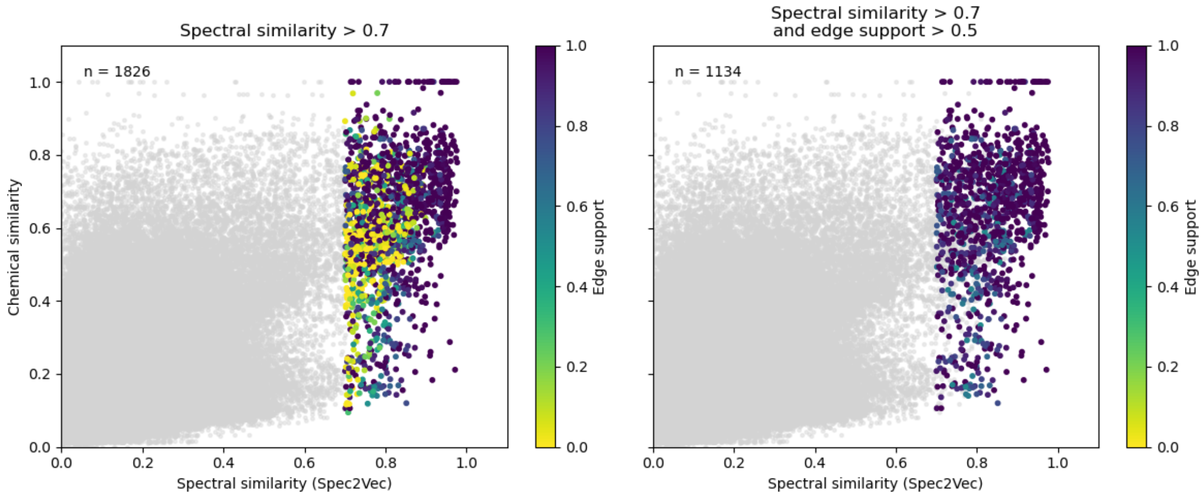

**Fig. S30.** Spectral versus chemical similarity for Spec2Vec before and after application of bootstrap-derived edge support filtering. Left, edges retained using a spectral similarity threshold alone (Spec2Vec ≥ 0.7). Right, edges retained after applying an additional edge support threshold

(≥ 0.5). Points are colored by edge support. Although Spec2Vec produces fewer high-similarity edges than MS2DeepScore, application of edge support similarly enriches for chemically related pairs (reduction from $n$ = 1826 to $n$ = 1134).

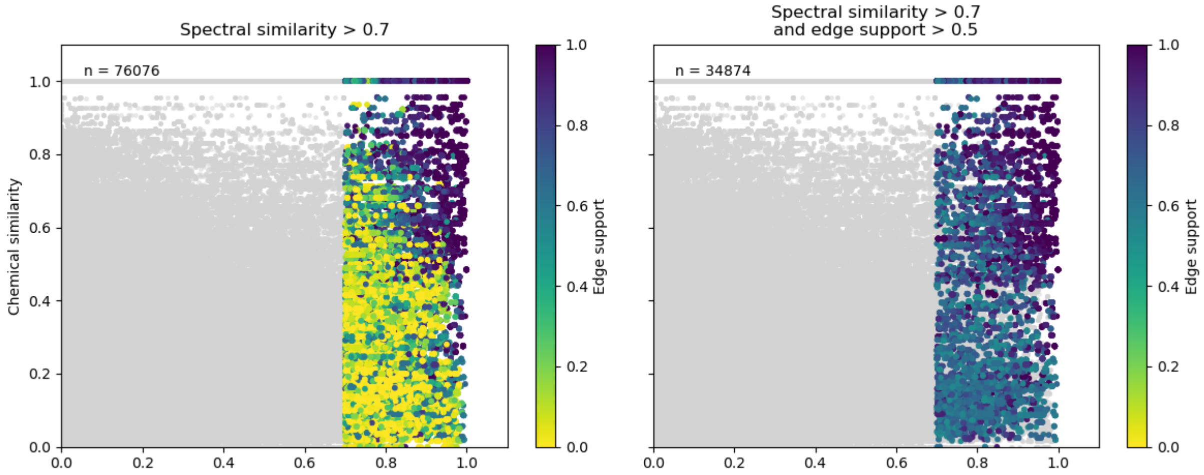

**Fig. S31.** Spectral versus chemical similarity for flash cosine before and after application of bootstrap-derived edge support filtering in the MSn-COCONUT subset. Left, edges retained using a spectral similarity threshold alone (Spec2Vec ≥ 0.7). Right, edges retained after applying an additional edge support threshold (≥ 0.5). Points are colored by edge support. Although Spec2Vec produces fewer high-similarity edges than MS2DeepScore, application of edge support similarly enriches for chemically related pairs (reduction from $n$ = 76076 to $n$ = 34874).

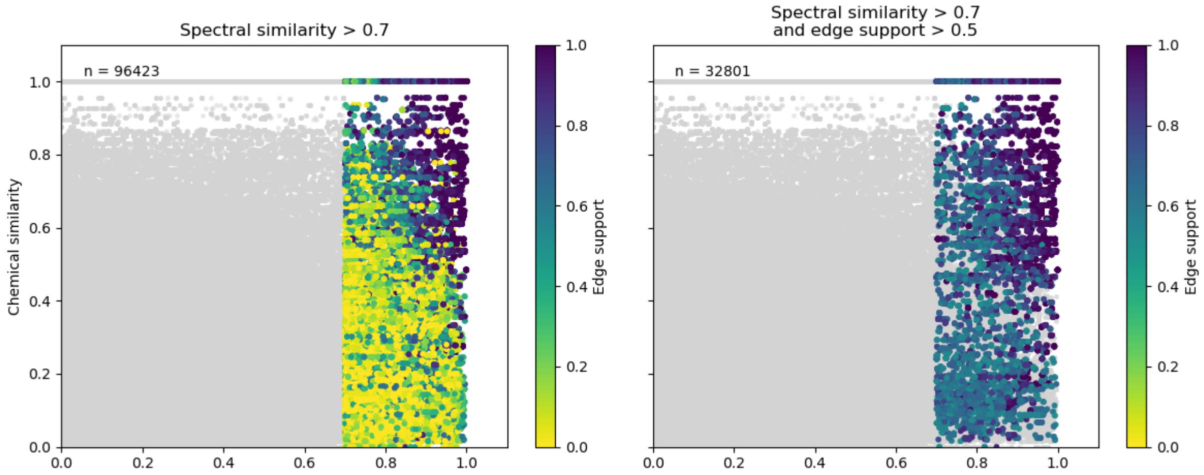

**Fig. S32.** Spectral versus chemical similarity for flash modified cosine before and after application of bootstrap-derived edge support filtering in the MSn-COCONUT subset. Left, edges retained using a spectral similarity threshold alone (Modified cosine ≥ 0.7). Right, edges retained after applying an additional edge support threshold (≥ 0.5). Points are colored by edge support.

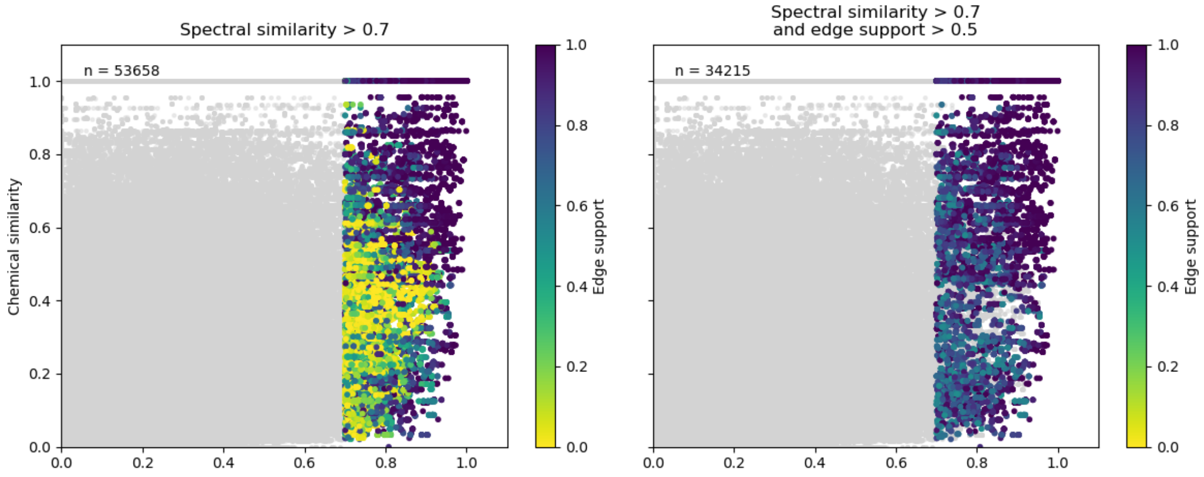

**Fig. S33.** Spectral versus chemical similarity for flash modified cosine before and after application of bootstrap-derived edge support filtering in the MSn-COCONUT subset. Left, edges retained using a spectral similarity threshold alone (Spec2Vec ≥ 0.7). Right, edges retained after applying an additional edge support threshold (≥ 0.5). Points are colored by edge support.

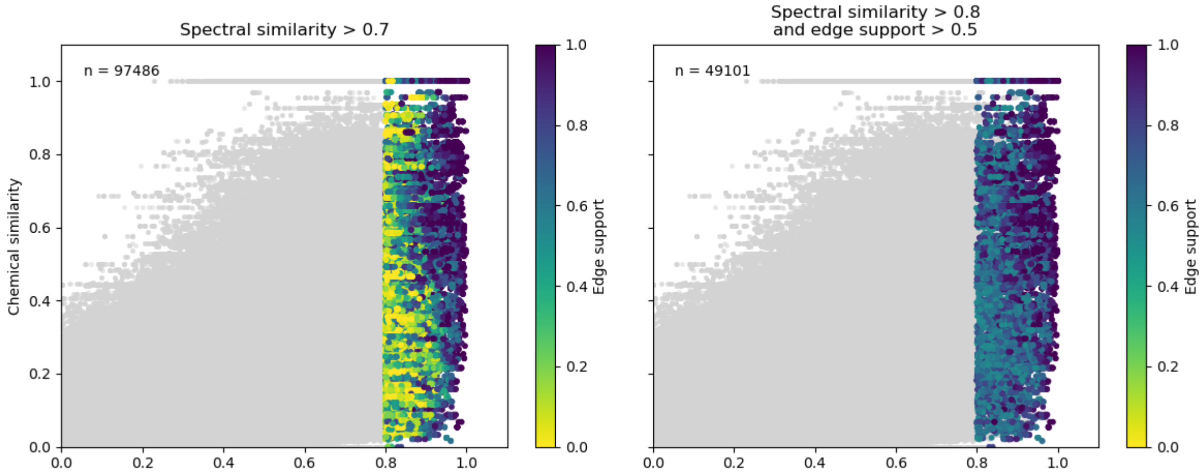

**Fig. S34.** Spectral versus chemical similarity for flash modified cosine before and after application of bootstrap-derived edge support filtering in the MSn-COCONUT subset. Left, edges retained using a spectral similarity threshold alone (MS2DeepScore ≥ 0.8). Right, edges retained after applying an additional edge support threshold (≥ 0.5). Points are colored by edge support.

**Table S2.** Pairwise and three-way overlap statistics for edges retained using modified cosine, MS2DeepScore, and Spec2Vec under different filtering scenarios. Edges were selected using spectral similarity thresholds alone, or in combination with edge support filter (> 0.5), and optionally restricted to chemically similar pairs based on Tanimoto similarity thresholds (≥ 0.7 or ≥ 0.5). For each metric pair (A, B), the table reports the number of retained edges (|A|, |B|), their intersection (|A ∩ B|), Jaccard index, and directional containment. The final column reports the number of edges shared across all three metrics under each scenario.

| Scenarios | A | B | \|A\| | \|B\| | \|A∩B\| | Jaccard | Containment A→B | Containment B→A | Shared by all metrics |
|---|---|---|---|---|---|---|---|---|---|
| Spectral similarity treshold only | ModCos | MS2DeepScore | 2626 | 2629 | 1808 | 0.52 | 0.69 | 0.69 | 1444 |
| | ModCos | Spec2Vec | 2626 | 1826 | 1683 | 0.61 | 0.64 | 0.92 | |
| | MS2DeepScore | Spec2Vec | 2629 | 1826 | 1541 | 0.53 | 0.59 | 0.84 | |
| Spectral similarity treshold only (Tanimoto similarity > 0.7) | ModCos | MS2DeepScore | 764 | 836 | 656 | 0.69 | 0.86 | 0.78 | 515 |
| | ModCos | Spec2Vec | 764 | 548 | 531 | 0.68 | 0.70 | 0.97 | |
| | MS2DeepScore | Spec2Vec | 836 | 548 | 528 | 0.62 | 0.63 | 0.96 | |
| Spectral similarity treshold only (Tanimoto similarity > 0.5) | ModCos | MS2DeepScore | 1918 | 2235 | 1569 | 0.61 | 0.82 | 0.70 | 1254 |
| | ModCos | Spec2Vec | 1918 | 1440 | 1330 | 0.66 | 0.69 | 0.92 | |
| | MS2DeepScore | Spec2Vec | 2235 | 1440 | 1336 | 0.57 | 0.60 | 0.93 | |
| Spectral similarity treshold + Edge support (> 0.5) | ModCos | MS2DeepScore | 1597 | 1589 | 1058 | 0.50 | 0.66 | 0.67 | 783 |
| | ModCos | Spec2Vec | 1597 | 1140 | 942 | 0.52 | 0.59 | 0.83 | |
| | MS2DeepScore | Spec2Vec | 1589 | 1140 | 894 | 0.49 | 0.56 | 0.78 | |
| Spectral similarity treshold + Edge support (> 0.5) (Tanimoto similarity > 0.7) | ModCos | MS2DeepScore | 612 | 670 | 513 | 0.67 | 0.84 | 0.77 | 389 |
| | ModCos | Spec2Vec | 612 | 449 | 403 | 0.61 | 0.66 | 0.90 | |
| | MS2DeepScore | Spec2Vec | 670 | 449 | 421 | 0.60 | 0.63 | 0.94 | |
| Spectral similarity treshold + Edge support (> 0.5) (Tanimoto similarity > 0.5) | ModCos | MS2DeepScore | 1239 | 1396 | 962 | 0.58 | 0.78 | 0.69 | 711 |
| | ModCos | Spec2Vec | 1239 | 907 | 776 | 0.57 | 0.63 | 0.86 | |
| | MS2DeepScore | Spec2Vec | 1396 | 907 | 802 | 0.53 | 0.57 | 0.88 | |

### Molecular network topology before and after edge support-based rebooting

To visualize the structural impact of SpecReBoot's dual-filter strategy on molecular network topology, we compared the base molecular networks, constructed using a spectral similarity threshold alone (≥ 0.7), with the corresponding threshold networks obtained after additionally applying a bootstrap-derived edge support threshold (≥ 0.5). In the base networks, the large number of retained edges results in densely connected components that frequently incorporate weakly supported spectral connections. Application of the edge support filter substantially reduces network complexity, removing spurious connections while preserving edges that are consistently recovered across bootstrap replicates. The resulting threshold networks are more modular, with components that better reflect genuine chemical relationships.

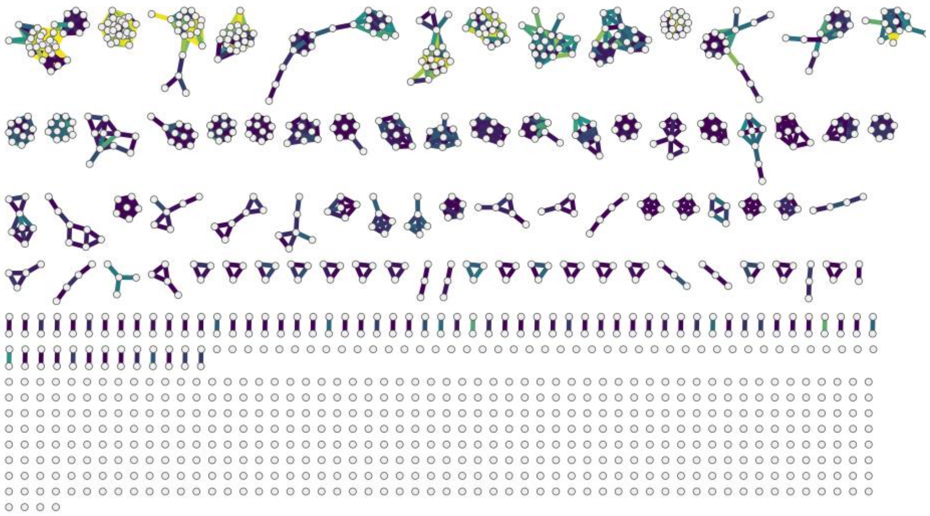

**Fig. S35.** Base molecular network of the NIH Natural Products Library constructed using the flash cosine similarity score. The network was generated using a spectral similarity threshold of 0.7 and standard topology-based parameters, without incorporating bootstrap-derived edge support. Nodes represent MS/MS spectra, and edges represent pairwise spectral similarities.

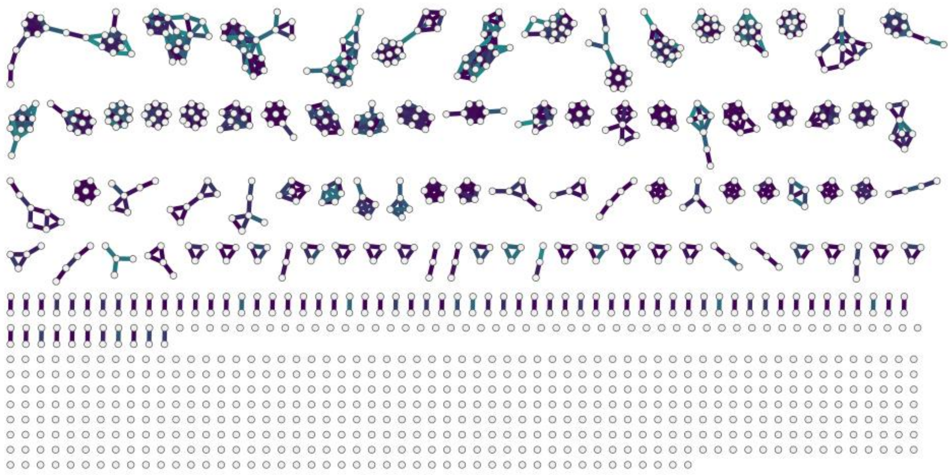

**Fig. S36.** Threshold molecular network of the NIH Natural Products Library obtained after applying SpecReBoot to the flash cosine–based molecular network. Edges shown correspond to spectrally similar connections retained after applying both the spectral similarity threshold (≥ 0.7) and a bootstrap-derived edge support filter (≥ 0.5), resulting in a reduced network enriched in stable and chemically coherent relationships.

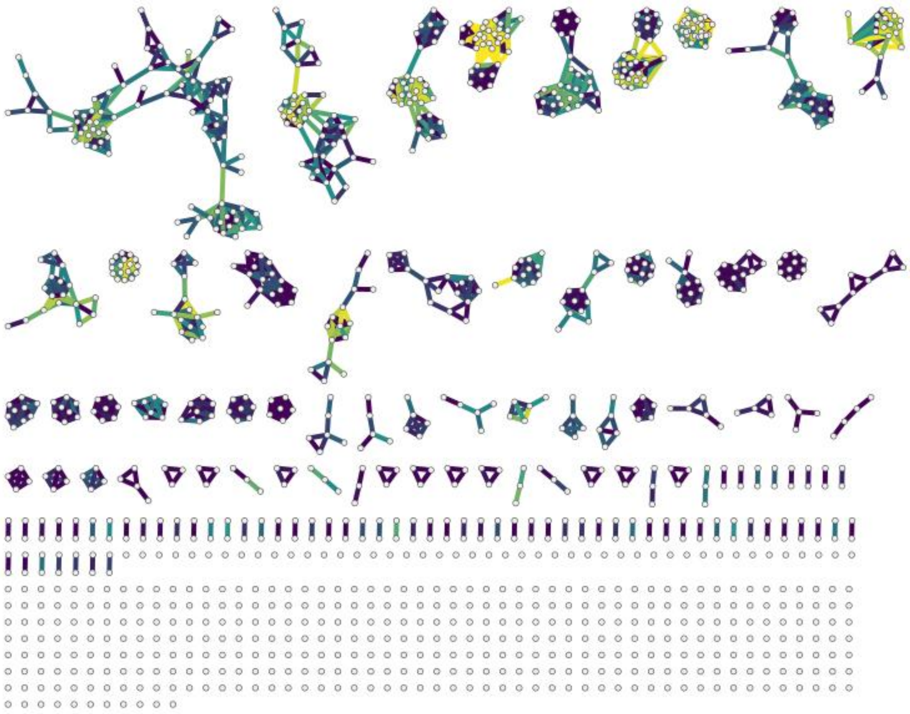

**Fig. S37.** Base molecular network of the NIH Natural Products Library constructed using the modified cosine similarity score. The network was generated using a spectral similarity threshold of 0.7 and standard topology-based parameters, without incorporating bootstrap-derived edge support. Nodes represent MS/MS spectra, and edges represent pairwise spectral similarities.

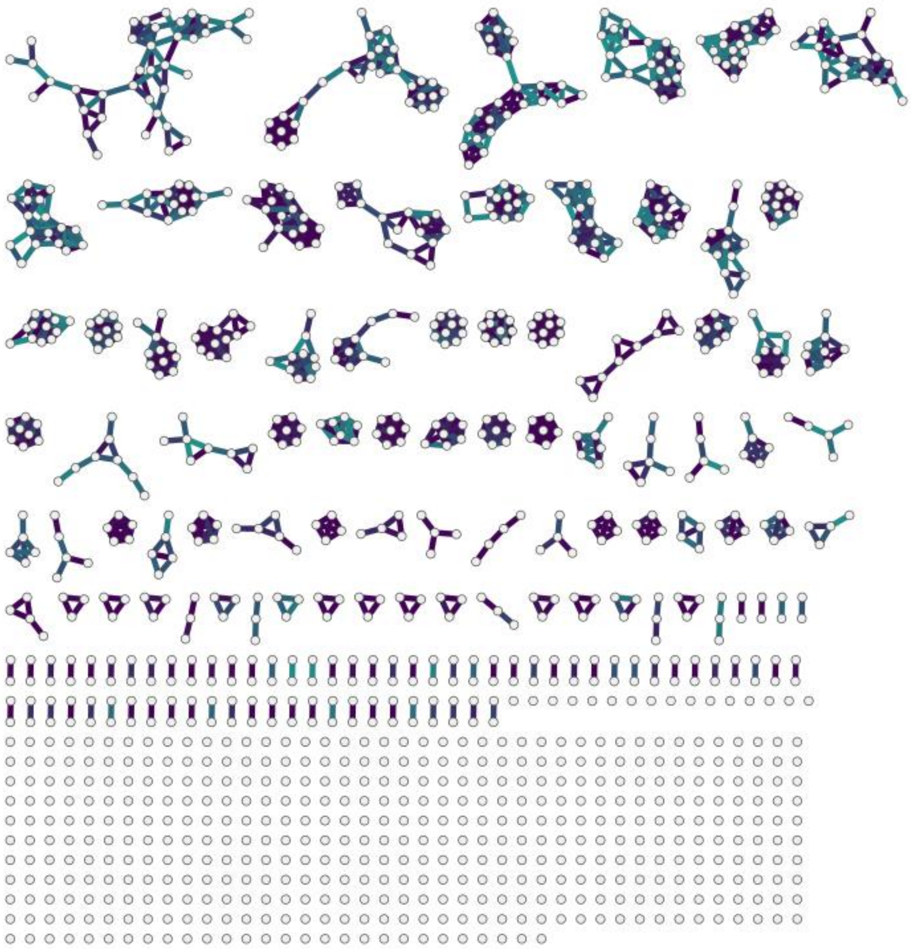

**Fig. S38.** Threshold molecular network of the NIH Natural Products Library obtained after applying SpecReBoot to the modified cosine–based molecular network. Edges shown correspond to spectrally similar connections retained after applying both the spectral similarity threshold (≥ 0.7) and a bootstrap-derived edge support filter (≥ 0.5), resulting in a reduced network enriched in stable and chemically coherent relationships.

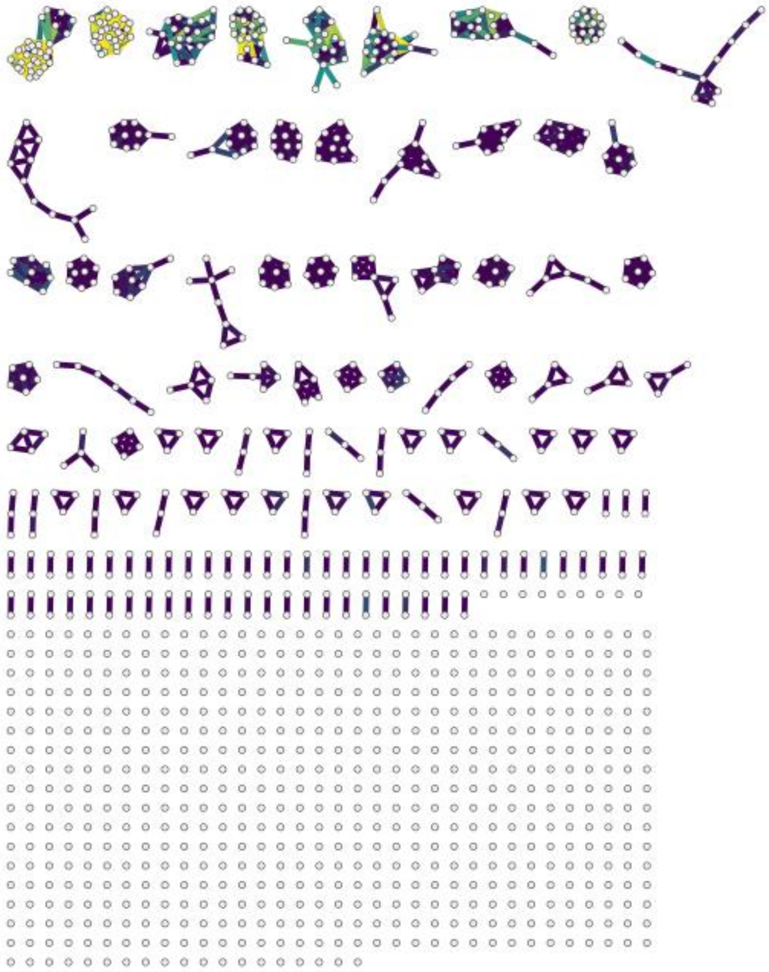

**Fig. S39.** Base molecular network of the NIH Natural Products Library constructed using the Spec2Vec similarity score. The network was generated using a spectral similarity threshold of 0.7 and topology-based parameters as for cosine-based metrics.

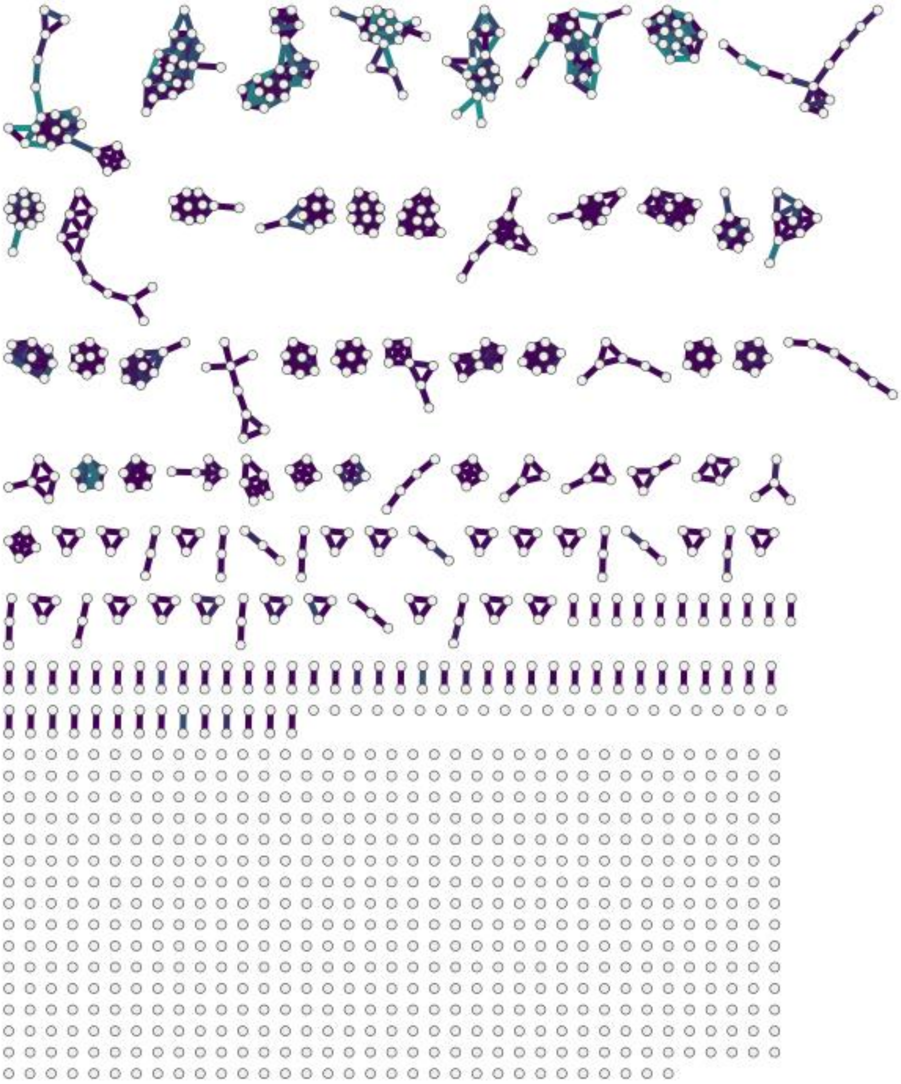

**Fig. S40.** Threshold molecular network of the NIH Natural Products Library obtained after applying SpecReBoot to the Spec2Vec–based molecular network. Edges shown correspond to spectrally similar connections retained after applying both the spectral similarity threshold (≥ 0.7) and a bootstrap-derived edge support filter (≥ 0.5), resulting in a reduced network enriched in stable and chemically coherent relationships.

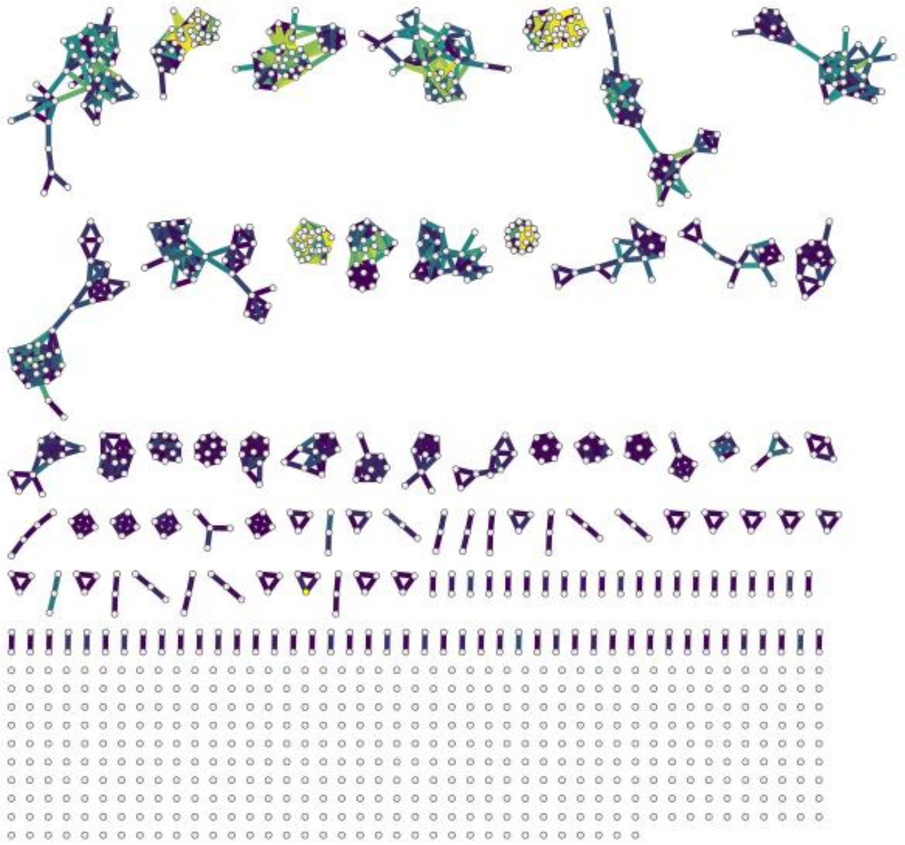

**Fig. S41.** Base molecular network of the NIH Natural Products Library constructed using the MS2Deepscore similarity score. The network was generated using a spectral similarity threshold of 0.8 and topology-based parameters as for cosine-based metrics and Spec2Vec.

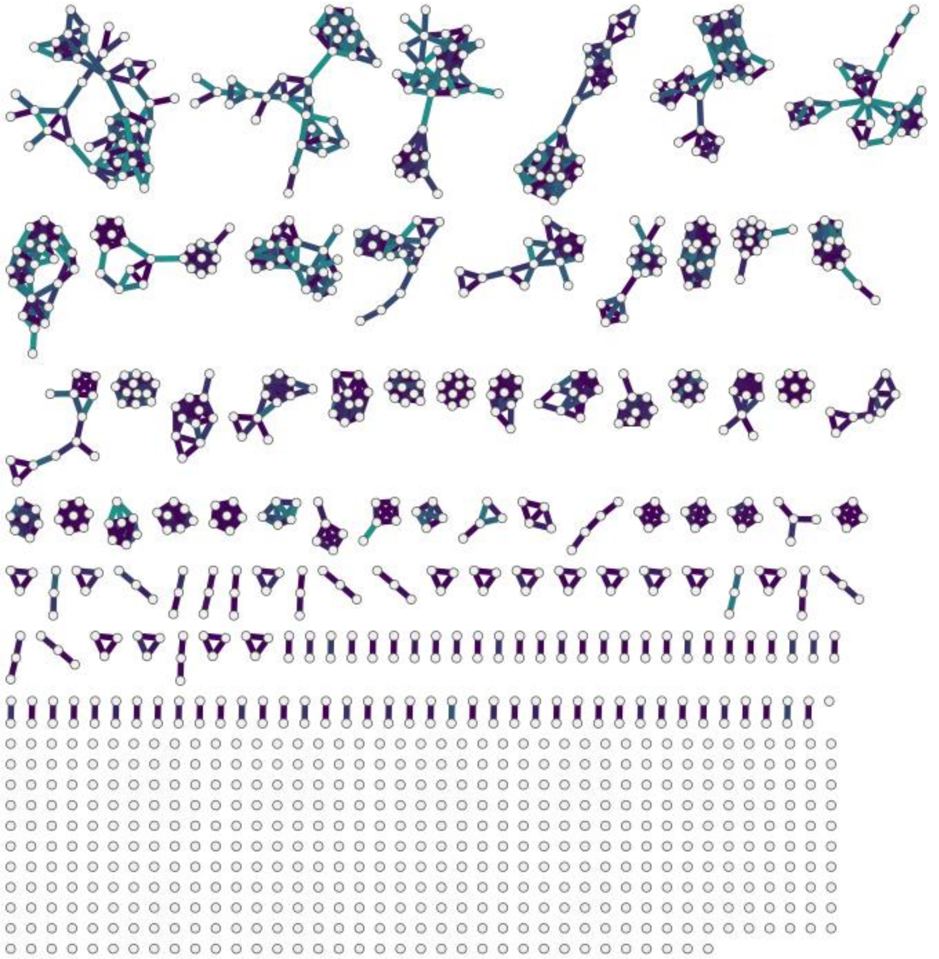

**Fig. S42.** Threshold molecular network of the NIH Natural Products Library obtained after applying SpecReBoot to the MS2DeepScore–based molecular network. Edges shown correspond to spectrally similar connections retained after applying both the spectral similarity threshold (≥ 0.8) and a bootstrap-derived edge support filter (≥ 0.5), resulting in a reduced network enriched in stable and chemically coherent relationships.

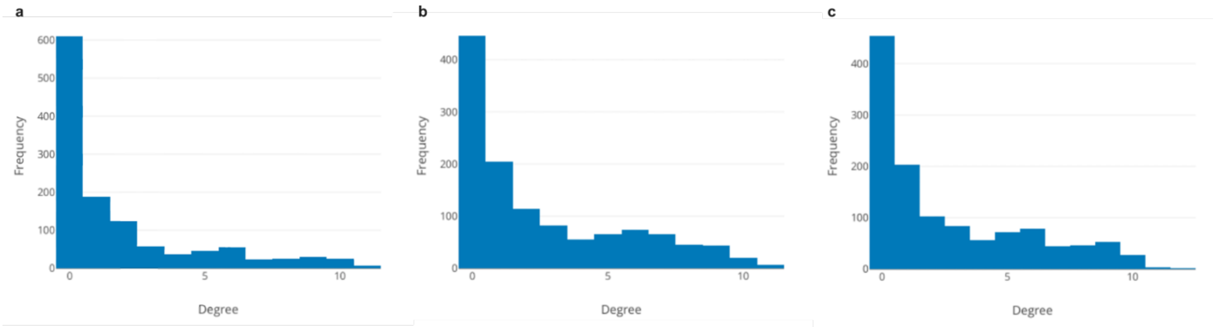

**Fig. S43.** Histograms show the node degree distributions for molecular networks built using the SpecReBoot dual-filter strategy based on (**a**) modified cosine, (**b**) MS2DeepScore, and (**c**) Spec2Vec for the NIH Natural Products Library. All networks are characterized by a predominance of low-degree nodes and a limited number of moderately connected molecular families, indicating comparable global organization across similarity metrics despite metric-specific differences in local connectivity.

**Table S3.** Global topological properties reported for Spec2Vec, MS2DeepScore, and modified cosine–based molecular networks of the NIH Natural Products Library after applying the SpecReBoot dual-filter strategy. All networks exhibit comparable global organization, including similar numbers of connected components, average degree ranges, and centralization values, indicating convergence toward a shared large-scale topology.

| Similarity metric | Singletons | Connected components | Avg degree | Density | Clustering | Centralization |
|---|---|---|---|---|---|---|
| Spec2Vec | 610 | 140 | 4.95 | 0.248 | 0.741 | 0.279 |
| MS2DeepScore | 446 | 141 | 4 | 0.098 | 0.372 | 0.128 |
| ModCosine | 454 | 147 | 3.32 | 0.09 | 0.367 | 0.134 |

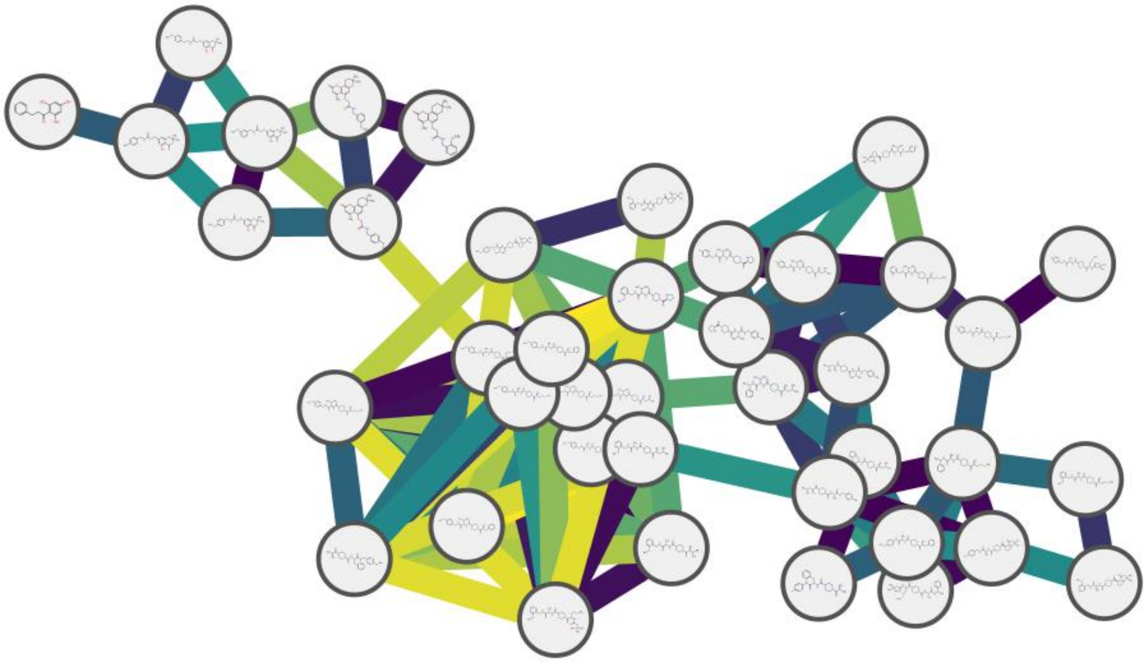

**Fig. S44.** Molecular family containing spectra *i* and *j* in the base molecular network of the NIH Natural Products Library constructed using the modified cosine similarity score. The network was generated using a spectral similarity threshold of 0.7 and standard topology-based parameters, without incorporating bootstrap-derived edge support (SpecReBoot). Nodes represent MS/MS spectra and edges represent pairwise spectral similarities, colored by similarity magnitude.

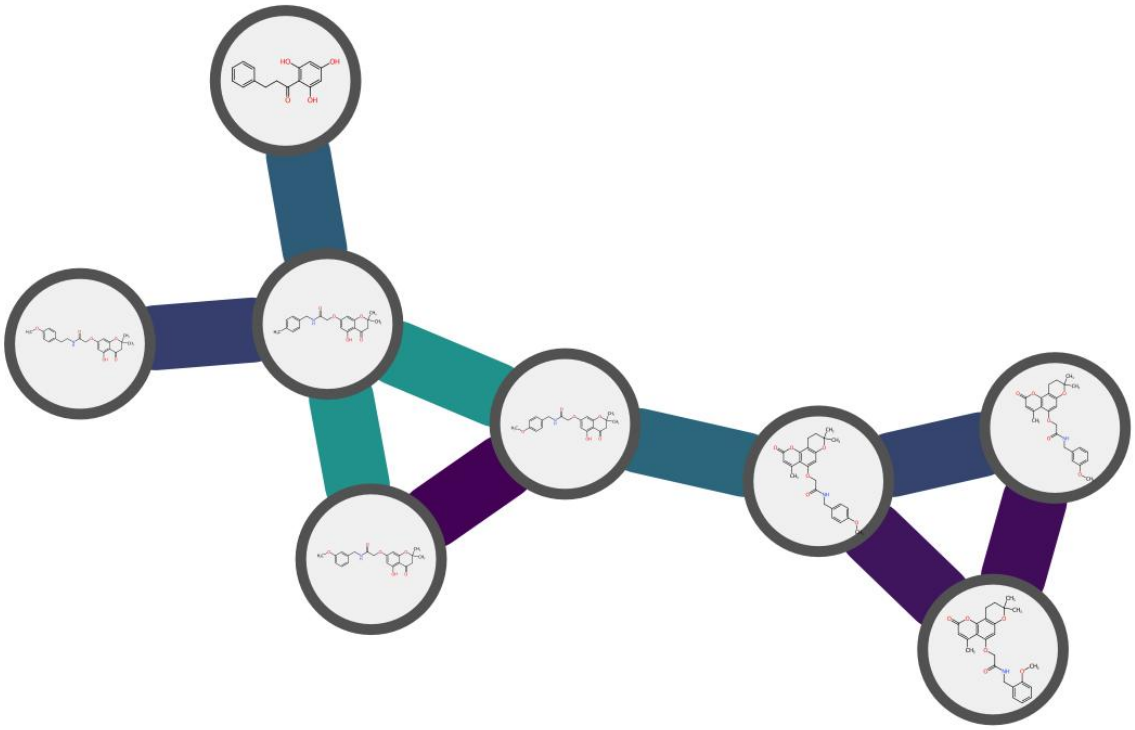

**Fig. S45.** Core molecular family for hybrid phenylpropanoids in the NIH Natural Products Library constructed using the modified cosine similarity score with SpecReBoot. Only high-confidence core edges retained after applying the bootstrap-derived edge support threshold are shown. Nodes represent MS/MS spectra and edges are colored according to edge support.

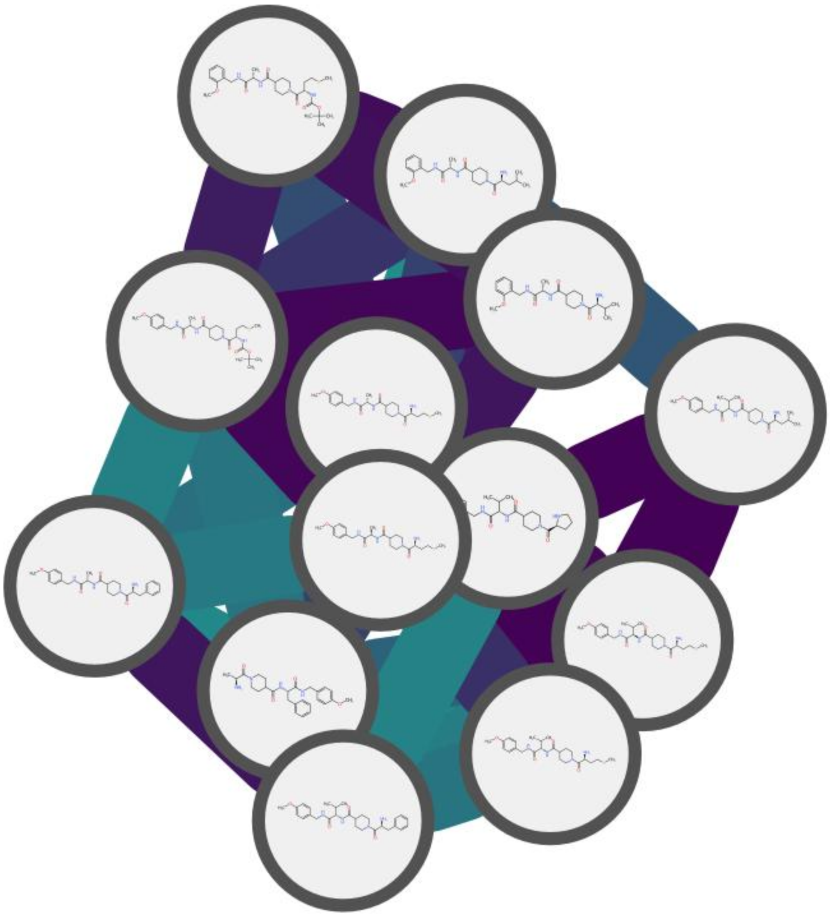

**Fig. S46.** Core molecular family for linear tetrapeptides in the NIH Natural Products Library constructed using the modified cosine similarity score with SpecReBoot. Only high-confidence core edges retained after applying the bootstrap-derived edge support threshold are shown.

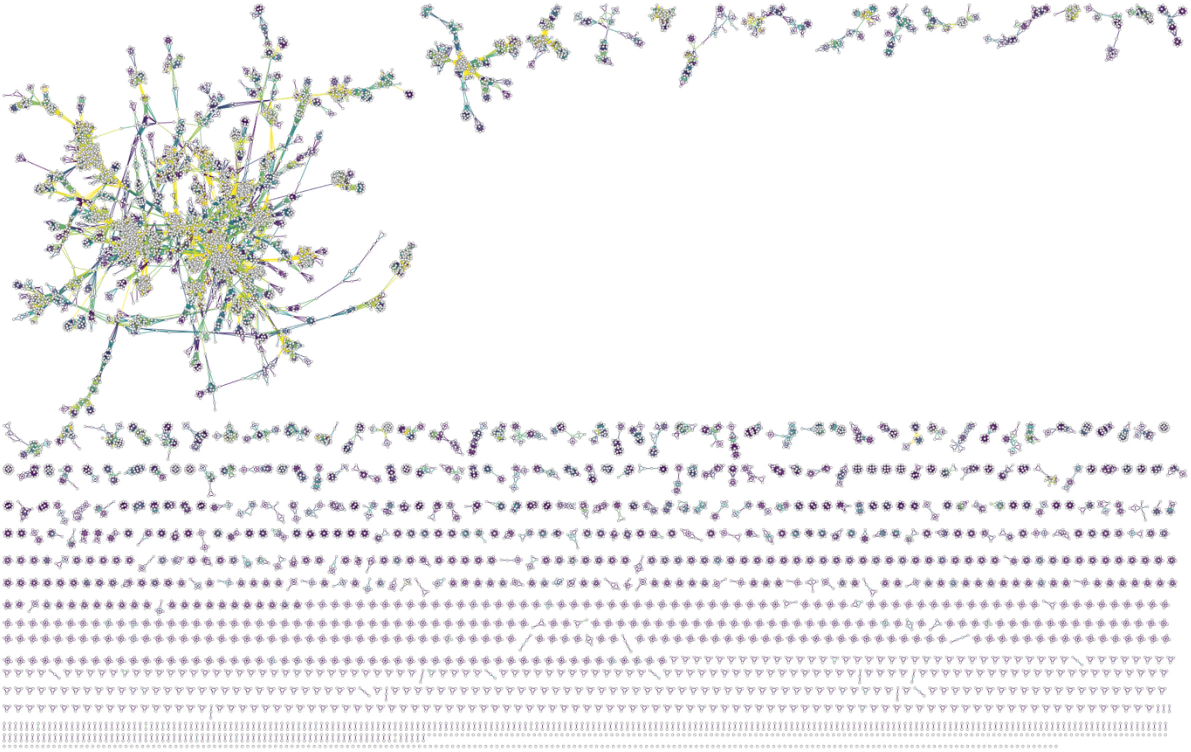

**Fig. S47.** Base molecular network of the the MSn-COCONUT dataset constructed using the flash cosine similarity score. The network was generated using a spectral similarity threshold of 0.7, without incorporating bootstrap-derived edge support. Nodes represent MS/MS spectra, and edges represent pairwise spectral similarities.

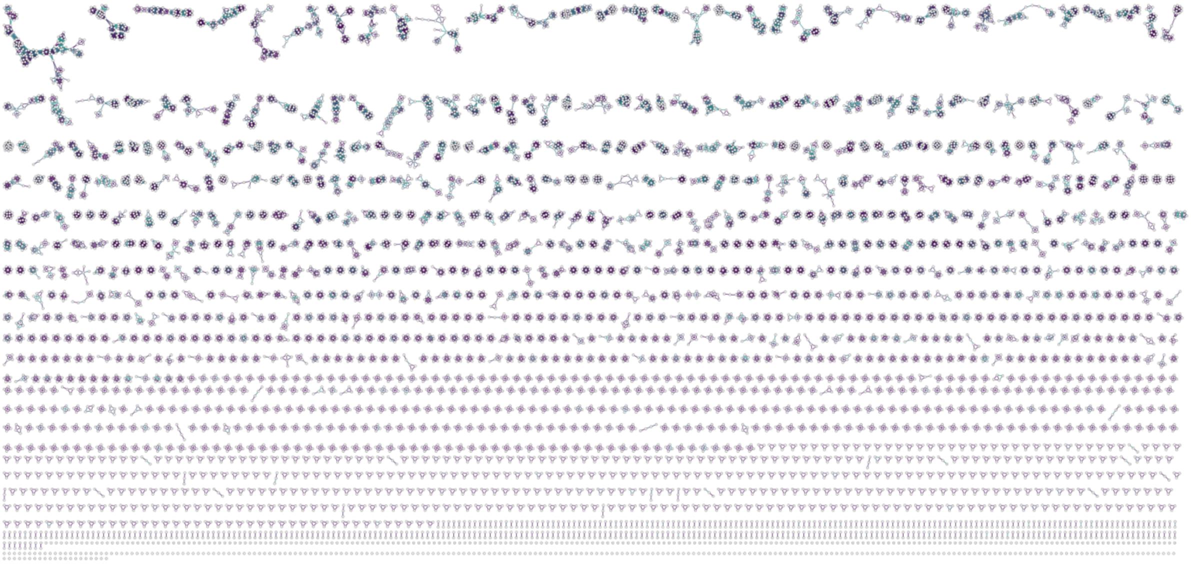

**Fig. S48.** Threshold molecular network of the MSn-COCONUT dataset obtained after applying SpecReBoot to the flash cosine–based molecular network. Edges shown correspond to spectrally similar connections retained after applying both the spectral similarity threshold (≥ 0.7) and a bootstrap-derived edge support filter (≥ 0.5), resulting in a reduced network enriched in stable and chemically coherent relationships.

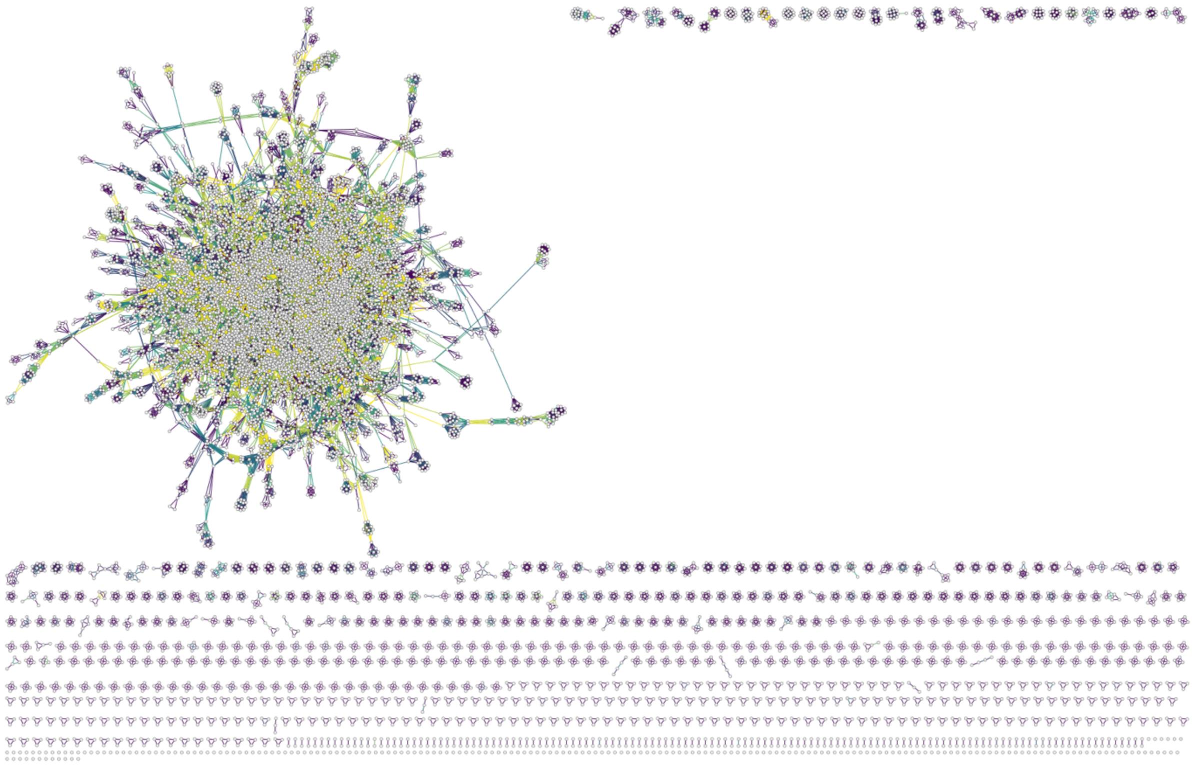

**Fig. S49.** Base molecular network of the the MSn-COCONUT dataset constructed using the flash modified cosine similarity score. The network was generated using a spectral similarity threshold of 0.7, without incorporating bootstrap-derived edge support. Nodes represent MS/MS spectra, and edges represent pairwise spectral similarities.

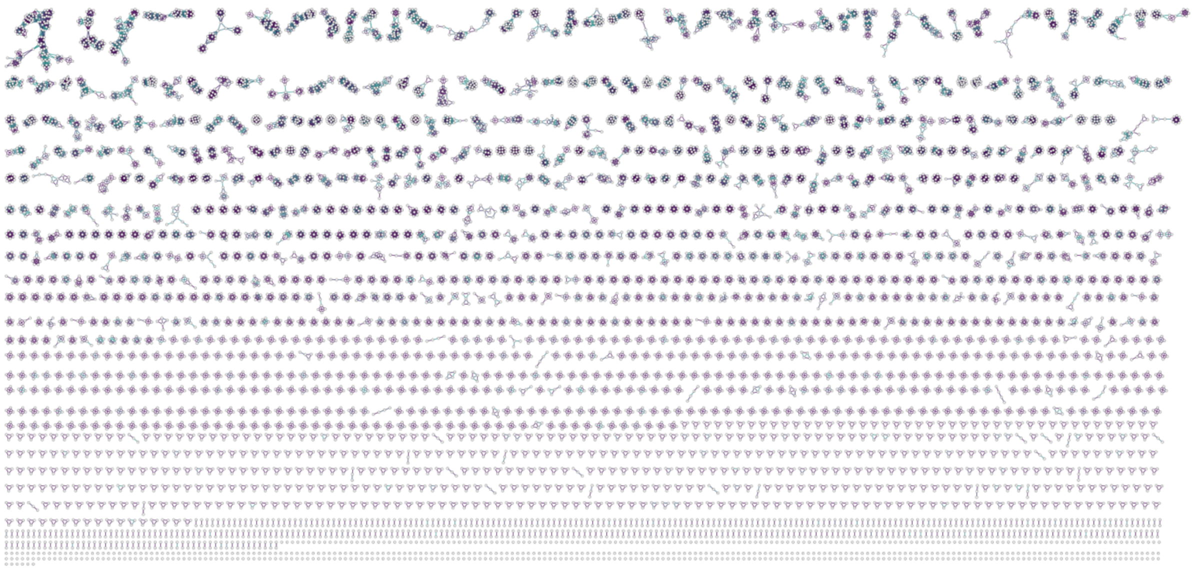

**Fig. S50.** Threshold molecular network of the MSn-COCONUT dataset obtained after applying SpecReBoot to the flash modified cosine–based molecular network. Edges shown correspond to spectrally similar connections retained after applying both the spectral similarity threshold (≥ 0.7) and a bootstrap-derived edge support filter (≥ 0.5), resulting in a reduced network enriched in stable and chemically coherent relationships.

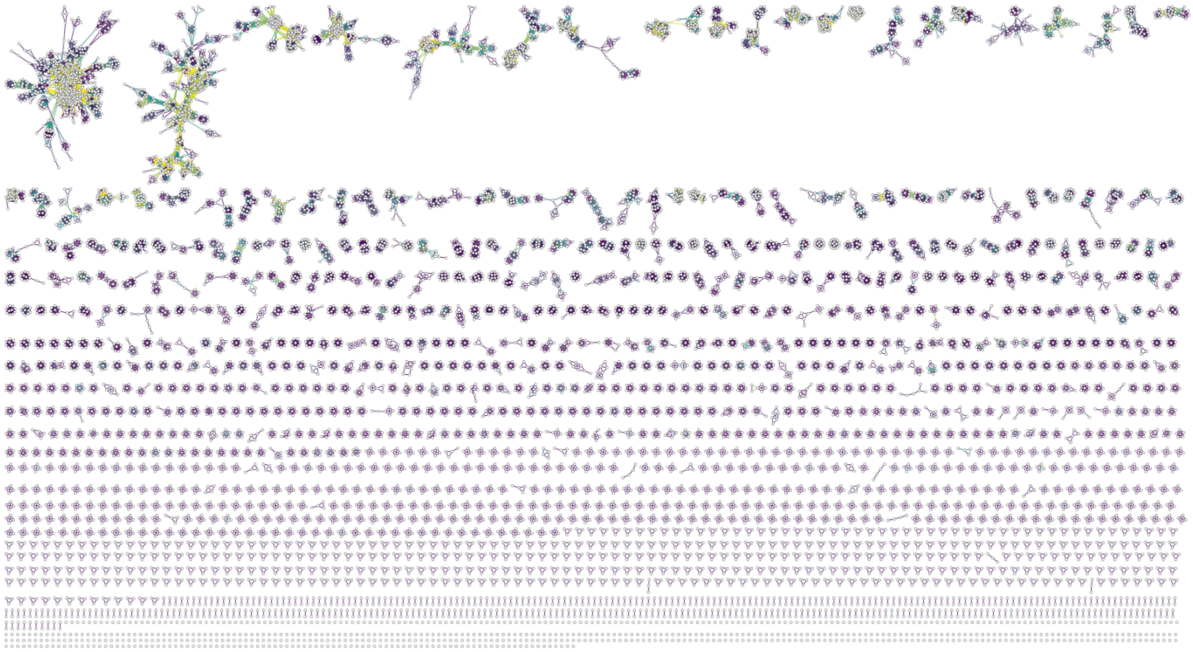

**Fig. S51.** Base molecular network of the the MSn-COCONUT dataset constructed using the Spec2Vec similarity score. The network was generated using a spectral similarity threshold of 0.7, without incorporating bootstrap-derived edge support. Nodes represent MS/MS spectra, and edges represent pairwise spectral similarities.

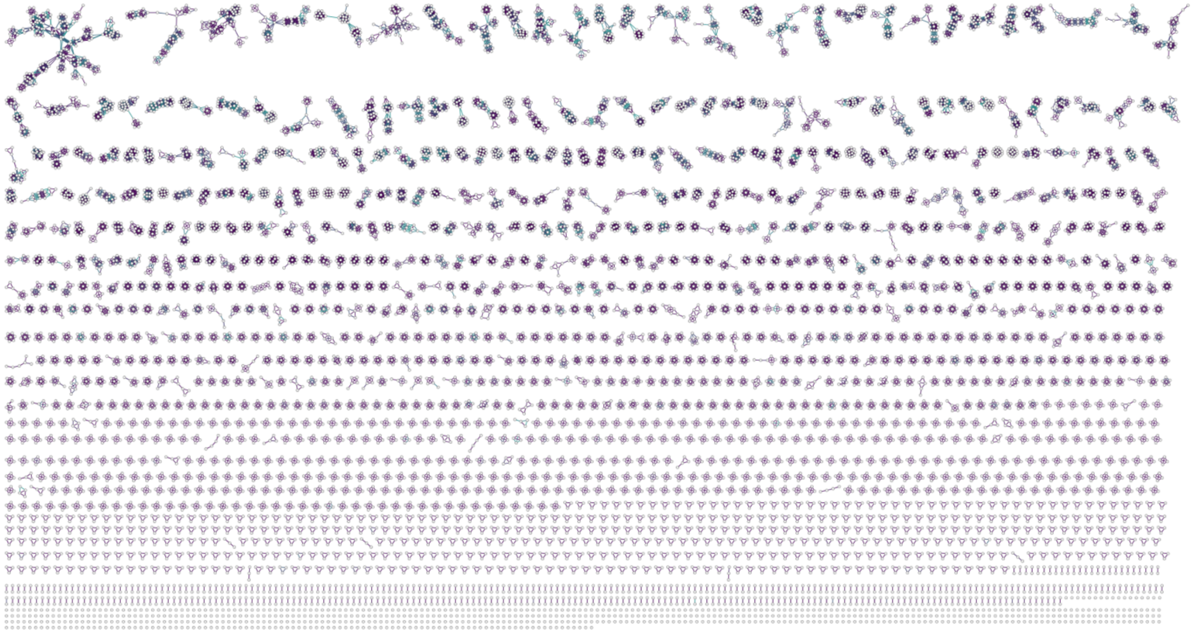

**Fig. S52.** Threshold molecular network of the MSn-COCONUT dataset obtained after applying SpecReBoot to the Spec2Vec–based molecular network. Edges shown correspond to spectrally similar connections retained after applying both the spectral similarity threshold (≥ 0.7) and a bootstrap-derived edge support filter (≥ 0.5), resulting in a reduced network enriched in stable and chemically coherent relationships.

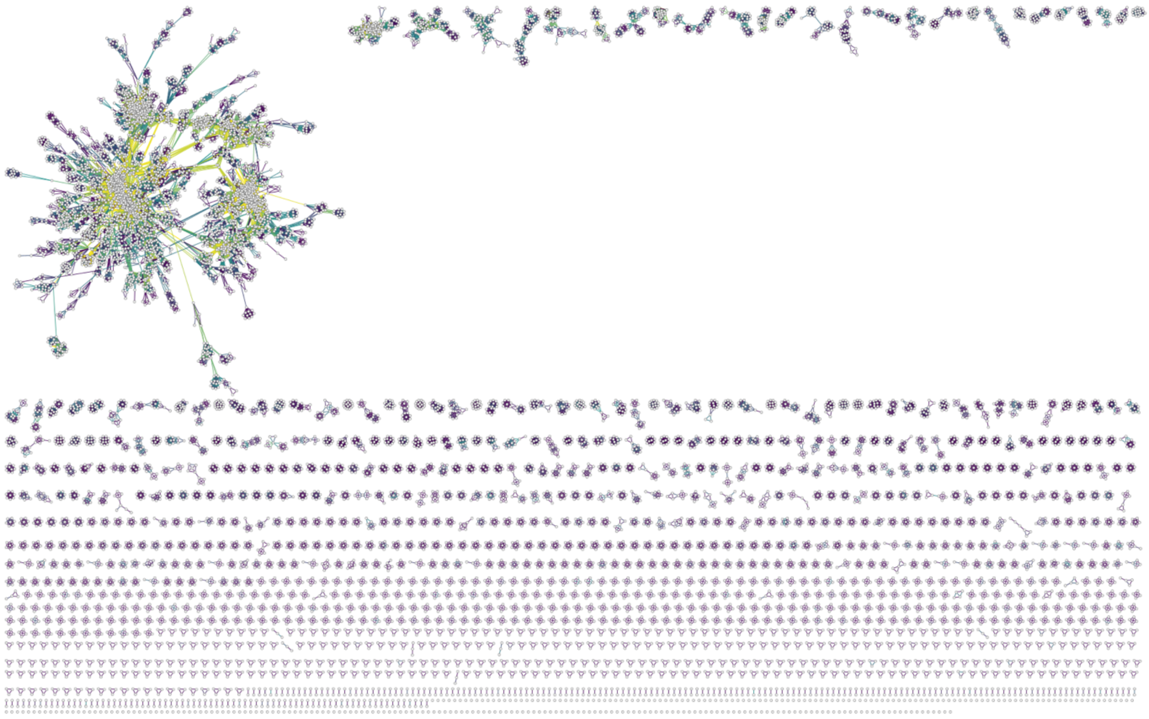

**Fig. S53.** Base molecular network of the the MSn-COCONUT dataset constructed using the MS2DeepScore similarity score. The network was generated using a spectral similarity threshold of 0.7, without incorporating bootstrap-derived edge support. Nodes represent MS/MS spectra, and edges represent pairwise spectral similarities.

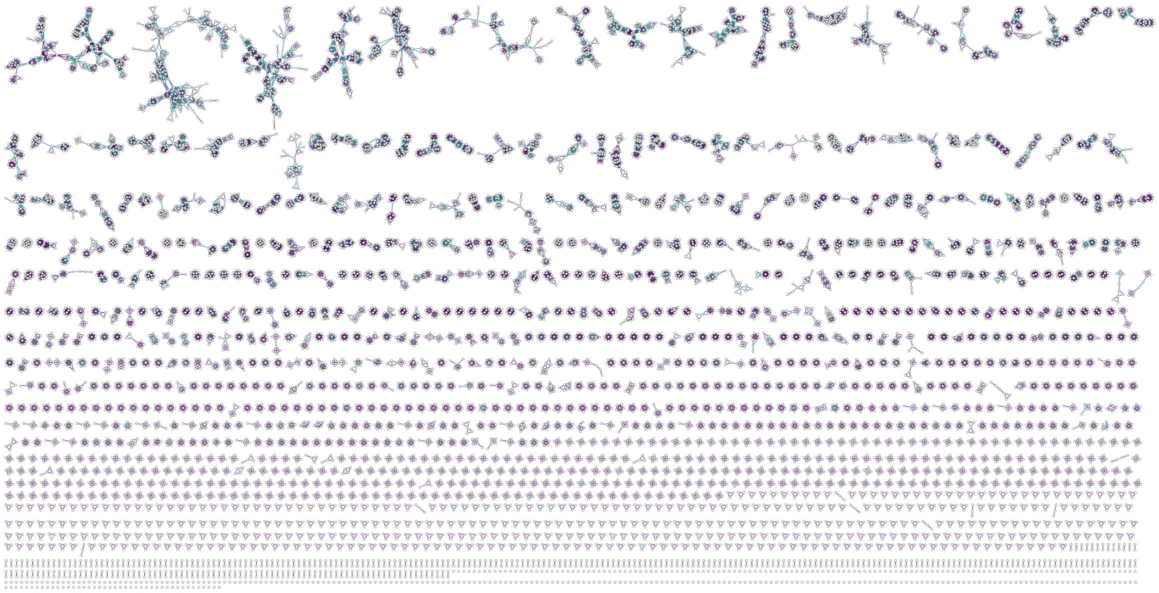

**Fig. S54.** Threshold molecular network of the MSn-COCONUT dataset obtained after applying SpecReBoot to the MS2DeepScore–based molecular network. Edges shown correspond to spectrally similar connections retained after applying both the spectral similarity threshold (≥ 0.7) and a bootstrap-derived edge support filter (≥ 0.5), resulting in a reduced network enriched in stable and chemically coherent relationships.

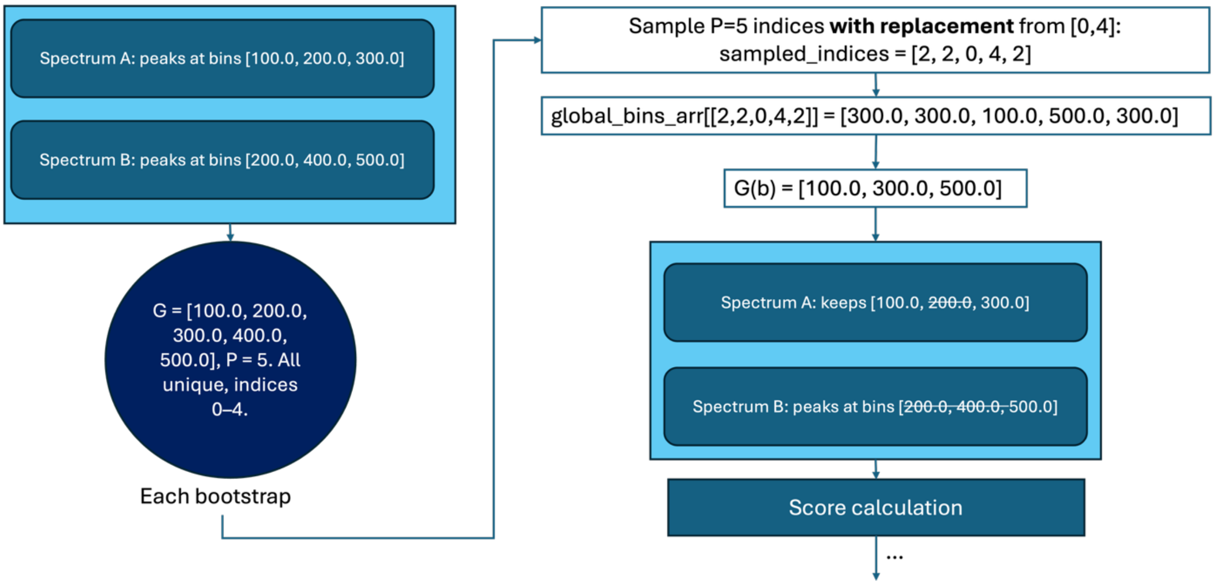

**Fig. S55.** Schematic overview of the SpecReBoot bootstrap resampling strategy. For each bootstrap replicate, a global bin space G is constructed from all unique *m/z* bins observed across the input spectra (here P=5 bins: [100.0, 200.0, 300.0, 400.0, 500.0]). P indices are sampled with replacement from G, and the unique values of the resulting sample define the bootstrap bin set G(b). Each spectrum is then masked to retain only peaks falling within G(b), while bins absent from G(b) are removed. The masked spectra are passed to the similarity scoring function, and this process is repeated across all bootstrap replicates.
